# Heteromorphic tertiary structures of MAT1-1-1 and MAT1-2-1 protein variants simultaneously produced in pairs by *Ophiocordyceps sinensis* strains

**DOI:** 10.64898/2026.02.16.706022

**Authors:** Xiu-Zhang Li, Yu-Ling Li, Wei Liu, Jian-Zhao Qi, Jia-Shi Zhu

## Abstract

The MATα_HMGbox domain in the MAT1-1-1 protein and the HMG-box_ROX1-like domain in the MAT1-2-1 protein play essential roles in DNA binding and regulating the transcription of genes that control sexual reproduction in *Ophiocordyceps sinensis*. Previous studies have reported differential occurrences, differential transcription, and alternative splicing of the *MAT1-1-1*, *MAT1-2-1*, and pheromone receptor genes in *Hirsutella sinensis* (Genotype #1 among 17 genome-independent genotypes of *O. sinensis* fungi). In this study, we further demonstrated that the DNA-binding domains of the paired MAT1-1-1 and MAT1-2-1 proteins derived from each of the 20 purportedly “pure” *O. sinensis* strains exhibit heteromorphic tertiary structures, as predicted by AlphaFold 3D structural modeling. These differentially paired mating proteins are characterized by different truncations, diverse 1−4 amino acid substitutions at distinct sites, altered hydrophobicity and secondary structures, and heteromorphic tertiary structures, indicating divergency in the fungal origins of the mating proteins within the mycologically and genetically impure *O. sinensis* strains. Our findings support the self-sterility hypothesis for *O. sinensis* and likely ensure the fidelity and genetic diversity of heterothallic or hybrid reproduction during the lifecycle of the *Cordyceps sinensis* insect–fungal complex.

## 1. Introduction

Natural *Cordyceps sinensis*, also known as Chinese cordyceps (non-Latin term) or DōngChóng XiàCǎo 冬虫夏草 in Chinese, exists only in high-altitude alpine meadows (3,000−5,000 m) on the Qinghai-Tibet Plateau, which contributes to its natural scarcity. It has a centuries-long history of clinical application in health maintenance, characterized by the dual reinforcement of “*YIN* 阴” and “*YANG* 阳”, as well as in disease amelioration, postillness and postoperative recovery, and antiaging therapy [1–4]. Modern science is increasingly elucidating the complex biochemical mechanisms (*e.g.*, metabolic (glucose and lipid) regulation, organ protection, immunomodulation, anti-inflammatory activity, antioxidant activity, hormone-like activity supporting libido and fertility, and the enhancement of energy–vitality–stamina, etc.) that underpin these traditional applications, proving its status as a valuable therapeutic resource, despite the complexity and rarity of this medicinal resource. This natural substance, with its unique therapeutic significance, embodies a fusion of nature’s ingenuity and traditional medicinal wisdom; however, its high value has led to overharvesting, making it a Level-II protected natural substance [5].

Natural *C. sinensis* comprises the fruiting body of *Ophiocordyceps sinensis* and the remains of mummified *Hepialidae* moth larva, which retains a thick body wall with numerous bristles, an intact larval intestine, head tissues, and fragments of other larval tissues [6–13]. Dual-fluorescence microscopy has revealed multicellular heterokaryotic microstructures of *C. sinensis* ascospores and hyphae, containing mono-, bi-, tri-, and tetra-nucleate cells [14]. Natural *C. sinensis* exhibits apparent genetic heterogeneity, harboring >90 fungal species from at least 37 fungal genera, as well as 17 genomically independent genotypes of *O. sinensis* that differentially naturally co-occur, the abundance of which undergoes dynamic and reciprocal alterations in different compartments of *C. sinensis* during maturation, supporting their genome-independent natural characteristics [9,10,12,13,15–44]. Thus, the Chinese Pharmacopoeia defines this medicinal microecosystem as an insect‒fungal complex.

Among the numerous intrinsic co-occurring fungal species [20,31–33,40,41], *Hirsutella sinensis* has been postulated to be the only anamorph of *O. sinensis* [45], although this hypothetical claim needs to be validated based on all 4 criteria of Koch’s postulates. In terms of validation, Wei *et al*. [46] reported a successful artificial cultivation project conducted in an industrial product-oriented setting. Unfortunately, they reported a species contradiction that arose between the anamorphic inoculants of 3 *H. sinensis* strains (GC-biased Genotype #1 of *O. sinensis*) inoculated on *Hepialidae* moth larvae and the genome-independent, sole teleomorphic AT-biased Genotype #4 of *O. sinensis* identified in the fruiting body of cultivated *C. sinensis*.

Since the 1840s, the Latin name *Cordyceps sinensis* has been indiscriminately applied to both the teleomorph/holomorph of the fungus *C. sinensis* and the wild insect‒fungal complex, disregarding Edward Doubleday’s original identification in 1842 that the insect portion of the natural material belongs to Agrotis [47]. This imprecise use of the Latin name has blurred the distinction between the fungus, its host, and the insect−fungal complex, resulting in ambiguity in the scientific literature, commercial trade in the mass market, and governmental regulations worldwide. The fungus was renamed *O. sinensis* in 2007, and *H. sinensis* strain EFCC7287 (GC-biased Genotype #1 of *O. sinensis*) was used as the nomenclature reference [9,10,48–50]. Zhang *et al*. [51] improperly implemented the “One Fungus=One Name” nomenclature rule set by the International Mycological Association (IMA) and replaced the anamorphic name *H. sinensis* with the teleomorphic name *O. sinensis* according to the international rule [52,53]. They interpreted *O. sinensis* as a single fungus to fulfill the prerequisite of “One Fungus” of the international rule but did not fully consider the 17 genomically independent genotypes of *O. sinensis* fungi, which differentially coexist in the different compartments of the *C. sinensis* insect–fungal complex. In the present paper, we retain the anamorphic name *H. sinensis* for GC-biased Genotype #1 among 17 genotypes of *O. sinensis* and refer to the genomically independent Genotypes #2‒17 fungi as *O. sinensis* on the basis of the taxonomic annotations used in several public depositories, including GenBank and AlphaFold, pending the formal determination and differentiation of the systematic taxonomic positions of these GC- and AT-biased Genotypes #2–17. We also maintain the long-standing use of the traditional name *C. sinensis* for both wild and cultivated insect‒fungal complexes because the 2007 renaming of *C. sinensis* to *O. sinensis*, which is based on *H. sinensis* strain EFCC7287 as the nomenclatural reference, did not address the long-standing, indiscriminate application of the Latin name to the insect‒fungal complex [9,10,48,51,54]. Nevertheless, this imperfect and temporarily necessary practice will likely be revised in the future through the differential use of proprietary and exclusive Latin names for the multiple genome-independent genotypes of *O. sinensis* fungi and the insect‒fungal complex as a unique medicinal entity.

The sexual reproductive behavior of ascomycetes is controlled by transcription factors encoded at the mating-type (*MAT*) locus, which constitute the key regulatory mechanism governing mating type recognition and compatibility and the development of fruiting bodies and sexual reproductive structures and cells such as ascocarps and ascospores [55–62]. The reproductive behavior of *O. sinensis* depends on the synergistic interplay of 2 core mating proteins, *MAT1-1* and *MAT1-2* idiomorphs. The MAT1-1-1 protein harbors a mating-type alpha high mobility group box (MATα_HMGbox) domain, whereas the MAT1-2-1 protein contains the high mobility group box ROX1-like (HMG-box_ROX1-like) domain [14,58,63–65]. These 2 functional domains of mating proteins play essential roles in regulating the expression of genes associated with sexual reproduction. Li *et al*. [65–68] documented the differential occurrence, differential transcription, and alternative splicing of mating-type and pheromone receptor genes in the genomes of *H. sinensis* strains, challenging the prior self-fertilization hypothesis for *H. sinensis via* homothallism or pseudohomothallism. Instead, these discoveries support self-sterility consistent with a heterothallic or hybrid lifestyle, indicating the necessity of compatible sexual partners to complete sexual reproduction within the *C. sinensis* insect‒fungus complex. Moreover, Li *et al*. [65,69] reported the heteromorphic tertiary structures of the DNA-binding domains of the full-length MAT1-1- 1 and MAT1-2-1 proteins in wild-type *C. sinensis* isolates, which might be generated by either GC-biased Genotype #1 (*H. sinensis*) or Genotype #3 of *O. sinensis*, further underscoring the complex heterothallic or hybrid sexual reproduction of *O. sinensis*.

Variations in the DNA-binding domains of the 2 mating proteins, arising from diverse amino acid substitutions, induce changes in their tertiary structures and synergistic interactions between the 2 idiomorphic mating proteins. These proteins are derived from numerous mycologically impure, wild-type *C. sinensis* isolates [65], which were characterized by Prof. Zhang Y-J and his associates [7,51,70–72]. The present study further analyzed differential N- and/or C-terminal truncations and diverse amino acid substitutions of the mating proteins derived from 20 purportedly “pure” *O. sinensis* strains reported by Prof. Yao Y-J and his collaborators [14,36] to clarify the protein structural basis of *O. sinensis* sexual reproduction. The changes in the primary, secondary, and tertiary structures of the MATα_HMGbox and the HMG-box_ROX1-like domains of the MAT1-1-1 and MAT1-2-1 proteins were correlated. Critically, the present study focused on the differential pairings of MAT1-1-1 and MAT1-2-1 protein variants generated simultaneously by 20 *O. sinensis* strains to explore their possible heterospecific fungal origins within these heterogeneous but purportedly “pure” *O. sinensis* strains.

## 2. Materials and Methods

### 2.1 MAT1-1-1 and MAT1-2-1 protein sequences from *O. sinensis* strains

The AlphaFold database provides AI-predicted 3D structural morphs for 20 truncated MAT1-1-1 proteins (AGW27517−AGW27536) and 5 truncated MAT1-2-1 proteins (AGW27543, AGW27548, AGW27552, AGW27554, and AGW27555) derived from 20 *O. sinensis* strains [14]. Li *et al*. [36] reported that 11 of the 20 strains were obtained from fresh caterpillar body samples collected from various *C. sinensis* production regions on the Qinghai‒Tibet Plateau, marked as “TS” in Table 1. The remaining 9 *O. sinensis* strains were from the cultures of monoascospores of *C. sinensis* insect‒fungal complexes that were collected from a single production region (Maqên, Guoluo, Qinghai Provence of China), which are marked as “SS” in Table 1.

**Table 1.**
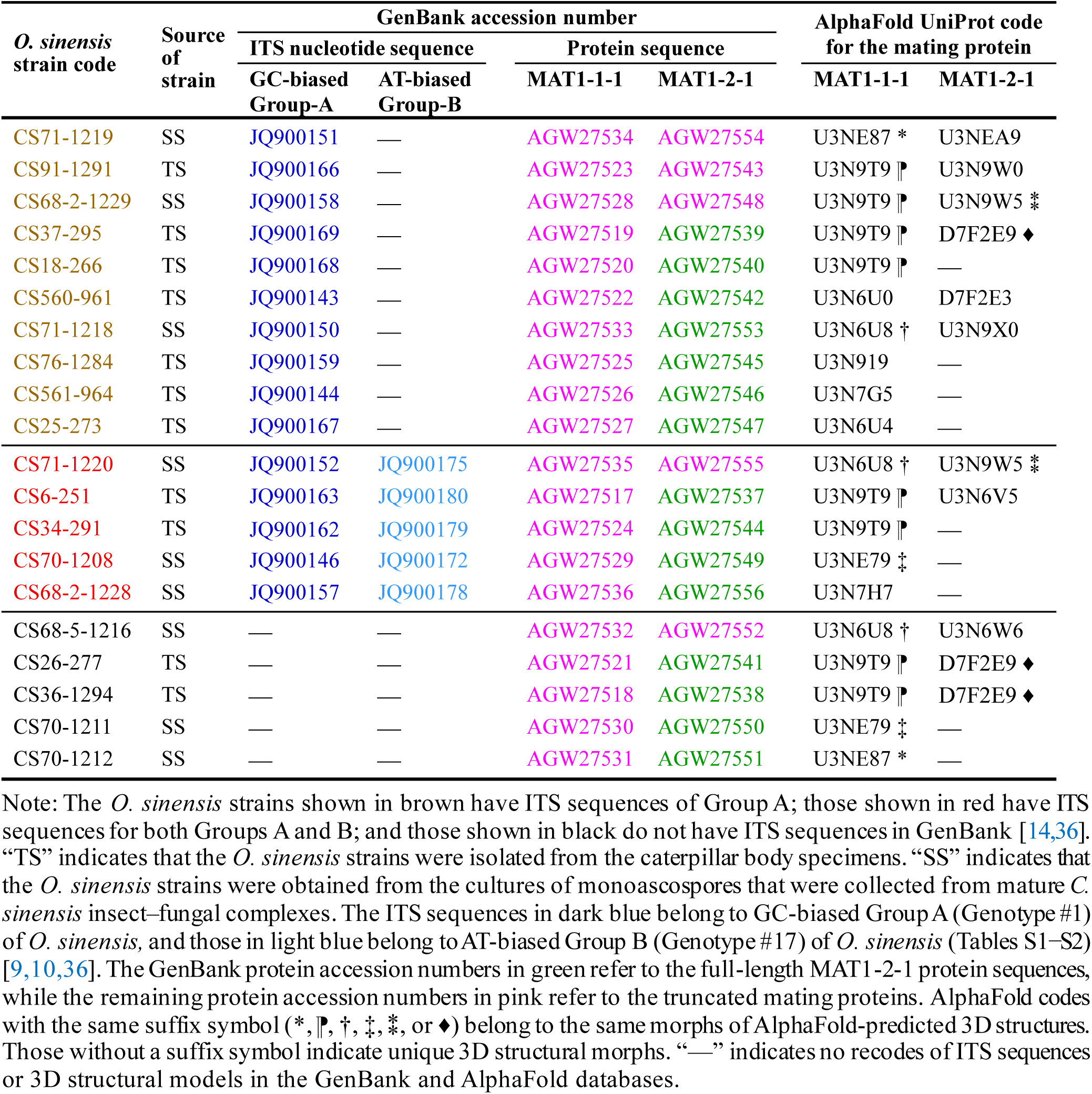
*O. sinensis* strains and their tissue sources [36], GenBank accession numbers for the internal transcribed spacer (ITS) nucleic acid sequences and mating proteins, and AlphaFold UniProt codes for the truncated and full-length MAT1-1-1 and MAT1-2-1 proteins derived from the *O. sinensis* strains.

The MAT1-1-1 protein sequences of different lengths were encoded by the *MAT1-1-1* cDNAs that were amplified from total RNA extracted from the *O. sinensis* strains using the primer pair *m1F3/m1R3*, while the *MAT1-2-1* cDNAs were amplified using the primer pair *Mat1-2F*/*Mat1-2R* (Figures S1−S2) [14]. Li *et al*. [36] deposited the internal transcribed spacer (ITS) sequences into GenBank for 15 of the 20 *O. sinensis* strains, including 10 homogeneous strains shown in brown in Table 1 that reportedly harbor only the GC-biased Group-A ITS sequences and 5 other strains shown in red in Table 1 that were characterized as heterogeneous, containing both GC-biased Group-A and AT-biased Group-B ITS sequences. The remaining 5 *O. sinensis* strains shown in black in Table 1 lack ITS sequence information in GenBank.

### 2.2 Alignment of the DNA-binding domain sequences of the mating proteins

The amino acid sequences of the MATα_HMGbox domains of the MAT1-1-1 proteins and the HMG-box_ROX1-like domains of the MAT1-2-1 proteins from the *O. sinensis* strains were aligned using the GenBank Blastp program (https://blast.ncbi.nlm.nih.gov/ (Bethesda, MD, USA), accessed from 18 June 2025 to 31 January 2026).

### 2.3 Amino acid properties and scale analysis

The amino acid sequences of the DNA-binding domains of the mating proteins were scaled based on the general properties of their side chains (https://web.expasy.org/protscale/; Basel, Switzerland, accessed from 18 May 2025 to 20 October 2025) (Table 3) [65,68,69,73–77]. The MATα_HMGbox domain of MAT1-1 proteins (amino acids 51→225 of the reference sequence AGW27560 derived from the *H. sinensis* strain CS68-2-1229 [14]) and the HMG-box_ROX1-like domain of MAT1-2-1 proteins (amino acids 127→197 of the reference sequence AEH27625 [70]), with additional 9 amino acid residues extending upstream and downstream of the domains to ensure comprehensive coverage, were plotted sequentially with a window size of 9 amino acid residues using the linear weight variation model of the ExPASy ProtScale algorithms [65,68,69,73–75] to generate ExPASy ProtScale plots and predict the potential changes in the hydrophobic properties and secondary structures of proteins related to α-helices, β-sheets, β-turns, and coils. The resulting topological structures and waveforms shown in the ProtScale plots were compared to exploring changes in the hydrophobicity and 2D structural features of the DNA-binding domains of the mating proteins.

### 2.4 AlphaFold-based prediction of the tertiary structures of the mating proteins

To analyze the heteromorphic stereostructures in this study, the 3D structures of the MAT1-1-1 and MAT1-2-1 proteins from *O. sinensis* strains were predicted computationally from their amino acid sequences utilizing artificial intelligence (AI)-driven machine learning technology AlphaFold (https://alphafold.com/, Cambridgeshire, UK; accessed from 18 June 2025 to 20 October 2025) [78−88].

The AlphaFold database provides per-residue model confidence, predicts scores between 0 and 100 using the partial local distance difference test (pLDDT), and assigns a per-residue score to each individual residue [78−81,83–85]. A color-coding scheme based on the pLDDT score is used for the 3D structure to visually represent the confidence levels: residues with very high confidence (pLDDT>90) are shown in dark blue, those with high confidence (90>pLDDT>70) appear in light blue, residues with low confidence (70>pLDDT>50) are shown in yellow, and residues with very low confidence (pLDDT<50) are shown in orange [65,86,87,89].

The main reasons for the low confidence in the AlphaFold predictions include the following: (1) The supporting data are insufficient, as the AlphaFold-based prediction relies on the quality of the multiple sequence alignment (MSA). If few homologous sequences are available in a certain region of the MSA, the model will lack sufficient evolutionary information to infer the structure, resulting in low confidence. (2) Some protein regions are inherently flexible under physiological conditions, such as the activation domains of transcription factors, which do not have a fixed 3D structure, leading to low confidence in the predicted structure assigned by AlphaFold. (3) In terms of special structural regions, such as small-molecule binding sites and artificial linkers in fused proteins, the AlphaFold system may have limited predictive power, leading to insufficient confidence in the predictions of complex structures. Thus, the low-confidence regions in AlphaFold predictions are informative flags for highlighting the challenges of prediction and should not be interpreted with excessive confidence.

### 2.5 Correlations of the primary, secondary, and tertiary structures of the DNA-binding domains of the paired MAT1-1-1 and MAT1-2-1 proteins

Among diverse AlphaFold 3D structural heteromorphs, the changes in the tertiary structure around the mutation sites within the MATα_HMGbox domains of MAT1-1-1 proteins or the HMG-box_ROX1-like domains of MAT1-2-1 proteins were locally magnified to visualize the subtle 3D variations. Primary sequence variations in the DNA-binding domains of the mating proteins, such as peptide chain truncations and amino acid replacements, were correlated with topological structural and waveform changes in hydrophobicity, α-helices, β-sheets, β-turns, and coils, as presented in the ExPASy ProtScale plots, and with the locally magnified 3D structures around the variation sites. The variant MAT1-1-1 and MAT1-2-1 proteins produced by the *O. sinensis* strains were paired to explore their diverse fungal origins.

## 3. Results

### 3.1 Primary structures of the truncated MAT1-1-1 proteins in *O. sinensis* strains

In the alignment with the sequence of the representative authentic full-length MAT1-1-1 protein AGW27560 derived from the *H. sinensis* strain CS68-2-1229 [14], the 20 MAT1-1-1 proteins were differentially truncated at both the N- and C-termini (Figure 1). These truncated proteins of varying lengths were encoded by the cDNAs amplified using the same pair of primers (the forward primer *m1F3* and the reverse complementary primer *m1R3* shown in Figure S1) from the *MAT1-1-1* genes of *O. sinensis* strains. The MATα_HMGbox domains of the authentic full-length MAT1-1-1 proteins represented by the protein AGW27560 are located at amino acids 51→225 (shown in pink in Figures 1 and S1). The MATα_HMGbox domains of the 20 MAT1-1-1 proteins are truncated at the N-termini and correspond to 9 distinct AlphaFold 3D structural morphs (Figure 2, Table 1).

**Figure 1.**
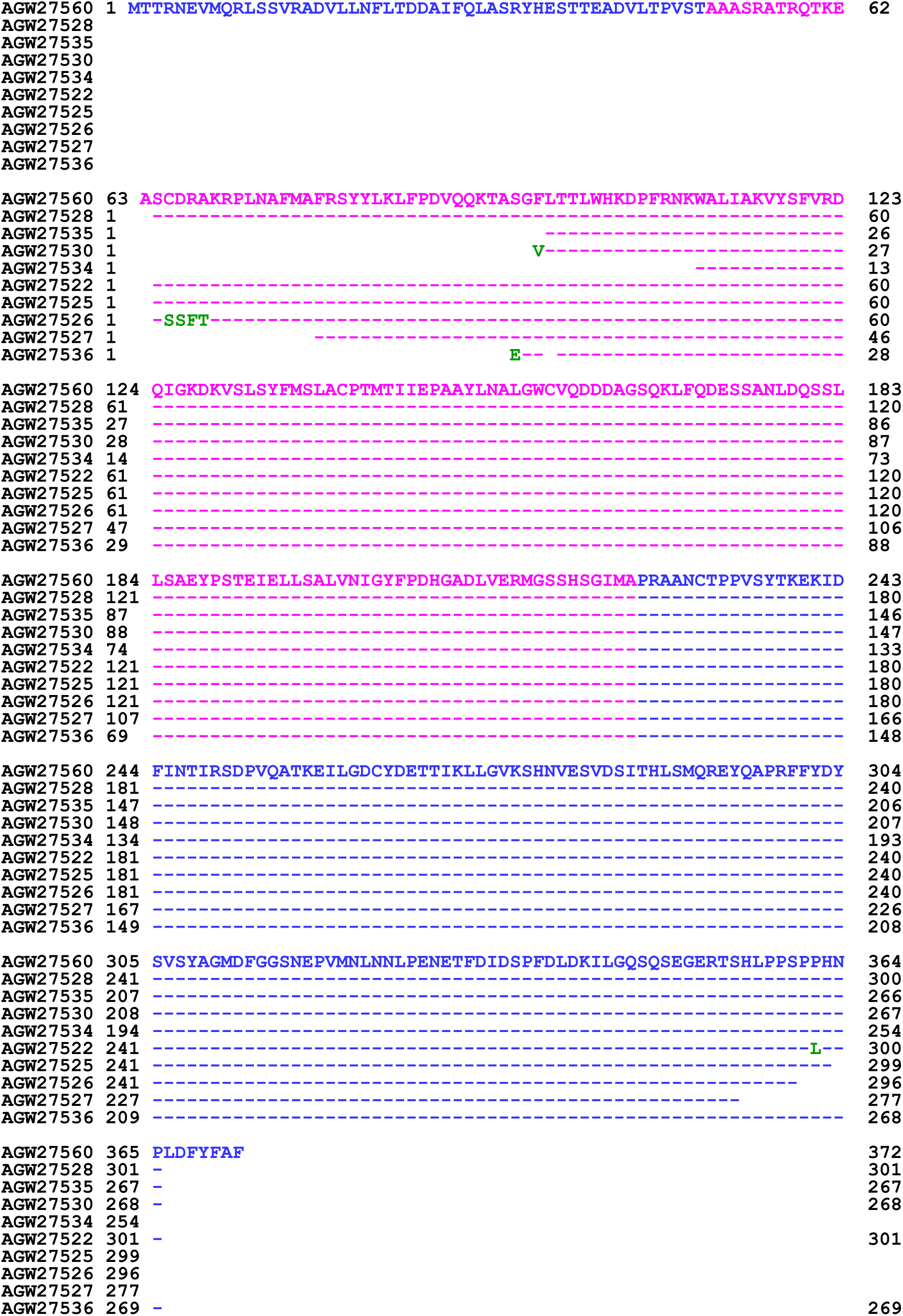
Alignment of the sequences of the query MAT1-1-1 protein AGW27560 derived from *O. sinensis* strain CS68-2-1229 and the 9 truncated MAT1-1-1 proteins (AGW27528, AGW27522, AGW27525, AGW27526, AGW27527, AGW27535, AGW27530, AGW27536, and AGW27534 [14]) that are truncated at both the N- and C-termini. These truncated proteins represent the 20 MAT1-1-1 proteins under 9 AlphaFold 3D structural models (U3N9T9, U3N6U0, U3N919, U3N7G5, U3N6U4, U3N6U8, U3NE79, U3N7H7, and U3NE87). The MATα_HMGbox domain (amino acids 51→225 in the AGW27560 sequence) is shown in pink, and those outside the domain are shown in blue. The amino acid substitutions are shown in green. The hyphens indicate identical amino acids, and the spaces denote unmatched sequence gaps.

**Figure 2.**
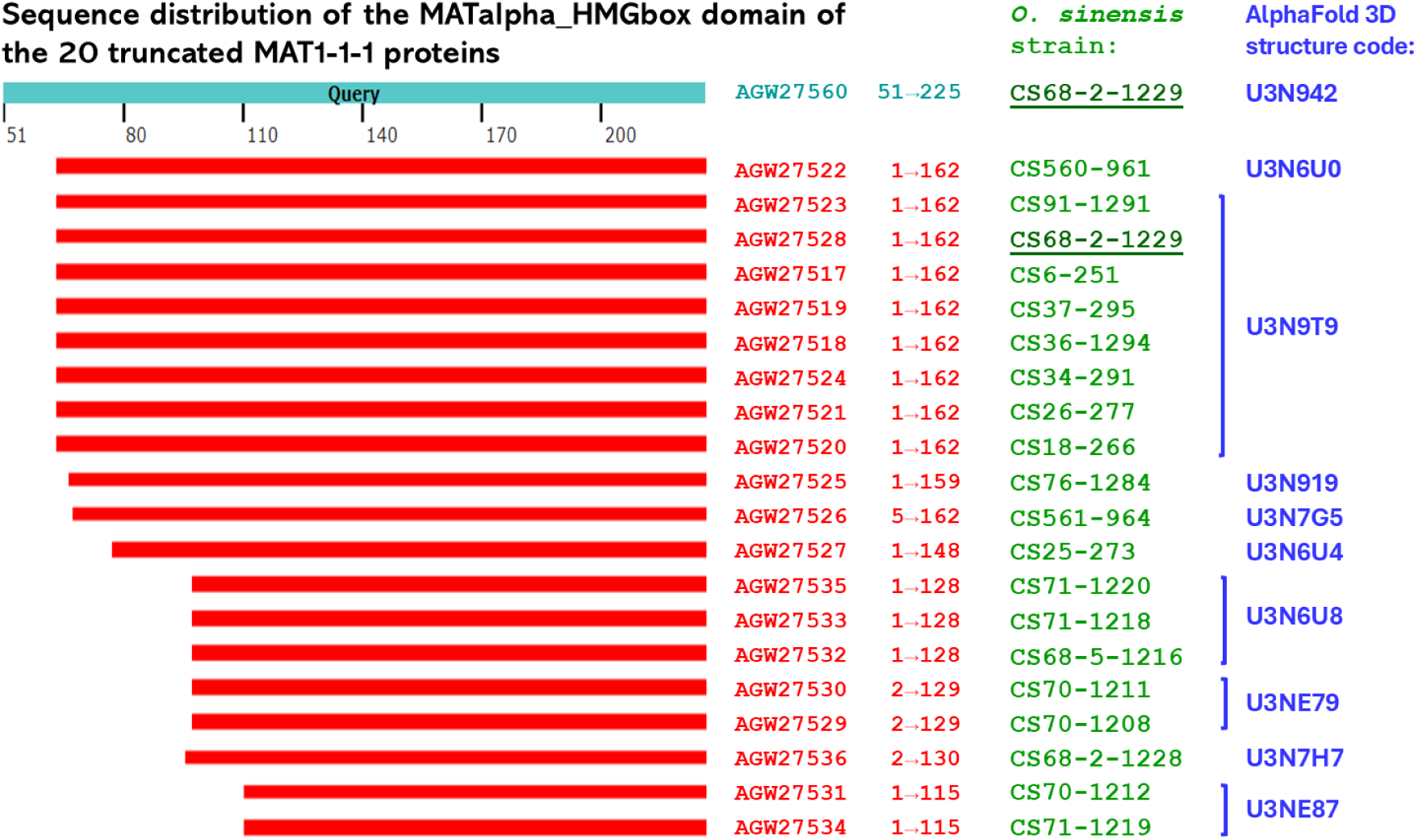
Sequence distribution of the N-terminally truncated MATα_HMGbox domains of 20 MAT1-1-1 proteins derived from *O. sinensis* strains (Table 1). The sequences are aligned with that of the full-length MATα_HMGbox domain (51→225) of the query MAT1-1-1 protein AGW27560 derived from the *H. sinensis* strain CS68-2-1229.

### 3.2 Primary structures of the truncated MAT1-2-1 proteins in *O. sinensis* strains

The AlphaFold database lists 5 MAT1-2-1 proteins produced by *O. sinensis* strains that are truncated differentially at their C-termini (Figure 3). These truncated MAT1-2-1 proteins of varying lengths are encoded by the *MAT1-2-1* cDNAs amplified using the same pair of primers (the forward primer *Mat1-2F* and the reverse complementary primer *Mat1-2R* shown in Figure S2 [14,70]) from the total RNA extracted from various *O. sinensis* strains. The full-length authentic MAT1-2-1 protein AEH27625 [70], under the AlphaFold code U3N942 derived from the *H. sinensis* strain CS2, is used as an alignment reference; the authentic MAT1-2-1 protein contains an HMG-box_ROX1-like domain at amino acids 127→197, as shown in pink in Figures 3 and S2. In alignment with the HMG-box_ROX1-like domain sequence of the reference protein AEH27625, the domain sequences of the MAT1-2-1 proteins derived from the 5 *O. sinensis* strains are truncated differentially at the C-termini and each contains a tyrosine-to-histidine (Y-to-H) substitution, contributing to diverse 3D structures under 4 AlphaFold codes (Figure 4). The truncated MAT1-2-1 protein AGW27554 contains an additional glutamine-to-threonine (Q-to-T) substitution, as shown in Figure 3.

**Figure 3.**
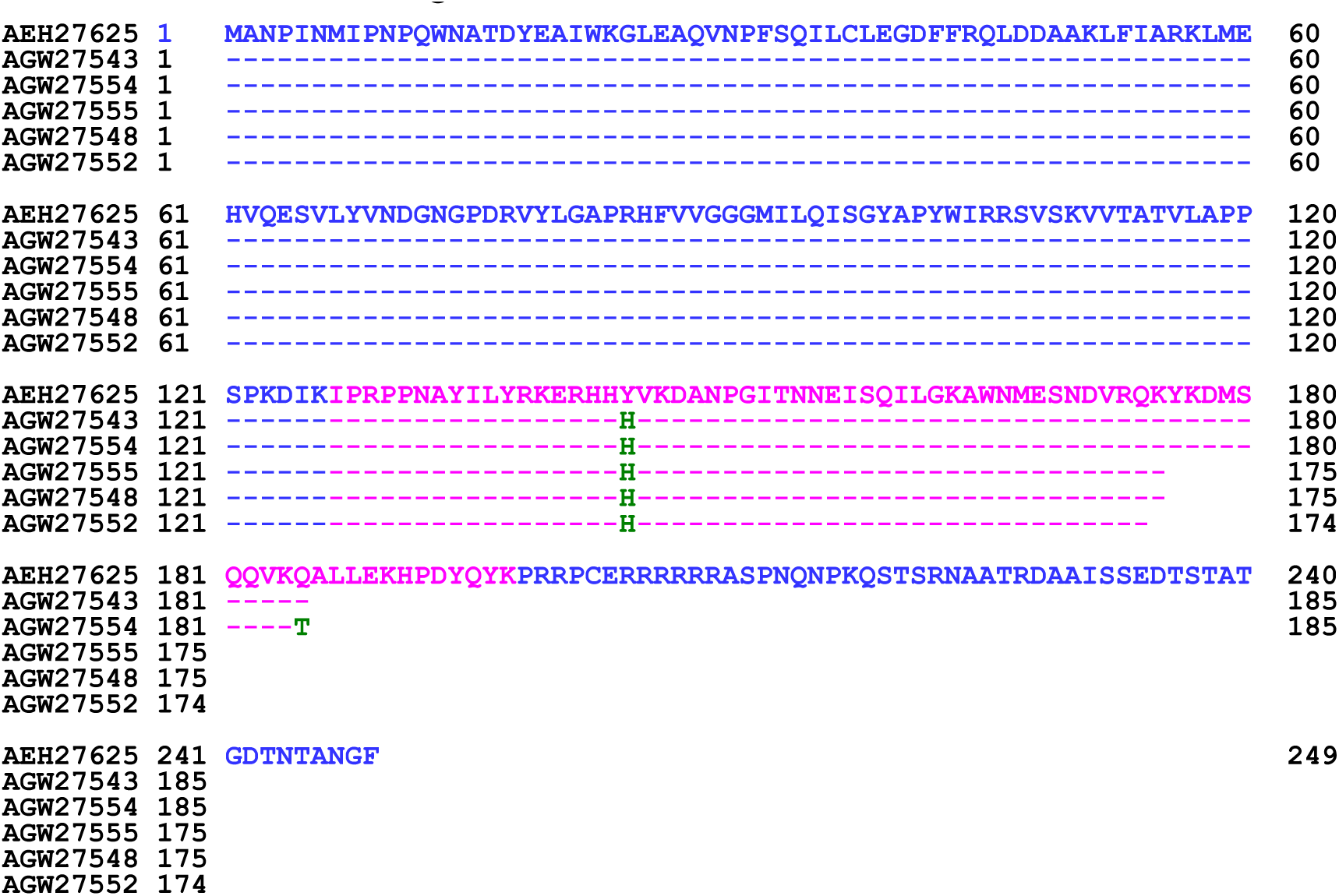
Alignment of the sequences of 5 truncated AMT1-2-1 proteins derived from the *O. sinensis* strains and the reference query of the full-length MAT1-2-1 protein AEH27625 from the *H. sinensis* strain CS2 under AlphaFold code D7F2E9. The amino acids in the HMG-box_ROX1-like domain (amino acids 127→197 in the AEH27625 sequence) are shown in pink, and those outside the domain are shown in blue. The amino acid substitutions are shown in green. The hyphens indicate identical amino acids, and the spaces denote unmatched sequence gaps.

**Figure 4.**
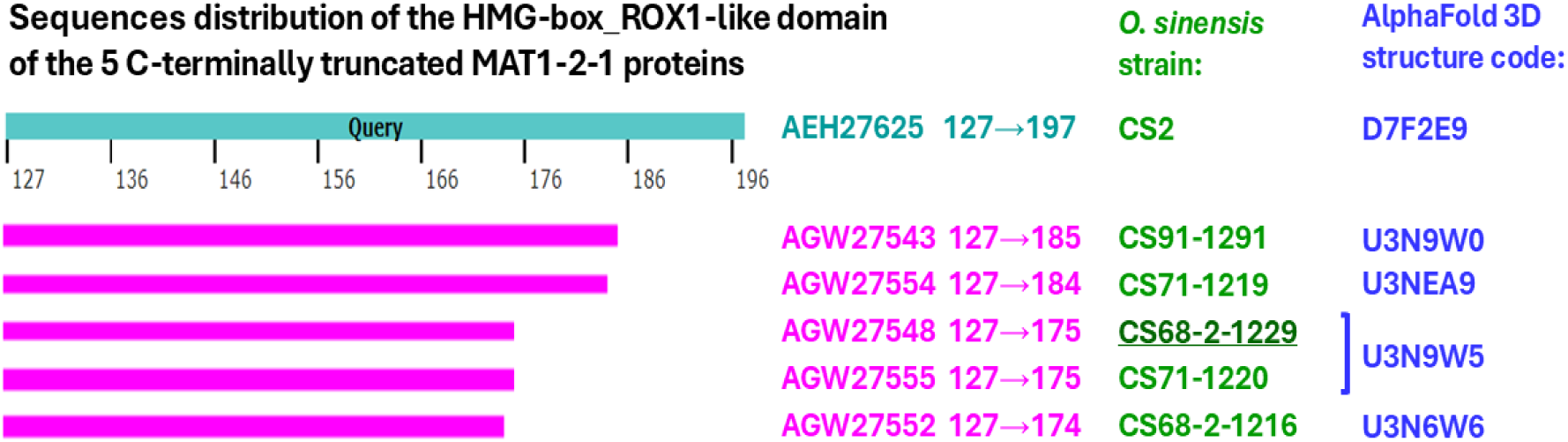
Sequence distribution of the C-terminally truncated HMG-box_ROX1-like domains of 5 MAT1-2-1 proteins derived from the *O. sinensis* strains (*cf*. Table 1). The sequences are aligned with the HMG-box_ROX1-like domain (127→197) of the full-length MAT1-2-1 protein AEH27625 from *H. sinensis* strain CS2.

### 3.3 Heteromorphic tertiary structures of the MATα_HMGbox domains of the truncated MAT1-1-1 proteins derived from *O. sinensis* strains

This section reports the results of the correlation analyses between changes in the hydrophobicity and structures of the MATα_HMGbox domains of truncated MAT1-1-1 proteins derived from various *O. sinensis* strains, which correspond to 9 AlphaFold codes (Table 1, Figures 1−2), compared to the reference MAT1-1-1 proteins. The reference protein, represented by AGW27560 (derived from the *O. sinensis* strain CS68-2-1229; Table S4), is associated with the AlphaFold code U3N942 [14]. The sequences of the MATα_HMGbox domain (amino acid residues 51→225) were extended by 9 amino acid residues both upstream and downstream of the domains and were used in the sequential ExPASy ProtScale plot to analyze hydropathy, α-helices, β-sheets, β-turns, and coils with a window size of 9 amino acid residues (*cf*. Section 2.3).

Figure 5 compares the hydrophobicity and structural characteristics of the MATα_HMGbox domains between 2 groups of truncated MAT1-1-1 proteins: (1) the protein AGW27522 (under the AlphaFold code U3N6U0) derived from the *O. sinensis* strain CS560-961; and (2) 8 proteins, AGW27517, AGW27518, AGW27519, AGW27520, AGW27521, AGW27523, AGW27524, and AGW27528 (all under the AlphaFold code U3N9T9), derived from *O. sinensis* strains CS6-251, CS36-1294, CS37-295, CS18-266, CS26-277, CS91-1291, CS34-291, and CS68-2-1229, respectively (Table 1). The reference protein AGW27560 (under AlphaFold code U3N942) is derived from the *H. sinensis* strain CS68-2-1229 [14]. Panel (A) of Figure 5 shows that the MATα_HMGbox domains of both groups of subject proteins exhibit 13-residue truncations at N-termini of the DNA-binding domains. The truncations result in changes in the topological structure and waveform of the ExPASy ProtScale plots, as illustrated in Panels (B)−(F) for hydropathy, α-helices, β-sheets, β-turns, and coils, respectively. Panel (G) shows the 3D structure of the full-length reference protein AGW27560, where the 3 α-helices within the MATα_HMGbox domain form a core L-shaped stereostructure. Panels (H)−(I) illustrate that the N-and C-terminal truncations (*cf*. Figure 1) significantly alter the 3D structures of the proteins under the heteromorphic AlphaFold codes U3N6U0 and U3N9T9, respectively. The N-terminally truncated regions of these variant proteins are located upstream of the core structure of 3 α-helices and may exert a steric auxiliary effect on stabilizing the hydrophobic core structures of the MATα_HMGbox domains [90,91].

**Figure 5.**
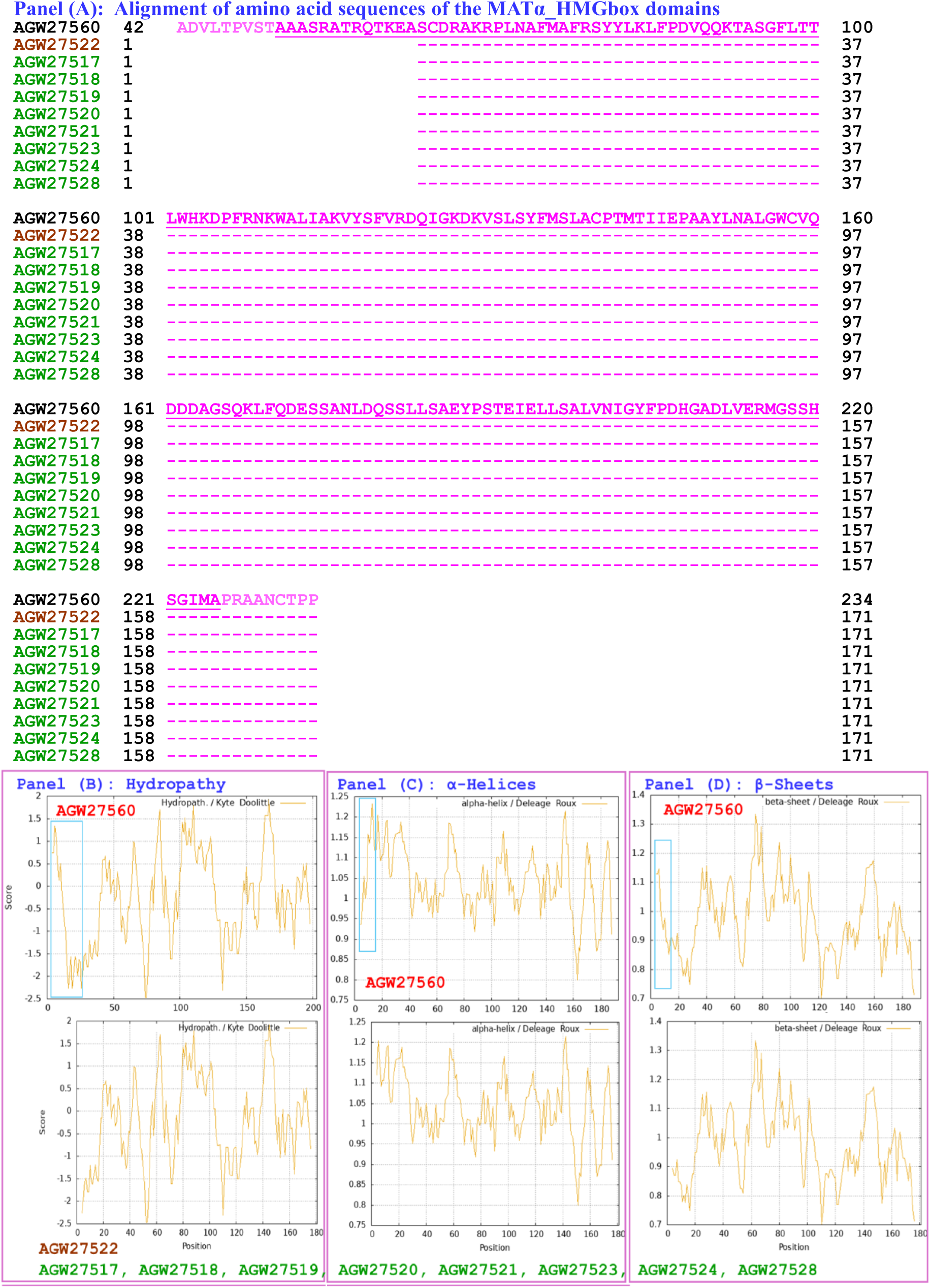
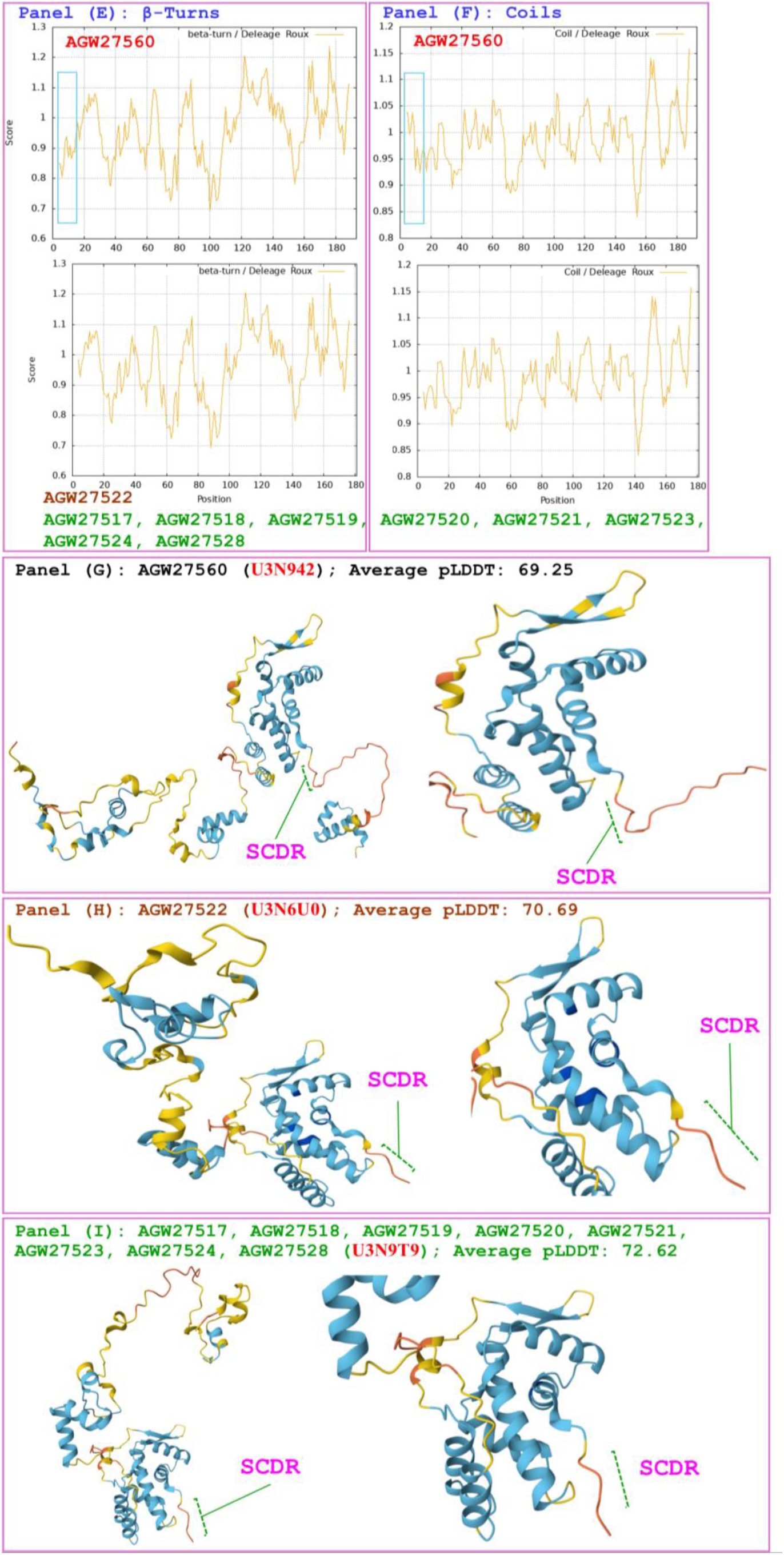
Correlations of the changes in the hydrophobicity and the primary, secondary, and tertiary structures of the MATα_HMGbox domains of MAT1-1-1 proteins: the reference protein AGW27560 (under the AlphaFold code U3N942) derived from H. sinensis strain CS68-2-1229 and the 2 groups of the truncated proteins: (1) AGW27522 shown in brown (under the AlphaFold code U3N6U0) derived from the O. sinensis strain CS560-961 and (2) the truncated proteins shown in green (under the AlphaFold code U3N9T9) derived from the O. sinensis strains CS6-251, CS36-1294, CS37-295, CS18-266, CS26-277, CS91-1291, CS34-291, and CS68-2-1229. Panel (A) shows an alignment of the amino acid sequences of the MATα_HMGbox domains of the MAT1-1-1 proteins, where the hyphens indicate identical amino acid residues and the spaces denote unmatched sequence gaps. The ExPASy ProtScale plots show the changes in hydrophobicity and the 2D structures in Panels (B) –(F) for hydropathy, α-helices, β-sheets, β-turns, and coils of the proteins, respectively; the open rectangles in blue highlight the N-terminally truncated region in the ExPASy plots. Panels (G)–(I) show the 3D structures of the full-length proteins on the left, and the locally magnified structures surrounding the truncated region are shown on the right. The model confidence for the AlphaFold-predicted 3D structures is as follows: 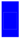 very high (pLDDT>90); 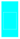 high (90>pLDDT>70); 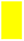 low (70>pLDDT>50); and 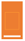 very low (pLDDT<50).

The hydrophobicity and structural characteristics of the MATα_HMGbox domain of the truncated MAT1-1-1 protein AGW27525 are compared with those of the reference full-length protein AGW27560 in Figure 6. The protein AGW27525 (under AlphaFold code U3N919) is derived from the *O. sinensis* strain CS76-1284 (Table 1), while the reference protein AGW27560 (under AlphaFold code U3N942) is derived from the *H. sinensis* strain CS68-2-1229 [14]. Panel (A) of Figure 6 shows a 16-residue truncation at the N-terminus of the MATα_HMGbox domain of the protein AGW27525. This truncation results in altered topological structures and waveforms in the ExPASy ProtScale plots, as illustrated in Panels (B)−(F) for hydropathy, α-helices, β-sheets, β-turns, and coils, respectively. Compared with the 3D structure of the reference protein AGW27560 shown in Panel (G), Panel (H) demonstrates that the truncated variant significantly alters the 3D structure of the protein AGW27525 under the AlphaFold code U3N919. Notably, the N-terminal amino acids 2−4 (AKR) shown in Panel (A) have hydropathy indices of 1.8, -3.9, and -4.5, respectively (Table S3) [73] and collectively exhibit hydrophilic characteristics. These amino acids initiate the formation of the β-sheet structure in the truncated protein AGW27525 shown in Panel (H); in contrast, the amino acids AKR in the full-length reference protein AGW27560 shown in Panel (G) are not part of the β-sheet structure or the hydrophobic core composed of the 3 critical α-helices within the MATα_HMGbox domain. The truncated region of the AGW27525 protein is located upstream of the 3 α-helix core within the MATα_HMGbox domain and may not directly participate in stabilizing the hydrophobic core of this domain [90,91].

**Figure 6.**
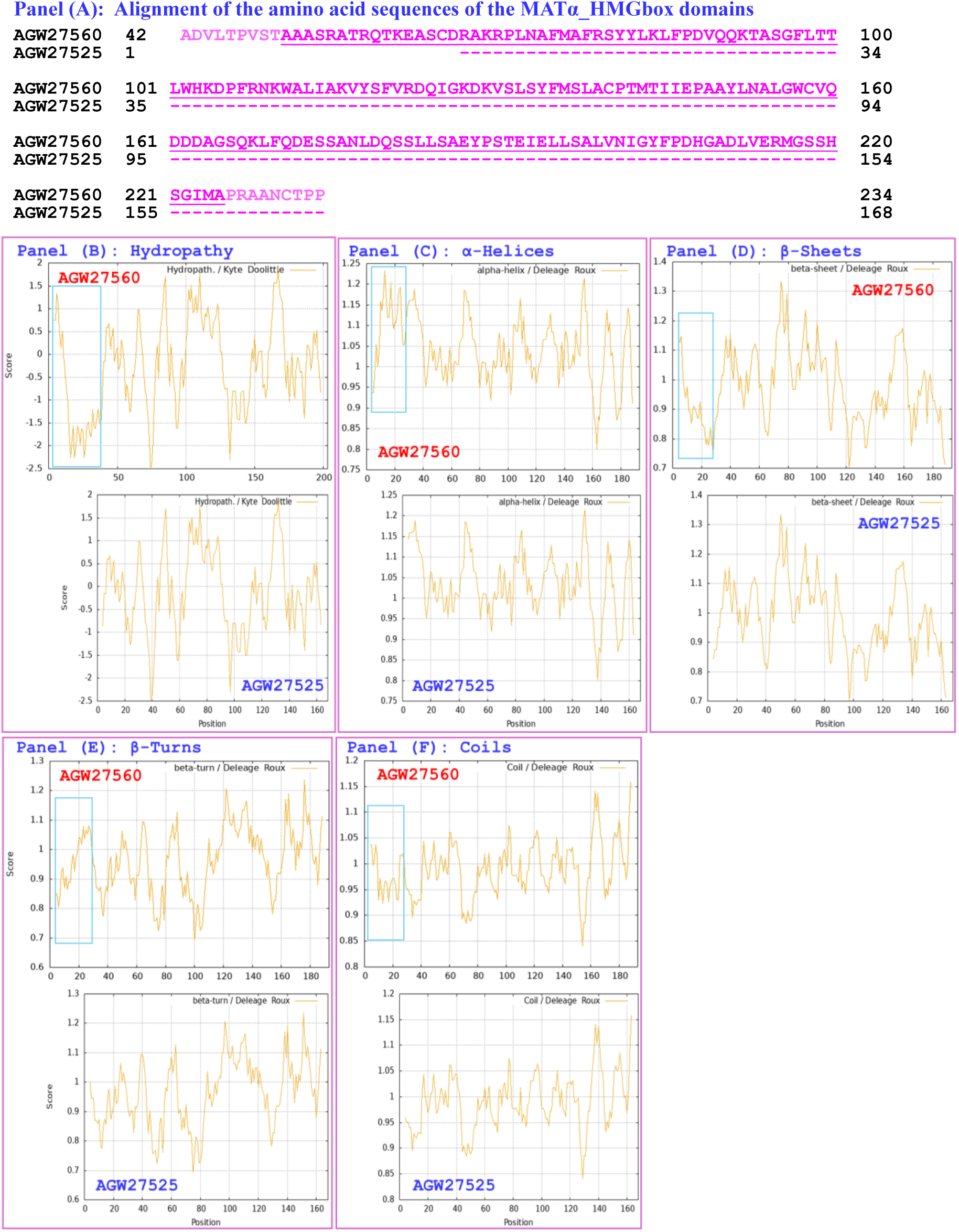
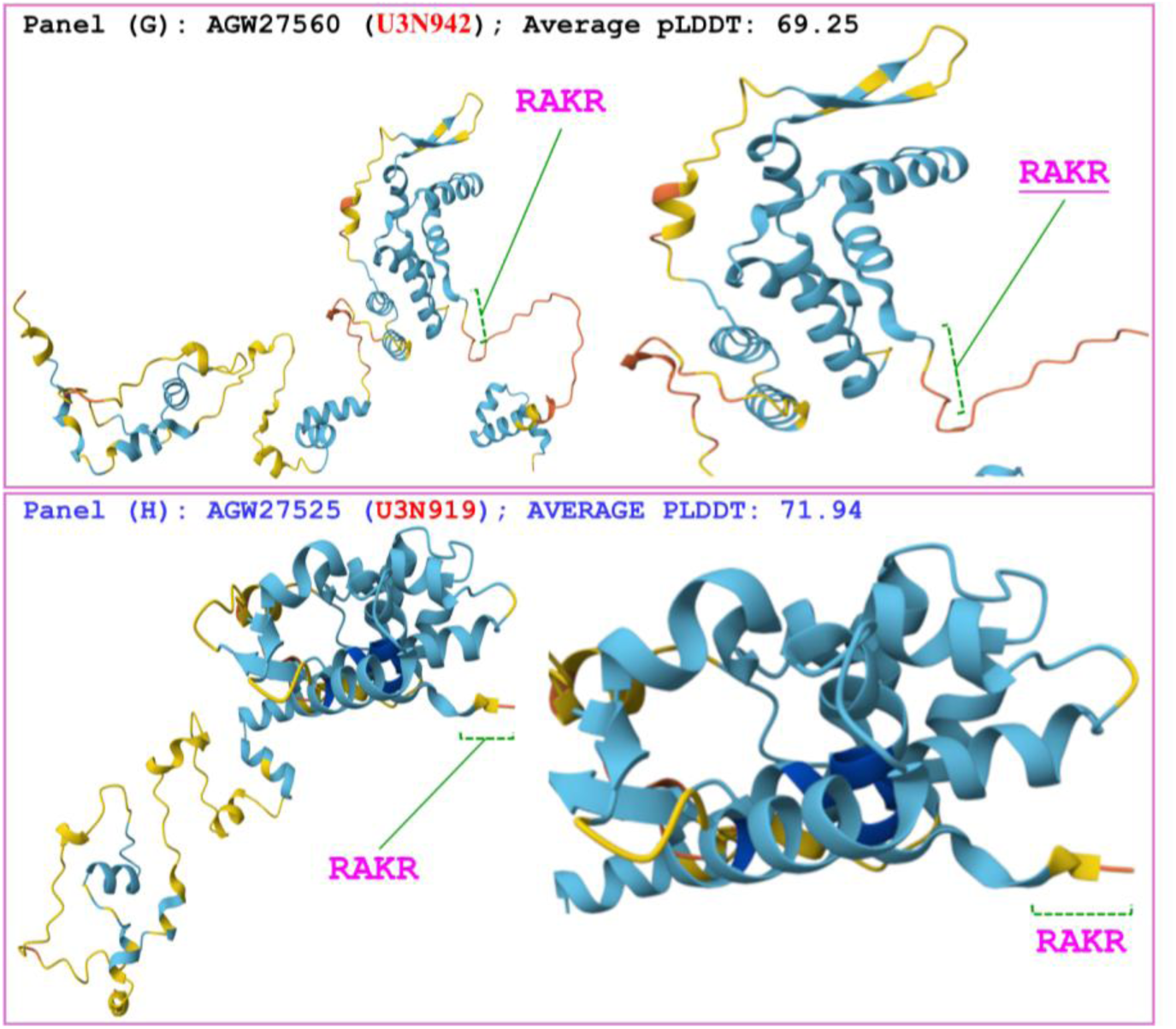
Correlations of the changes in hydrophobicity and the primary, secondary, and tertiary structures of the MATα_HMGbox domains of MAT1-1-1 proteins: the reference protein AGW27560 (under the AlphaFold code U3N942) derived from the *H. sinensis* strain CS68-2-1229 and the variant protein AGW27525 (under the AlphaFold code U3N919) derived from the *O. sinensis* strain CS76-1284. Panel (A) shows an alignment of the amino acid sequences of the MATα_HMGbox domains of the MAT1-1-1 proteins, where the hyphens indicate identical amino acid residues and the spaces denote unmatched sequence gaps. The ExPASy ProtScale plots show the changes in hydrophobicity and the 2D structure in Panels (B)−(F) for hydropathy, α-helices, β-sheets, β-turns, and coils of the protein, respectively; the open rectangles in blue highlight the N-terminally truncated region in the ExPASy plots. Panels (G)−(H) show the 3D structures of the full-length proteins on the left, and the locally magnified structures surrounding the variation region are shown on the right. The model confidence for the AlphaFold-predicted 3D structures is as follows: 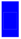 very high (pLDDT>90); 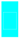 high (90>pLDDT>70); 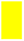 low (70>pLDDT>50); and 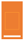 very low (pLDDT<50).

The hydrophobicity and structural characteristics of the MATα_HMGbox domain of the variant MAT1-1-1 protein AGW27526 are compared with those of the full-length reference protein AGW27560 in Figure 7. The protein AGW27526 (under the AlphaFold code U3N7G5) is derived from the *O. sinensis* strain CS561-964 (Table 1), while the reference protein AGW27560 (under the AlphaFold code U3N942) is derived from the *H. sinensis* strain CS68-2-1229 [14]. Panel (A) of Figure 7 shows a 13-residue truncation at the N-terminus of the MATα_HMGbox domain of the protein AGW27526, along with 4 amino acid substitutions: C-to-S (cysteine to serine, both with polar neutral side chains; hydropathy indices changed from -3.5 to -0.8; *cf*. Table S3), D-to-S (acidic aspartic acid to serine with polar neutral side chains; hydropathy indices changed from 2.5 to -0.8), R-to-F (basic arginine to aromatic phenylalanine; hydropathy indices changed from -4.5 to 2.8), and A-to-T (aliphatic alanine to threonine with polar neutral side chains; hydropathy indices changed from 1.8 to -0.7) (Table S3) [73]. The truncation and amino acid substitutions result in changes in the hydrophobicity and secondary structures of the variant protein, which are illustrated by the altered topological structure and waveform in the ExPASy ProtScale plots for hydropathy, α-helices, β-sheets, β-turns, and coils in Panels (B)−(F), respectively. Compared with the 3D structure of the reference protein AGW27560 shown in Panel (G), the truncation and replacement of amino acids in the variant protein AGW27526 shown in Panel (H) are located upstream of the 3 core α-helices of the MATα_HMGbox domain. These modifications caused apparent changes in the tertiary structure of the variant protein AGW27526 under the AlphaFold code U3N7G5.

**Figure 7.**
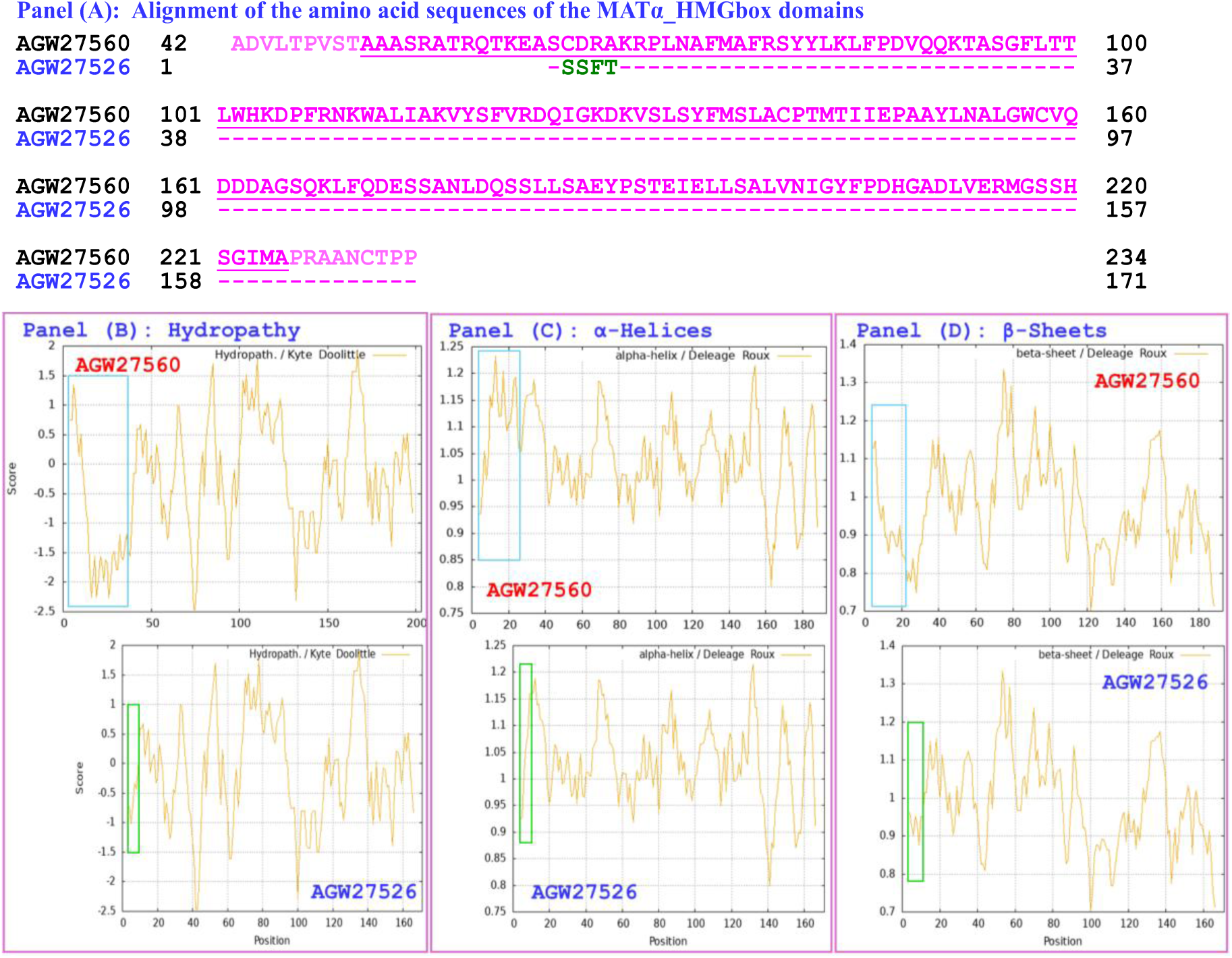
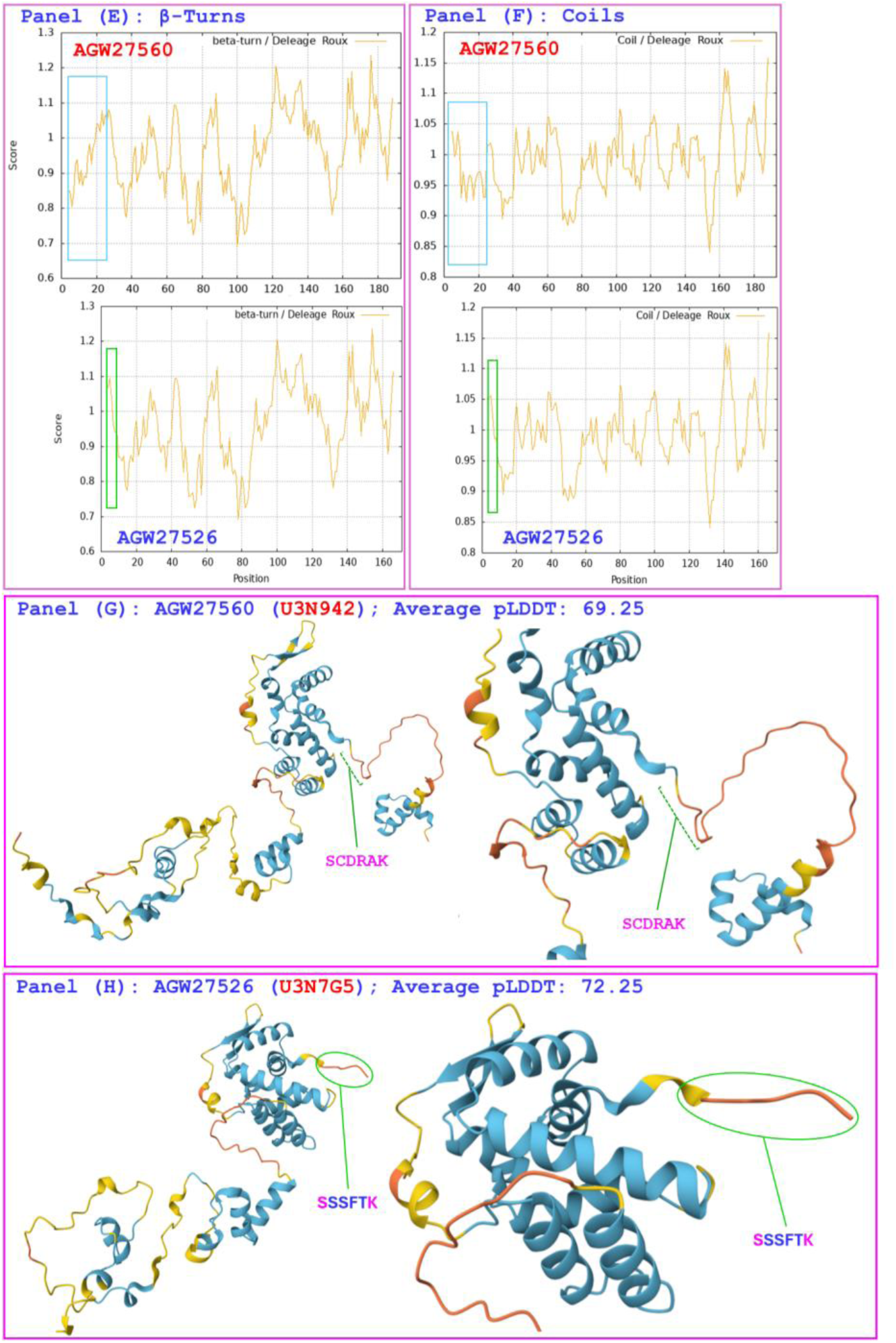
Correlations of the changes in the hydrophobicity and the primary, secondary, and tertiary structures of the MATα_HMGbox domains of MAT1-1-1 proteins: the reference protein AGW27560 (under the AlphaFold code U3N942) derived from the H. sinensis strain CS68-2-1229 and the variant protein AGW27526 (under the AlphaFold code U3N7G5) derived from the O. sinensis strain CS561-964. Panel (A) shows an alignment of the amino acid sequences of the MATα_HMGbox domains of the MAT1-1-1 proteins, where the hyphens indicate identical amino acid residues and the spaces denote unmatched sequence gaps. The ExPASy ProtScale plots show the changes in hydrophobicity and the 2D structure in Panels (B)–(F) for hydropathy, α-helices, β-sheets, β-turns, and coils of the protein, respectively; the open rectangles in blue highlight the N-terminally truncated regions, and those in green highlight the altered topological structure and waveform in the ExPASy plots. Panels (G)–(H) show the 3D structures of the full-length proteins on the left, and the locally magnified structures surrounding the regions of variation are shown on the right. The model confidence for the AlphaFold-predicted 3D structures is as follows: 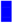 very high (pLDDT>90); 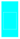 high (90>pLDDT>70); 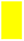 low (70>pLDDT>50); and 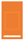 very low (pLDDT<50).

The hydrophobicity and structural characteristics of the MATα_HMGbox domain of the variant MAT1-1-1 protein AGW27527 are compared with those of the reference protein AGW27560 in Figure 8. The protein AGW27527 (under the AlphaFold code U3N6U4) is derived from the *O. sinensis* strain CS25-273 (Table 1), while the reference protein AGW27560 (under the AlphaFold code U3N942) is derived from the *H. sinensis* strain CS68-2-1229 [14]. Panel (A) of Figure 8 shows a 27-residue truncation at the N-terminus of the MATα_HMGbox domain of the protein AGW27527. This truncation results in changes in the topological structure and waveform in the ExPASy ProtScale plots, which illustrate hydropathy, α-helices, β-sheets, β-turns, and coils in Panels (B)−(F), respectively. Compared with the 3D structure of the reference protein AGW27560 shown in Panel (G), Panel (H) shows that the truncation is located within the first of the 3 core α-helices of the MATα_HMGbox domain and that the tertiary structure of the truncated protein AGW27527 is apparently altered under the AlphaFold code U3N6U4.

**Figure 8.**
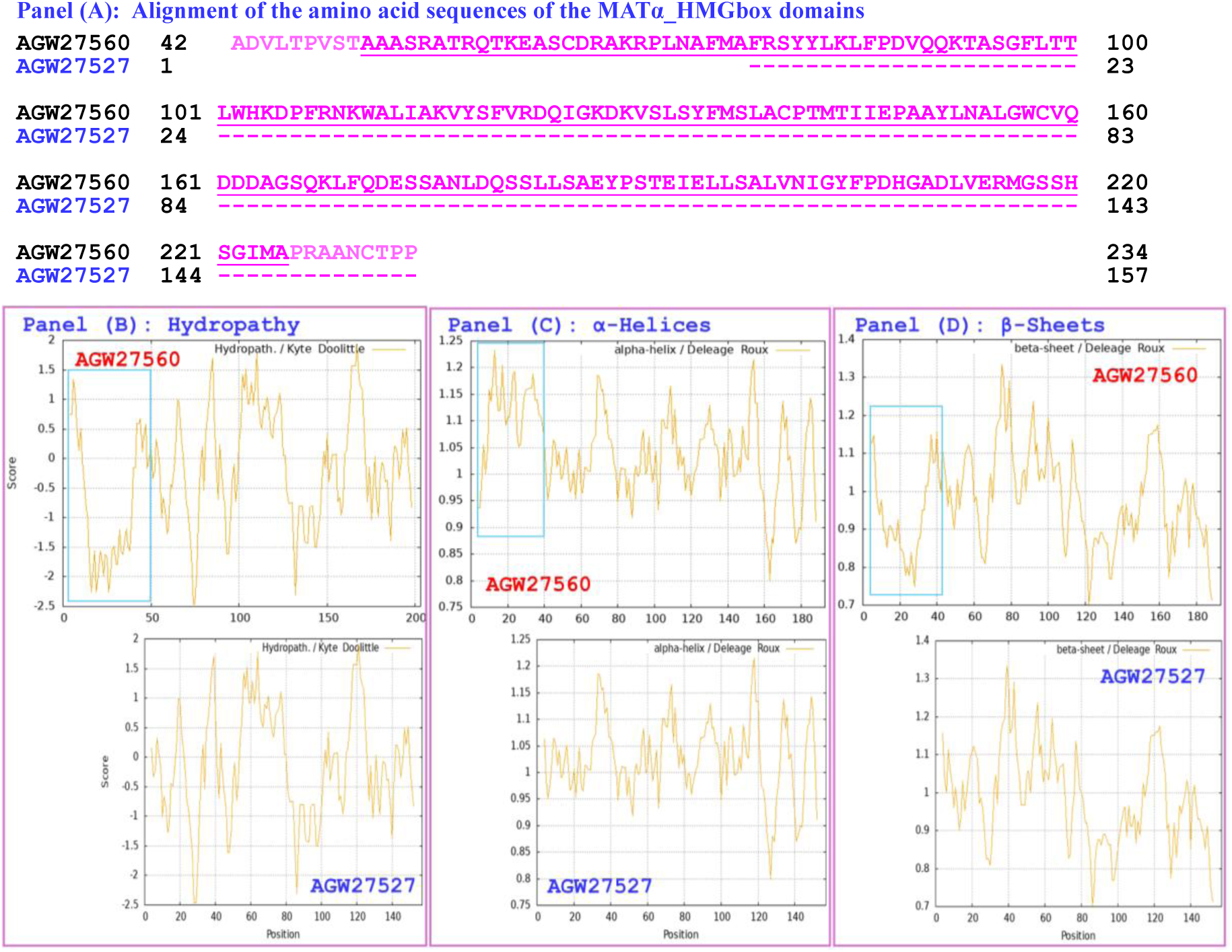
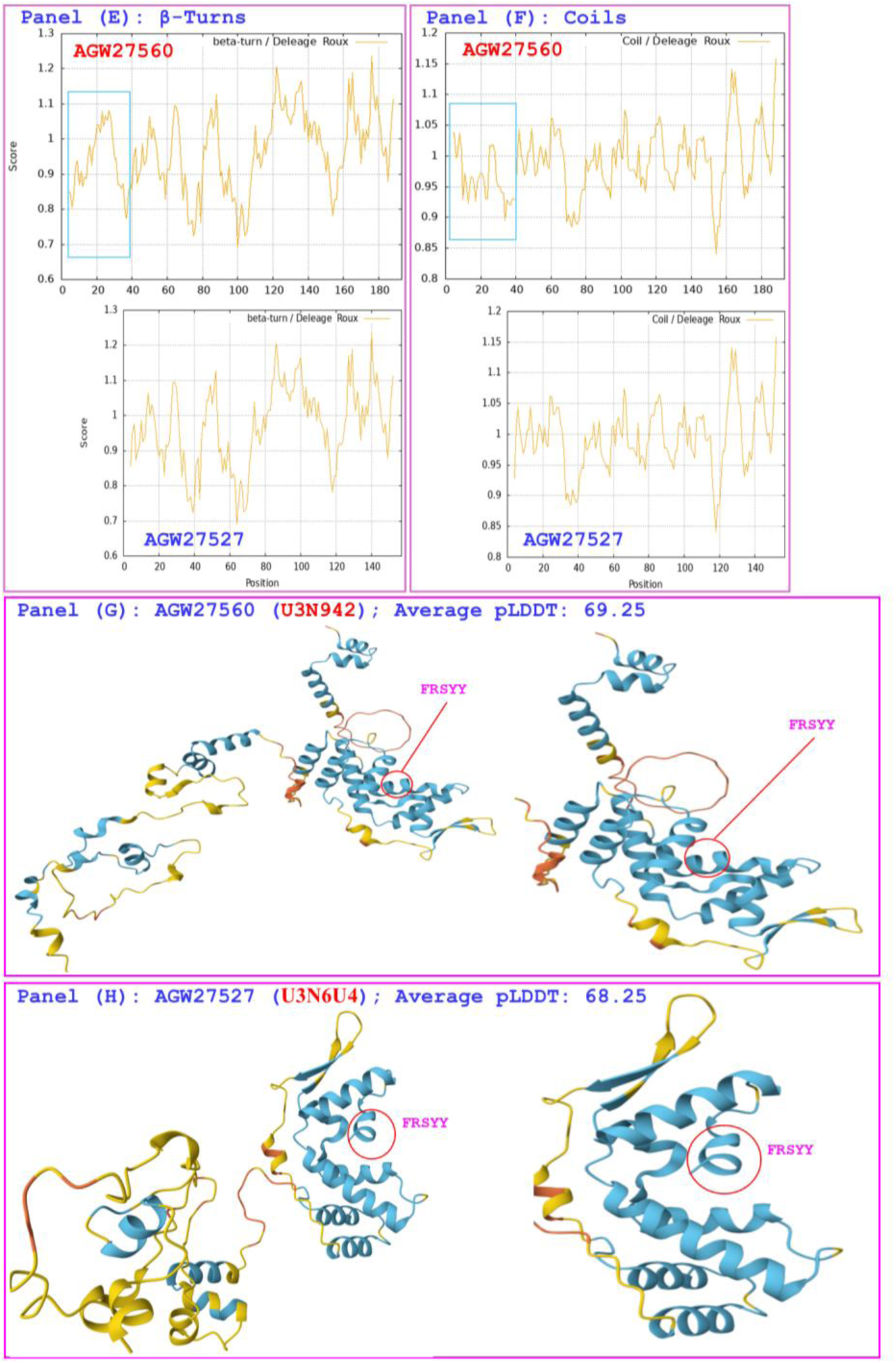
Correlations of the changes in the hydrophobicity and the primary, secondary, and tertiary structures of the MATα_HMGbox domains of MAT1-1-1 proteins: the reference protein AGW27560 (under the AlphaFold code U3N942) derived from the H. sinensis strain CS68-2-1229 and the variant protein AGW27527 (under the AlphaFold code U3N6U4) derived from the O. sinensis strain CS25-273. Panel (A) shows an alignment of the amino acid sequences of the MATα_HMGbox domains of the MAT1-1-1 proteins, where the hyphens indicate identical amino acid residues and the spacesdenoteunmatched sequence gaps. The ExPASy ProtScale plots) show the changes in hydrophobicity and the 2D structure in Panels (B)–(F for hydropathy, α-helices, β-sheets, β-turns, and coils of the protein, respectively; the open rectangles in blue highlight the N-terminal truncation regions in the ExPASy plots. Panels (G)–(H) show the 3D structures of the full-length proteins on the left, and the locally magnified structures surrounding the truncated region are shown on the right. The model confidence for the AlphaFold-predicted 3D structures is as follows: 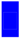 very high (pLDDT>90); 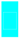 high (90>pLDDT>70); 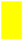 low (70>pLDDT>50); and 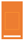 very low (pLDDT<50).

The hydrophobicity and structural characteristics of the MATα_HMGbox domain of the variant MAT1-1-1 proteins AGW27532, AGW27533, and AGW27535 are compared with those of the reference protein AGW27560 in Figure 9. These three variant proteins (all under the AlphaFold code U3N6U8) are derived from the *O. sinensis* strains CS68-5-1216, CS71-1218, and CS71-1220 (Table 1), whereas the reference protein AGW27560 (under the AlphaFold code U3N942) is derived from the *H. sinensis* strain CS68-2-1229 [14]. Panel (A) of Figure 9 shows 47-residue truncations at the N-termini of the MATα_HMGbox domains of the variant proteins. These truncations result in changes in the topological structure and waveform in the ExPASy ProtScale plots, which illustrate hydropathy, α-helices, β-sheets, β-turns, and coils in Panels (B)−(F), respectively. A comparison of the tertiary structures of the MAT1-1-1 proteins is illustrated in Panels (G)−(H). The truncations are located within the first of the 3 core α-helices of the MATα_HMGbox domains, which results in significant changes in the tertiary structures of the variant proteins AGW27532, AGW27533, and AGW27535 under the AlphaFold code U3N6U8.

**Figure 9.**
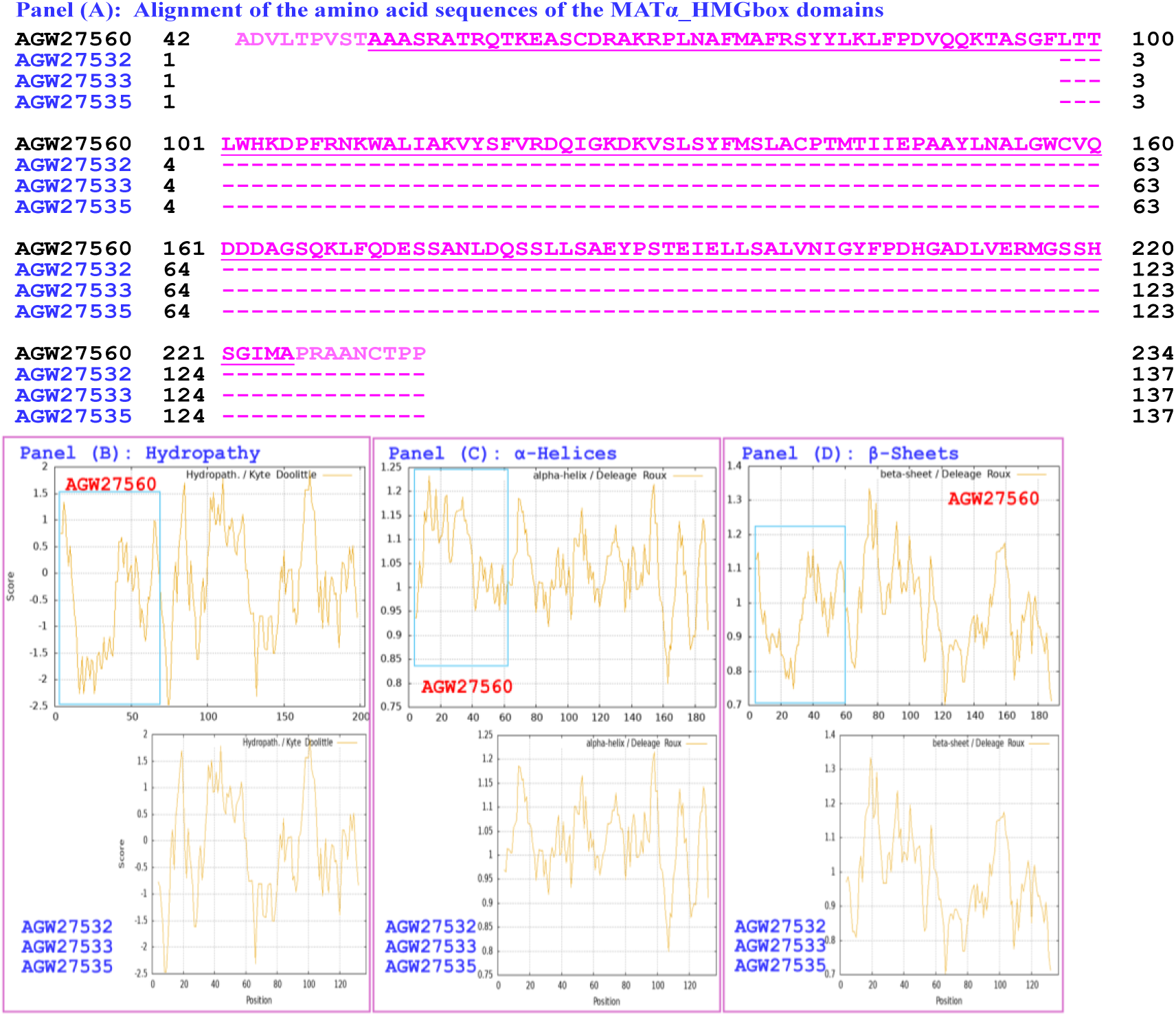
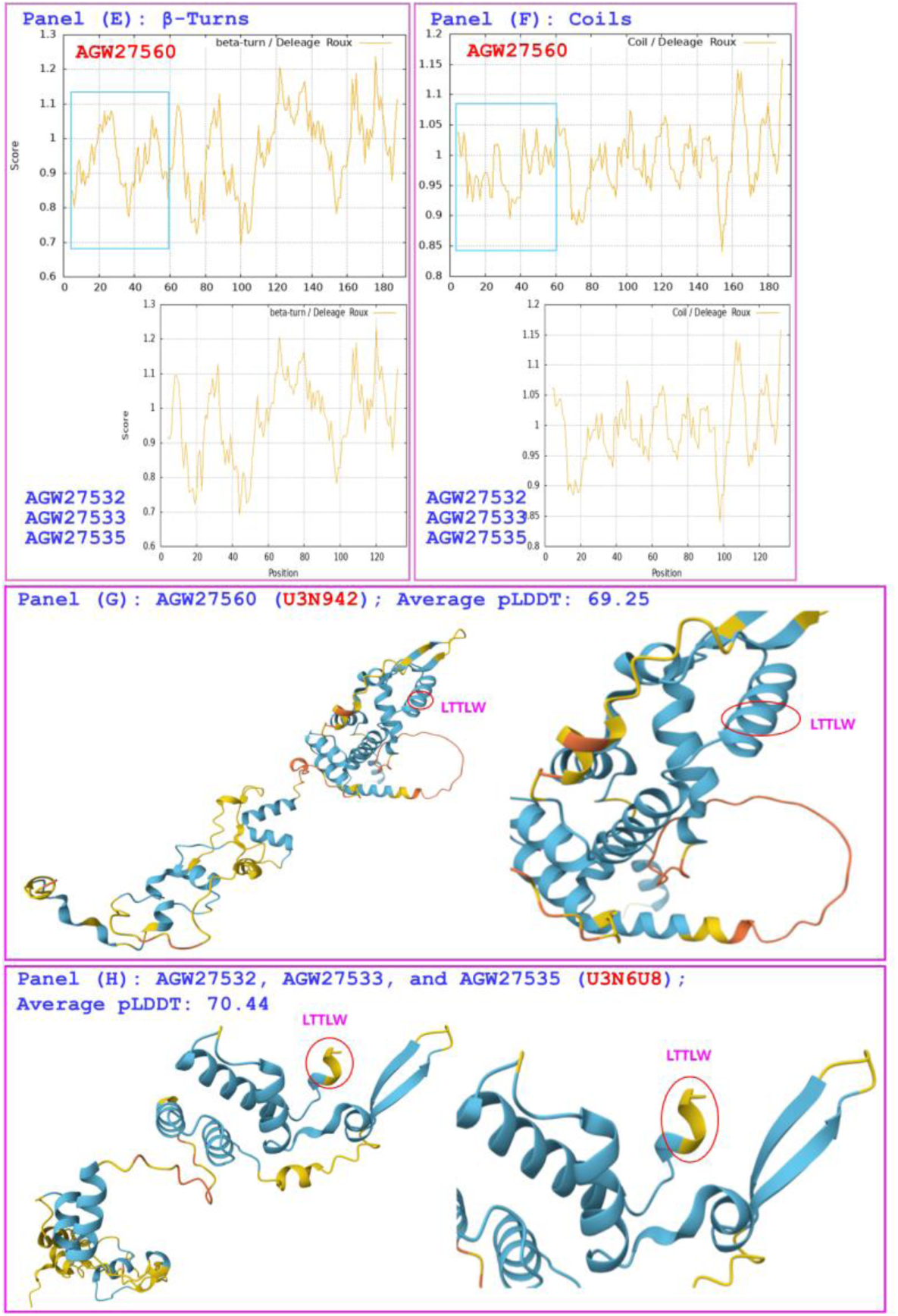
Correlations of the changes in the hydrophobicity and the primary, secondary, and tertiary structures of the MATα_HMGbox domains of MAT1-1-1 proteins: the reference protein AGW27560 (under the AlphaFold code U3N942) derived from the H. sinensis strain CS68-2-1229 and the variant proteinsAGW27532, AGW27533, and AGW27535 (under the AlphaFold code U3N6U8) derived from the O. sinensis strains CS68-5-1216, CS71-1218, and CS71-1220, respectively. Panel (A) shows an alignment of the amino acid sequences of the MATα_HMGbox domains of the MAT1-1-1 proteins, where the hyphens indicate identical amino acid residues and the spaces denote unmatched sequence gaps. The ExPASy ProtScale plots show the changes in hydrophobicity and the 2D structure in Panels (B)–(F) for hydropathy, α-helices, β-sheets, β-turns, and coils of the protein, respectively; the open rectangles in blue highlight the N-terminally truncated regions in the ExPASy plots. Panels (G)–(I) show the 3D structures of the full-length proteins on the left and the locally magnified structures surrounding the truncation site on the right. The model confidence for the AlphaFold-predicted 3D structures is as follows: 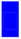 very high (pLDDT>90); 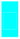 high (90>pLDDT>70); 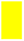 low (70>pLDDT>50); and 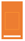 very low (pLDDT<50).

The hydrophobicity and structural characteristics of the MATα_HMGbox domains of the variant MAT1-1-1 proteins AGW27529 and AGW27530 are compared with those of the reference protein AGW27560 in Figure 10. The two variant proteins (under the AlphaFold code U3NE79) are derived from the *O. sinensis* strains CS70-1208 and CS70-1211 (Table 1), whereas the reference protein AGW27560 (under the AlphaFold code U3N942) is derived from the *H. sinensis* strain CS68-2-1229 [14]. Panel (A) of Figure 10 shows 46-residue truncations at the N-termini of the MATα_HMGbox domains of the proteins AGW27529 and AGW27530, along with F-to-V substitutions (aromatic phenylalanine to aliphatic valine; hydropathy indices changed from 2.8 to 4.2 (Table S3) [73] within the MATα_HMGbox domains, indicating increased hydrophobicity. The truncation and increased hydrophobicity result in an altered secondary structure, as illustrated by the changes in topological structure and waveform in the ExPASy ProtScale plots for hydropathy, α-helices, β-sheets, β-turns, and coils shown in Panels (B)−(F), respectively. Compared the 3D structures of the MAT1-1-1 proteins illustrated in Panels (G)−(H), the truncation and replacement of amino acids in the proteins AGW27529 and AGW27530 are located within the first of the 3 core α-helices of the MATα_HMGbox domains. These modifications result in the significantly altered tertiary structures of the proteins AGW27529 and AGW27530 corresponding to the AlphaFold code U3NE79.

**Figure 10.**
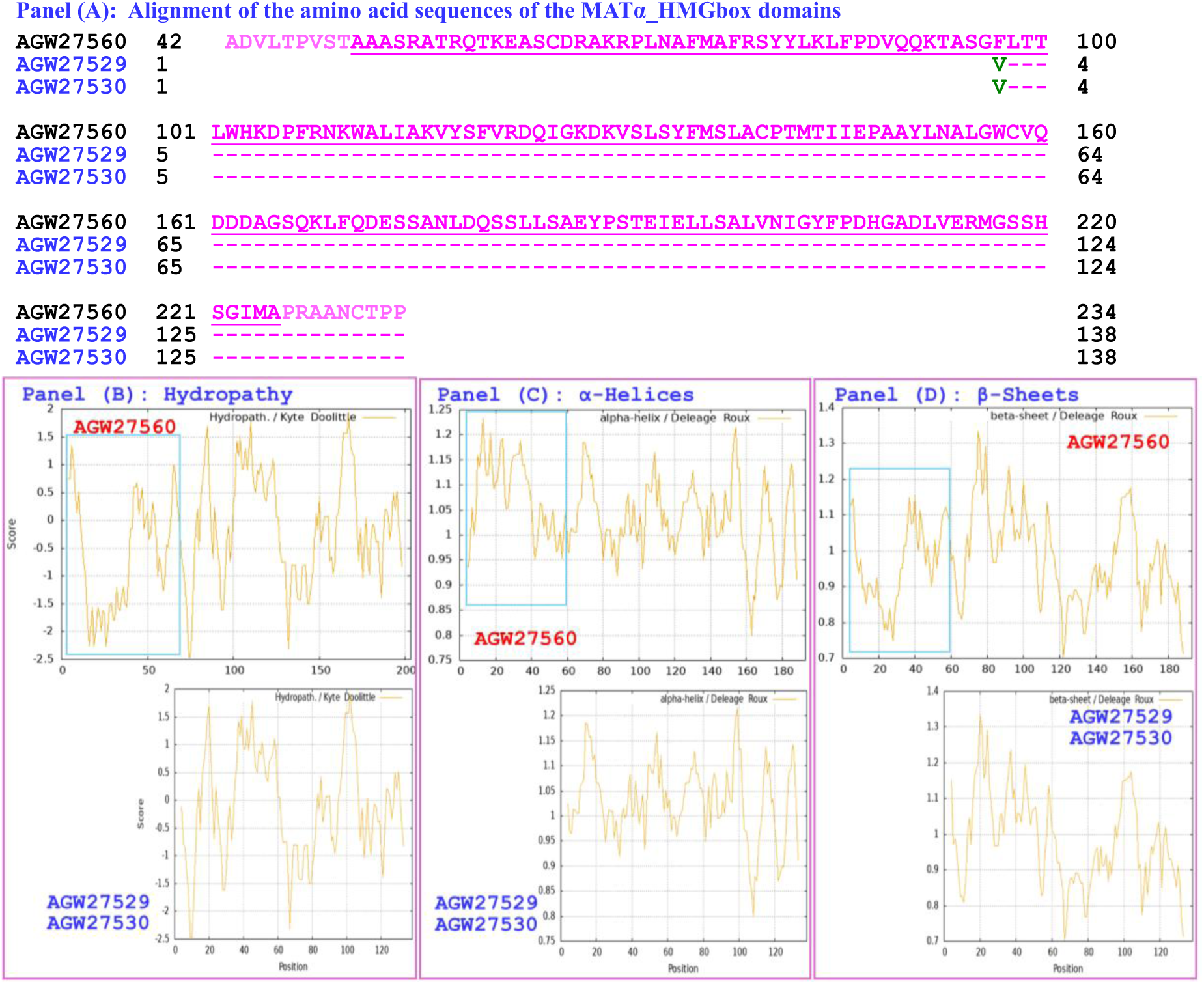
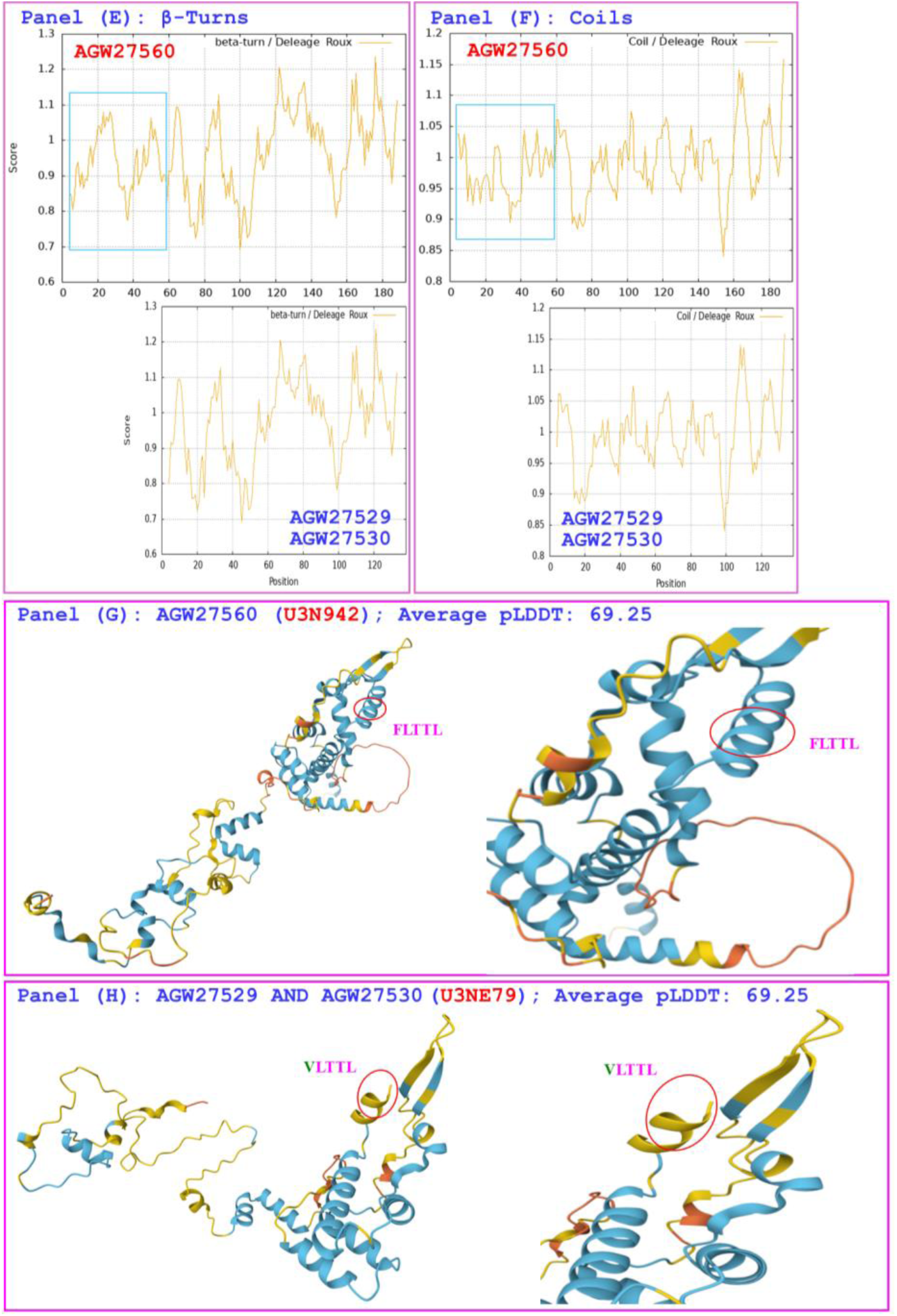
Correlations of the changes in the hydrophobicity and the primary, secondary, and tertiary structures of the MATα_HMGbox domains of MAT1-1-1 proteins: the reference protein AGW27560 (under the AlphaFold code U3N942) derived from the H. sinensis strain CS68-2-1229 and the variant proteins AGW27529 and AGW27530 (under the AlphaFold code U3NE79) derived from the O. sinensis strains CS70-1208 and CS70-1211. Panel (A) shows an alignment of the amino acid sequences of the MATα_HMGbox domains of the MAT1-1-1 proteins; amino acid substitutions are shown in green, the hyphens indicate identical amino acid residues, and the spaces denote unmatched sequence gaps. The ExPASy ProtScale plots show the changes in hydrophobicity and the 2D structure in Panels (B)–(F) for hydropathy, α-helices, β-sheets, β-turns, and coils of the protein, respectively; the open rectangles in blue highlight the N-terminally truncated regions in the ExPASy plots. Panels (G)–(I) show the 3D structures of the full-length proteins on the left, and the locally magnified structures surrounding the variation site are shown on the right. The model confidence for the AlphaFold-predicted 3D structures is as follows: 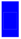 very high (pLDDT>90); 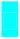 high (90>pLDDT>70); 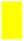 low (70>pLDDT>50); and 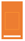 very low (pLDDT<50).

The hydrophobicity and structural characteristics of the MATα_HMGbox domain of the variant MAT1-1-1 protein AGW27536 are compared with those of the reference protein AGW27560 in Figure 11. The protein AGW27536 (under the AlphaFold code U3N7H7) is derived from the *O. sinensis* strain CS68-2-1228 (Table 1), whereas the reference protein AGW27560 (under the AlphaFold code U3N942) is derived from the *H. sinensis* strain CS68-2-1229 [14]. Panel (A) of Figure 11 shows a 44-residue truncation at the N-terminus of the MATα_HMGbox domain of the protein AGW27536, along with an S-to-E substitution and a deletion of leucine within the MATα_HMGbox domain. The S-to-E substitution involves a change from serine (with polar neutral side chains) to acidic glutamic acid, with the hydropathy index changing from -0.8 to -3.5 (Table S3) [73]; the deleted leucine has a hydropathy index of 3.8. The truncation and replacement and deletion of amino acids lead to reduced hydrophobicity and an altered secondary structure, as illustrated by the changes in topological structure and waveform in the ExPASy ProtScale plots for hydropathy, α-helices, β-sheets, β-turns, and coils in Panels (B)−(F), respectively. Compared the 3D structures of the MAT1-1-1 proteins illustrated in Panels (G)−(H), the truncation and replaced and deleted amino acids in the protein AGW27536 are located within the first of the 3 core α-helices of the MATα_HMGbox domain. These modifications result in significant changes in the 3D structure of the protein AGW27536, corresponding to the AlphaFold code U3N7H7.

**Figure 11.**
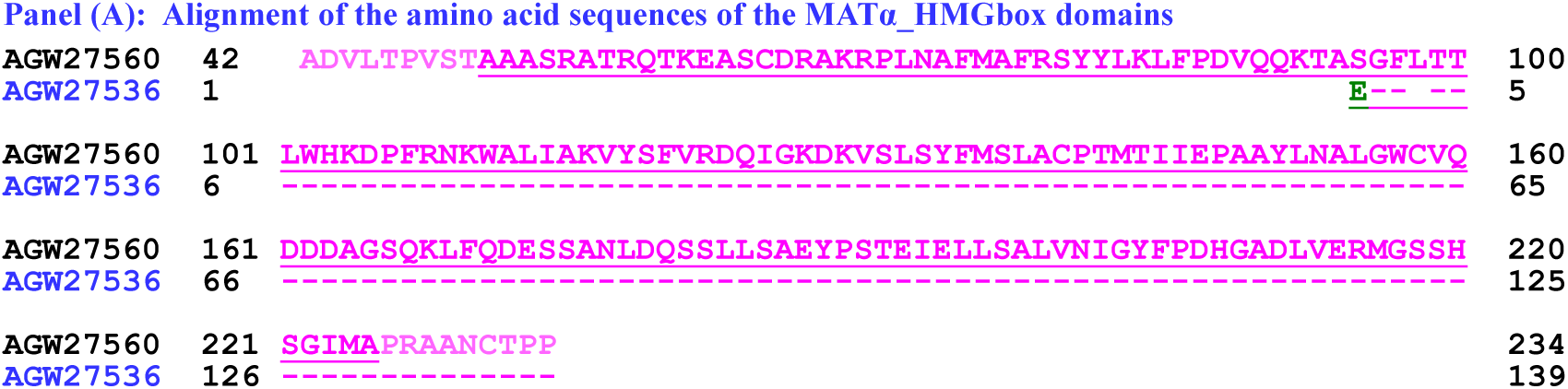
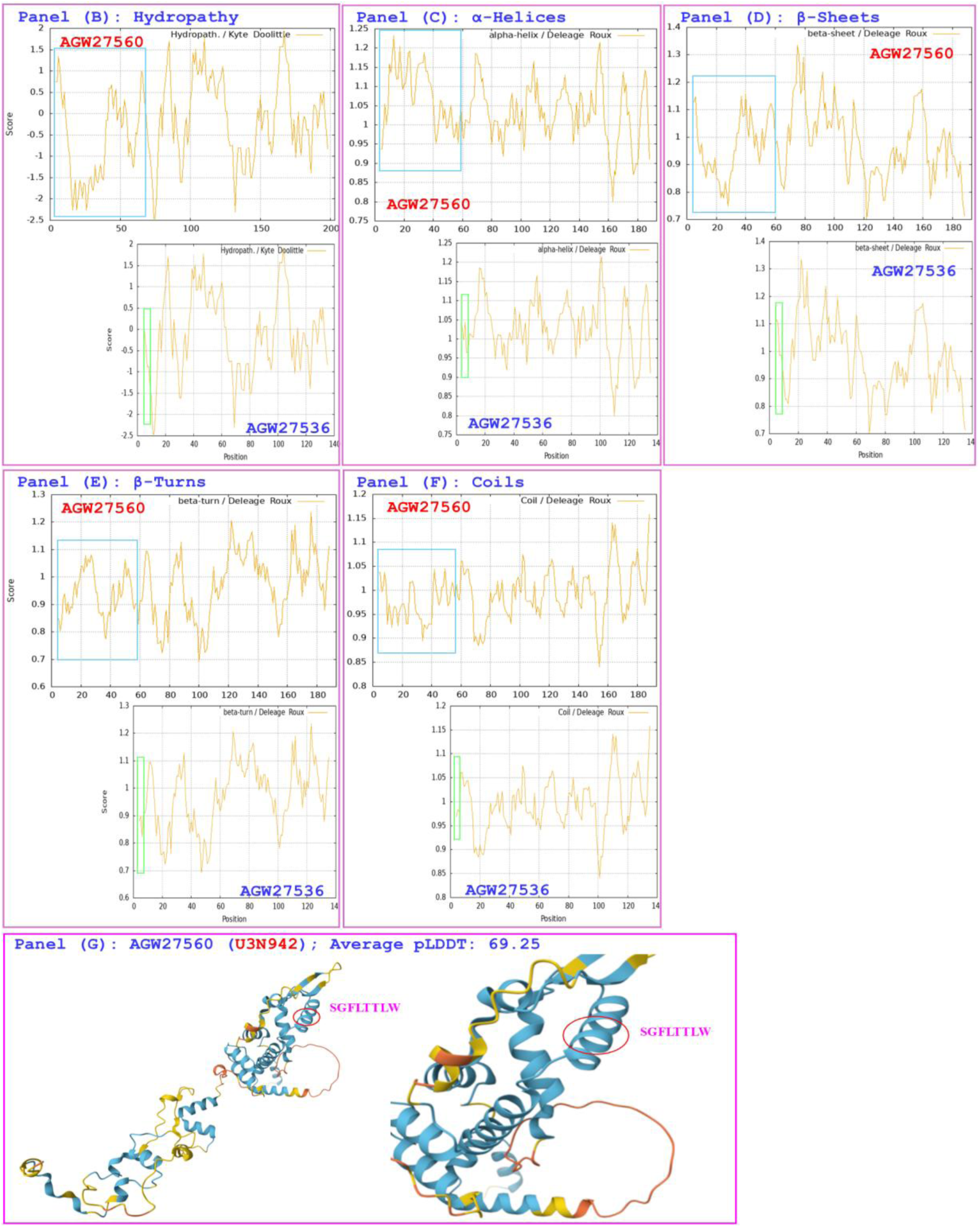
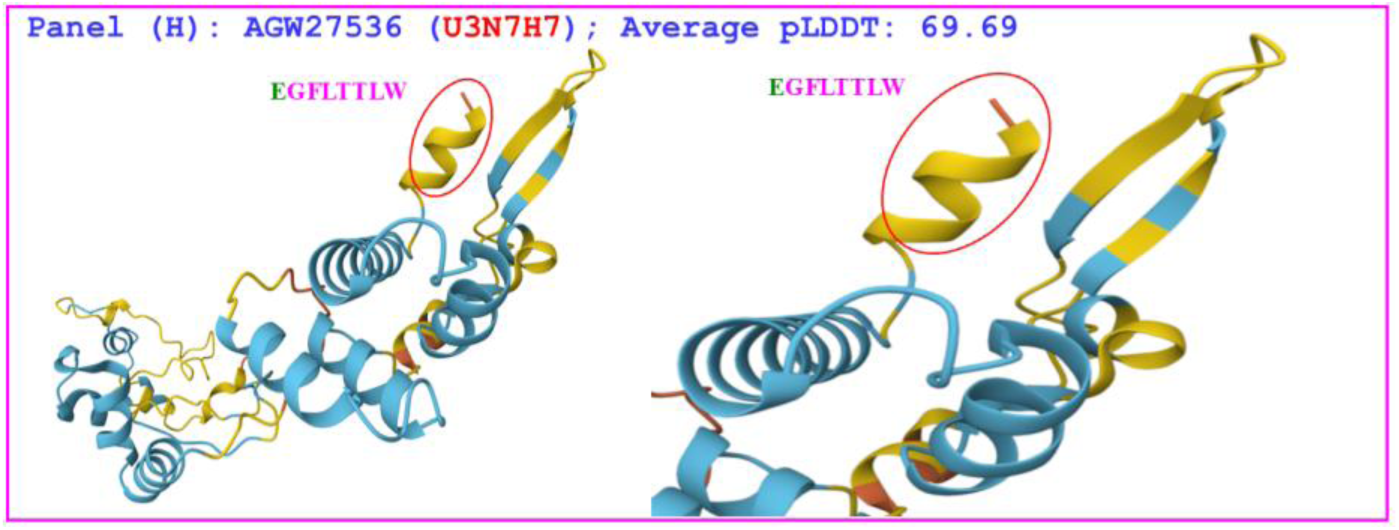
Correlations of the changes in the hydrophobicity and the primary, secondary, and tertiary structures of the MATα_HMGbox domains of MAT1-1-1 proteins: the reference protein AGW27560 (under the AlphaFold code U3N942) derived from the H. sinensis strain CS68-2-1229 and the variant protein AGW27536 (under the AlphaFold code U3N7H7) derived from the O. sinensis strain CS68-2-1228. Panel (A) shows an alignment of the amino acid sequences of the MATα_HMGbox domains of the MAT1-1-1 proteins, where the hyphens indicate identical amino acid residues and the spaces denote unmatched sequence gaps. The ExPASy ProtScale plots show the changes in hydrophobicity and the 2D structure in Panels (B)–(F) for hydropathy, α-helices, β-sheets, β-turns, and coils of the protein, respectively; the open rectangles in blue highlight the N-terminally truncated regions, and those in green highlight the altered topological structure and wave formin the ExPASy plots. Panels (G)–(H) show the 3D structures of the full-length proteins on the left, and the locally magnified structures surrounding the variation site are shown on the right. The model confidence for the AlphaFold-predicted 3D structures is as follows: 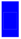 very high (pLDDT>90); 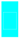 high (90>pLDDT>70); 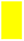 low (70>pLDDT>50); and 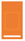 very low (pLDDT<50).

The hydrophobicity and structural characteristics of the MATα_HMGbox domains of the variant MAT1-1-1 proteins AGW27531 and AGW27534 are compared with those of the reference protein AGW27560 in Figure 12. These 2 variant proteins (both under the AlphaFold code U3NE87) are derived from the *O. sinensis* strains CS70-1212 and CS71-1219 (*cf*. Table 1), whereas the reference protein AGW27560 (under the AlphaFold code U3N942) is derived from the *H. sinensis* strain CS68-2-1229 [14]. Panel (A) of Figure 12 shows 60-residue truncations at the N-termini of the MATα_HMGbox domains of the proteins AGW27531 and AGW27534. The truncations result in altered hydrophobicity and secondary structures, as illustrated by the changes in topological structure and waveform in the ExPASy ProtScale plots for hydropathy, α-helices, β-sheets, β-turns, and coils in Panels (B)−(F), respectively. A comparison of the 3D structures of the MAT1-1-1 proteins, as illustrated in Panels (G)−(H), revealed that the truncations in the proteins AGW27531 and AGW27534 are located within the 3 core α-helices of the MATα_HMGbox domains and result in significantly altered tertiary structures of the variant proteins, both of which correspond to the same AlphaFold code U3NE87.

**Figure 12.**
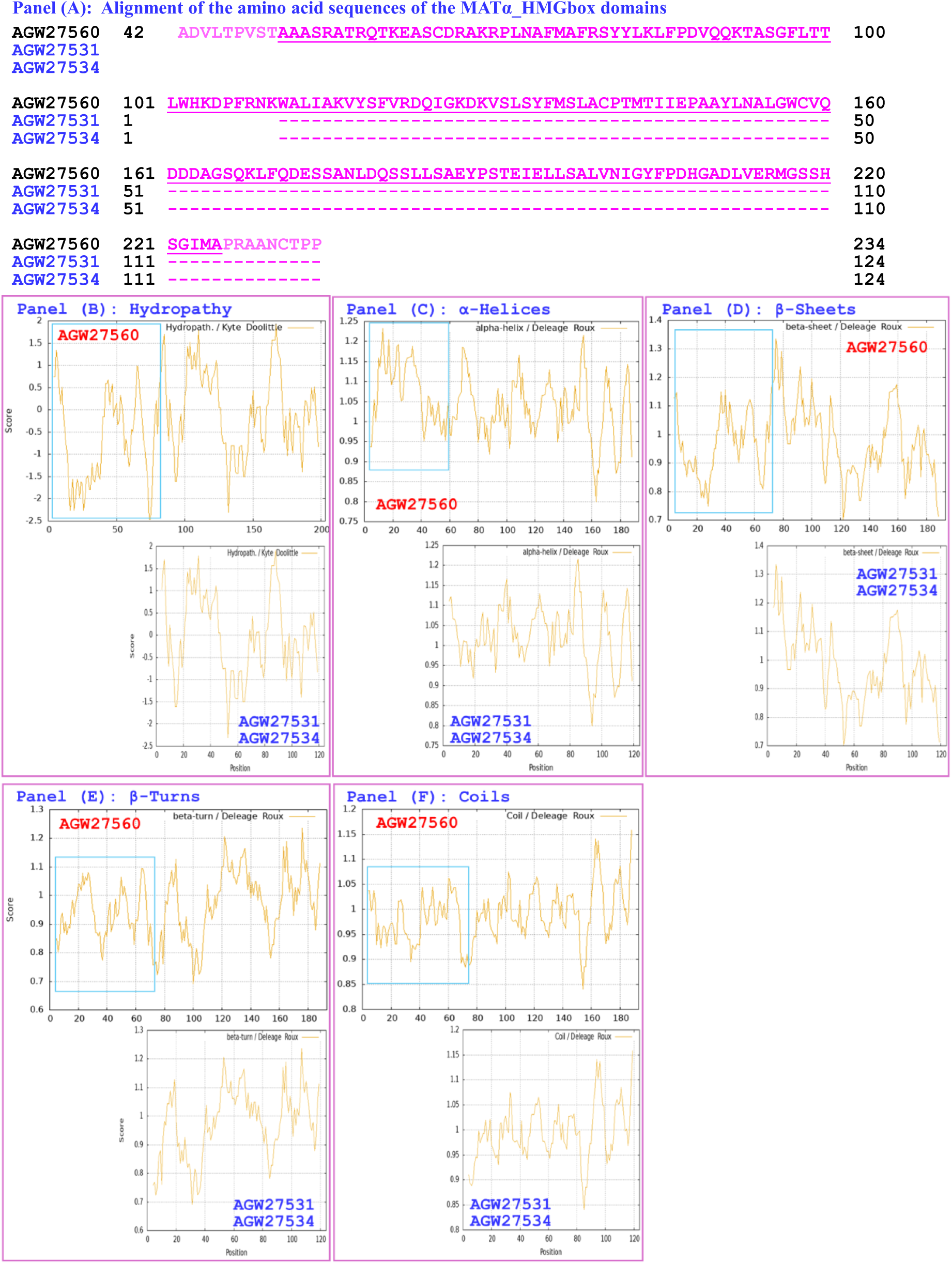
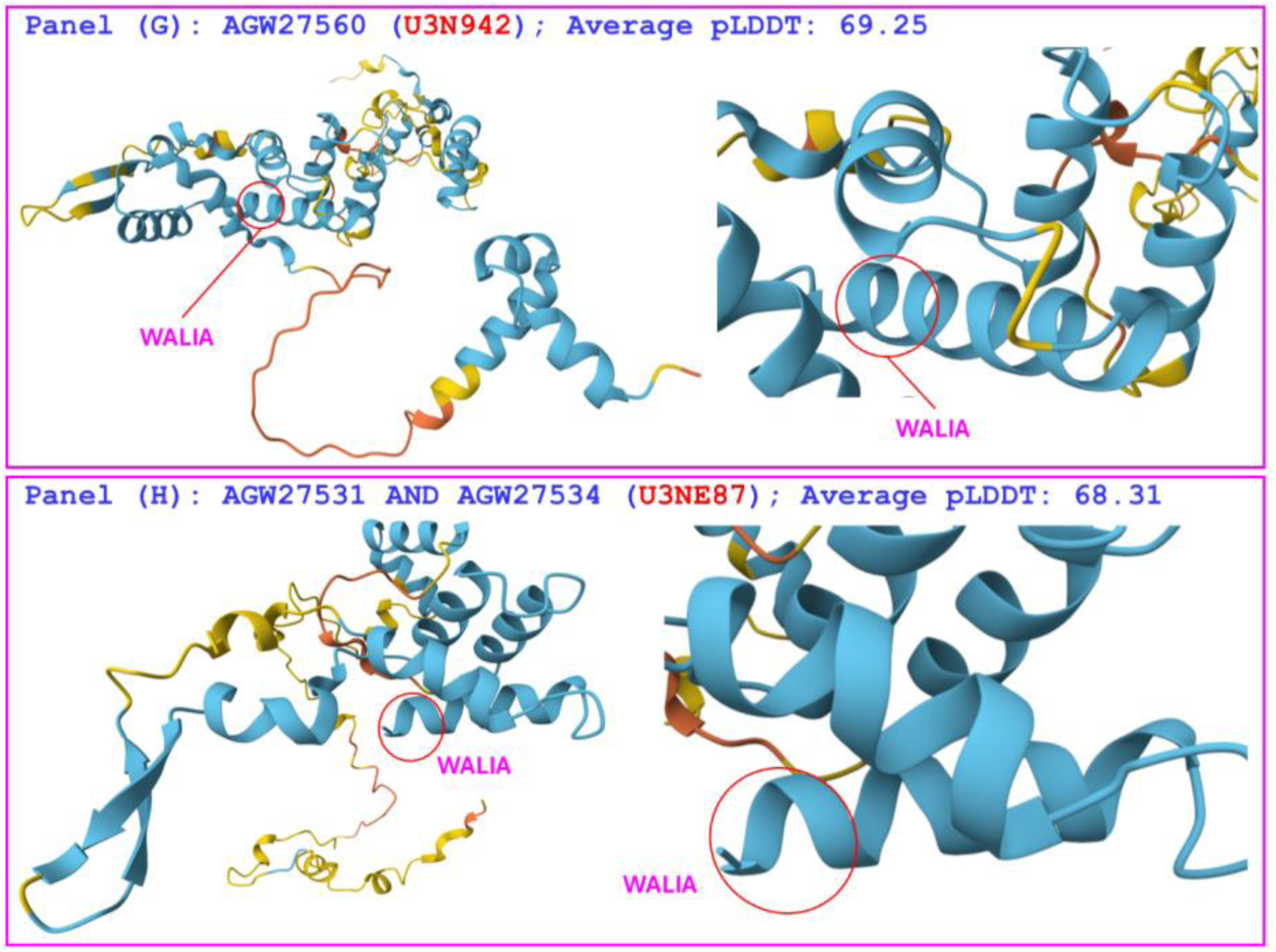
Correlations of the changes in the hydrophobicity and the primary, secondary, and tertiary structures of the MATα_HMGbox domains of MAT1-1-1 proteins: the reference protein AGW27560 (under the AlphaFold code U3N942) derived from the H. sinensis strain CS68-2-1229 and the variant proteins AGW27531 and AGW27534 (under the AlphaFold code U3NE87) derived from the O. sinensis strains CS70-1212 and CS71-1219, respectively. Panel (A) shows an alignment of the amino acid sequences of the MATα_HMGbox domains of the MAT1-1-1 proteins, where the hyphens indicate identical amino acid residues and the spaces denote unmatched sequence gaps. The ExPASy ProtScale plots show the changes in hydrophobicity and the 2D structure in Panels (B)–(F) for hydropathy, α-helices, β-sheets, β-turns, and coils of the protein; the open rectangles in blue highlight the N-terminally truncated regions in the ExPASy plots. Panels (G)–(H) show the 3D structures of the full-length proteins on the left, and the locally magnified structures surrounding the truncation site are shown on the right. The model confidence for the AlphaFold-predicted 3D structures is as follows: 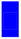 very high (pLDDT>90); 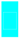 high (90>pLDDT>70); 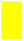 low (70>pLDDT>50); and 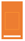 very low (pLDDT<50).

Collectively, the data in Figures 5–12 in Section 3.3 demonstrate that the MAT1-1-1 protein variants derived from various *O. sinensis* strains exhibit N-terminal truncations (ranging from 13 to 60 residues) of their MATα_HMGbox domains, with different amino acid substitutions or deletions. These modifications significantly alter the hydrophobicity, secondary structures, and tertiary structures of the MATα_HMGbox domains, with truncation and substitution locations occurring either upstream of or within the three core α-helices of the domains. The variant MAT1-1-1 proteins correspond to one of nine distinct AlphaFold 3D structural conformations (*cf*. Table 1, Figure 2), highlighting the structural diversity of the MAT1-1-1 proteins.

### 3.4 Heteromorphic tertiary structures of the HMG-box_ROX1-like domains of the truncated MAT1-2-1 proteins derived from *O. sinensis* strains

This section presents the results of the correlation analyses between the changes in the hydrophobic properties and structures of the HMG-box_ROX1-like domains of variant MAT1-2-1 proteins derived from various *O. sinensis* strains under 4 AlphaFold codes (*cf*. Table 1) compared with the reference full-length MAT1-2-1 proteins. This reference protein, represented by AEH27625 derived from the *H. sinensis* strain CS2 (*cf*. Table S5), is associated with the AlphaFold 3D structural code D7F2E9 [70]. The HMG-box_ROX1-like domain sequences (amino acid residues 127→197 of the authentic protein AEH27625) were extended both upstream and downstream by 9 external amino acids, which was used in the sequential ExPASy ProtScale plot for hydropathy, α-helices, β-sheets, β-turns, and coils with a window size of 9 amino acid residues (*cf*. Section 2.3).

The hydrophobicity and structural characteristics of the HMG-box_ROX1-like domain of the MAT1-2-1 protein AGW27543 are compared with those of the reference protein AEH27625 in Figure 13. The protein AGW27543 (under the AlphaFold code U3N9W0) is derived from the *O. sinensis* strain CS91-1291 (*cf*. Table 1) [14], while the reference protein AEH27625 (under the AlphaFold code D7F2E9) is derived from the *H. sinensis* strain CS2 [70]. Panel (A) of Figure 13 shows a 12-residue truncation at the C-terminus of the HMG-box_ROX1-like domain of the protein AGW27543, along with a Y-to-H substitution (tyrosine to histidine). This substitution changes the hydropathy index from -1.3 to -3.2 (Table S3) [73], resulting in reduced hydrophobicity. The C-terminal truncation and amino acid substitution, along with the accompanying reduction in hydrophobicity, resulted in altered secondary structures surrounding the variation sites within the HMG-box_ROX1-like domain of the protein AGW27543. This alteration is reflected by changes in the topological structure and waveform of the ExPASy ProtScale plots, which illustrate hydropathy, α-helices, β-sheets, β-turns, and coils in Panels (B)−(F), respectively. As shown in the 3D structures presented in Panels (G)−(H), the truncation and replaced amino acid residues are located in 2 of the 3 α-helices that form the core L-shaped stereostructure. These modifications result in an altered tertiary structure of the HMG-box_ROX1-like domain of the protein AGW27543 under the AlphaFold code U3N9W0.

**Figure 13.**
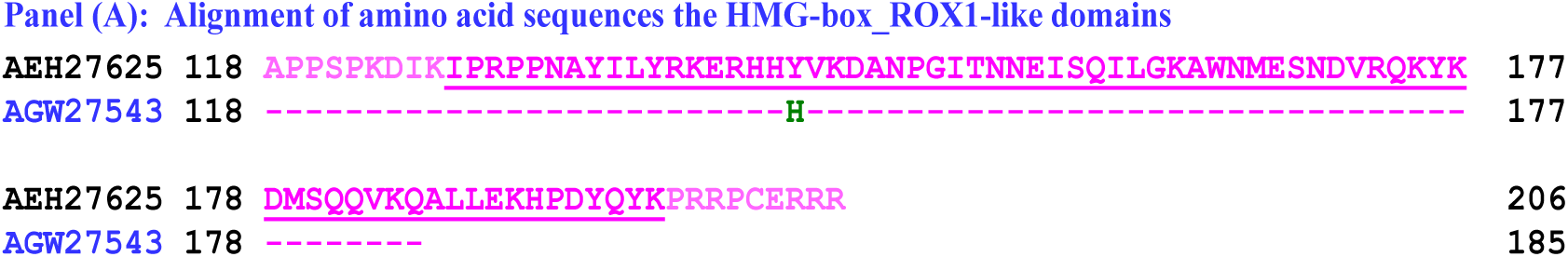
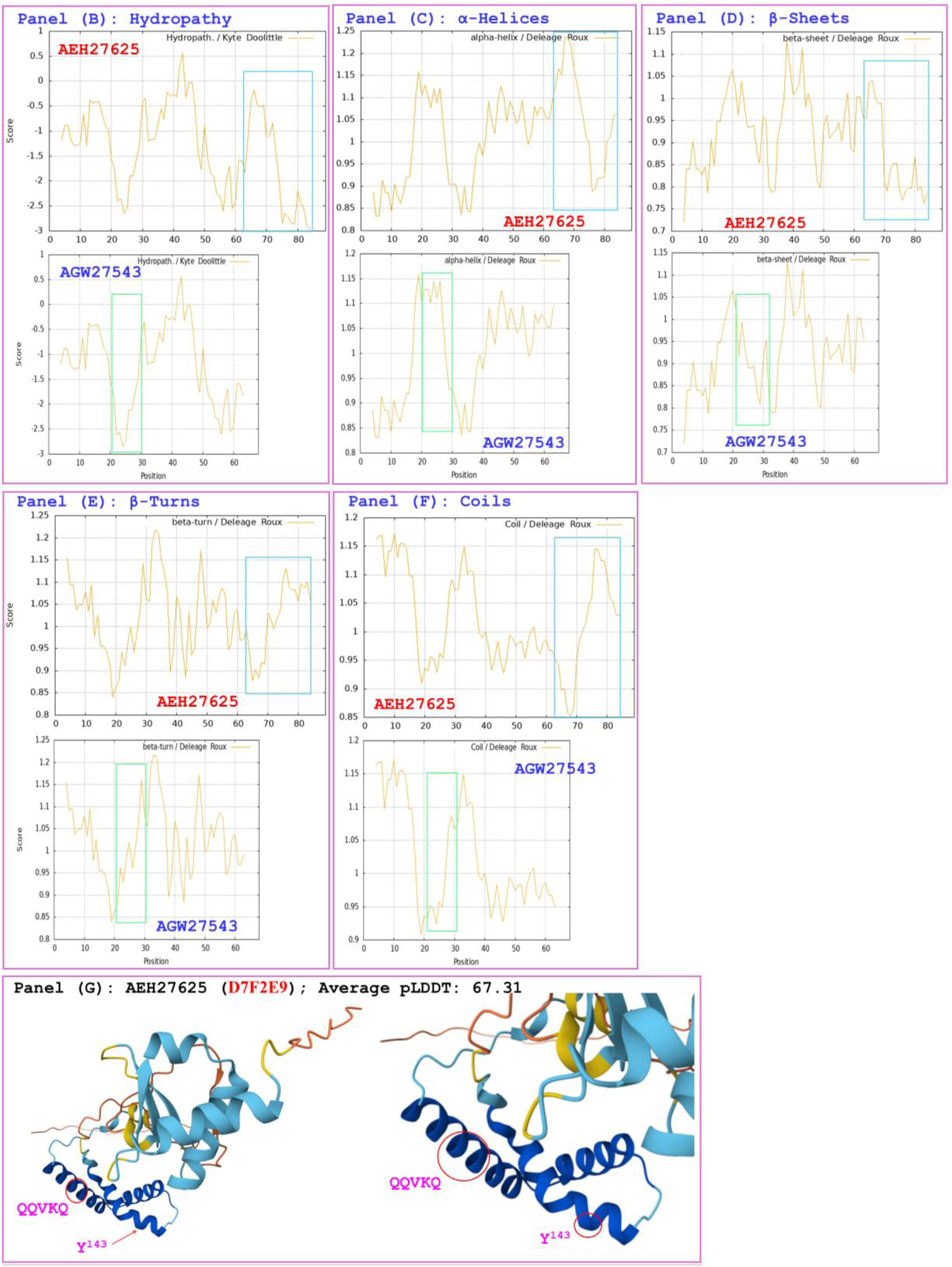
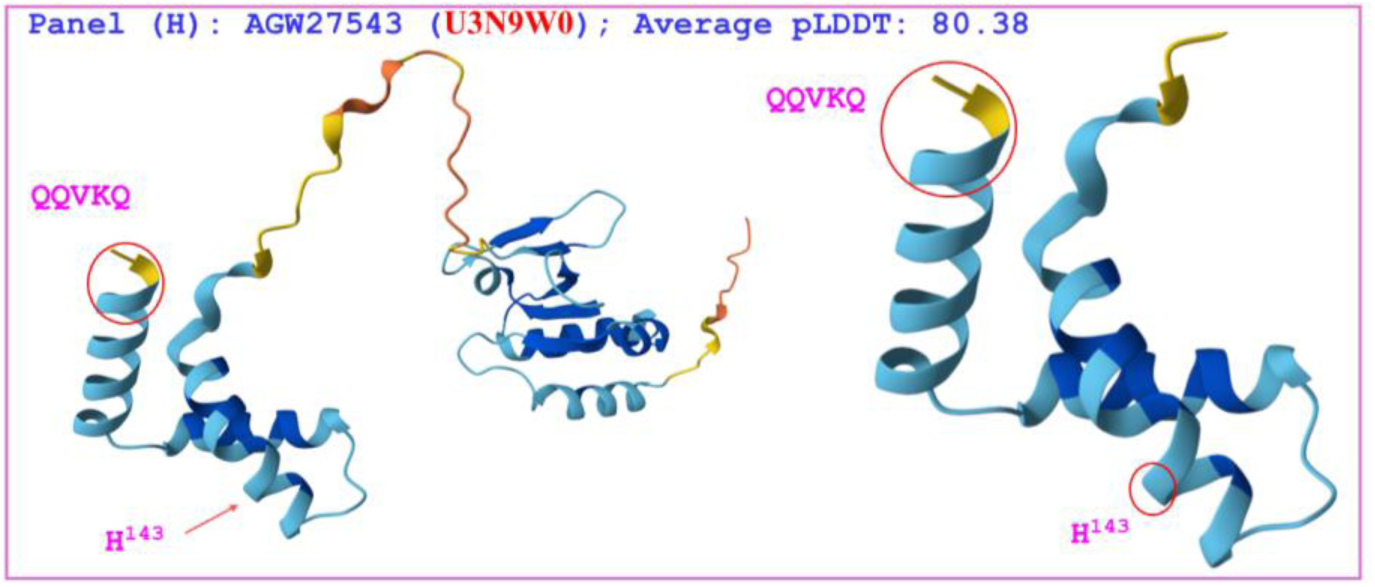
Correlations between the changes in hydrophobicity and the primary, secondary, and tertiary structures of the HMG-box_ROX1-like domains of MAT1-2-1 proteins. The reference protein AEH27625 (under the AlphaFold code D7F2E9) is derived from the H. sinensis strain CS2, and the variant MAT1-2-1 protein AGW27543 (under the AlphaFold code U3N9W0) is derived from the O. sinensis strain CS91-1291. Panel (A) shows an alignment of the amino acid sequences of the HMG-box_ROX1-like domains of the MAT1-2-1 proteins; the amino acid substitution is shown in green, the hyphens indicate identical amino acid residues, and the spaces denote unmatched sequence gaps. The ExPASy ProtScale plots show the changes in hydrophobicity and the 2D structure of the protein in Panels (B)–(F) for the hydropathy, α-helices, β-sheets, β-turns, and coils, respectively; the open rectangles in blue highlight the C-terminally truncated regions in the ExPASy plots, and those in green highlight the changes in topological structure and waveform observed in the plots. Panels (G) –(H) show representations of the 3D structures of the full-length proteins on the left; the locally magnified structures at the variation sites are shown on the right. The model confidence for the AlphaFold-predicted 3D structures is as follows: 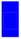 very high (pLDDT>90); 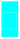 high (90>pLDDT>70); 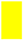 low (70>pLDDT>50); and 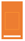 very low (pLDDT<50).

The hydrophobicity and structural characteristics of the HMG-box_ROX1-like domain of the MAT1-2-1 protein AGW27554 are compared with those of the reference protein AEH27625 in Figure 14. The protein AGW27554 (under the AlphaFold code U3NEA9) is derived from the *O. sinensis* strain CS71-1219 [14], whereas the reference protein AEH27625 (under the AlphaFold code D7F2E9) is derived from the *H. sinensis* strain CS2 [70]. Panel (A) of Figure 14 shows a 12-residue truncation at the C-terminus of the HMG-box_ROX1-like domain of the protein AGW27554, along with 2 amino acid substitutions: Y-to-H (tyrosine to histidine; hydropathy indices changed from -1.3 to -3.2) and Q-to-T (glutamine to threonine; hydropathy indices changed from -3.5 to -0.7 (Table S3) [73]. The C-terminal truncation and amino acid substitutions accompanied by altered hydrophobicity result in changes in the secondary structure surrounding the variation sites in the HMG-box_ROX1-like domain of the protein AGW27543. This alteration is reflected by the changes in the topological structure and waveform of the ExPASy ProtScale plots, which illustrate hydropathy, α-helices, β-sheets, β-turns, and coils, in Panels (B)−(F), respectively. As illustrated in the 3D structures in Panels (G)−(H), the truncated and replaced residues are located in 2 of the 3 core α-helices. These modifications result in an altered tertiary structure of the HMG-box_ROX1-like domain of the protein AGW27554 under the AlphaFold code U3NEA9.

**Figure 14.**
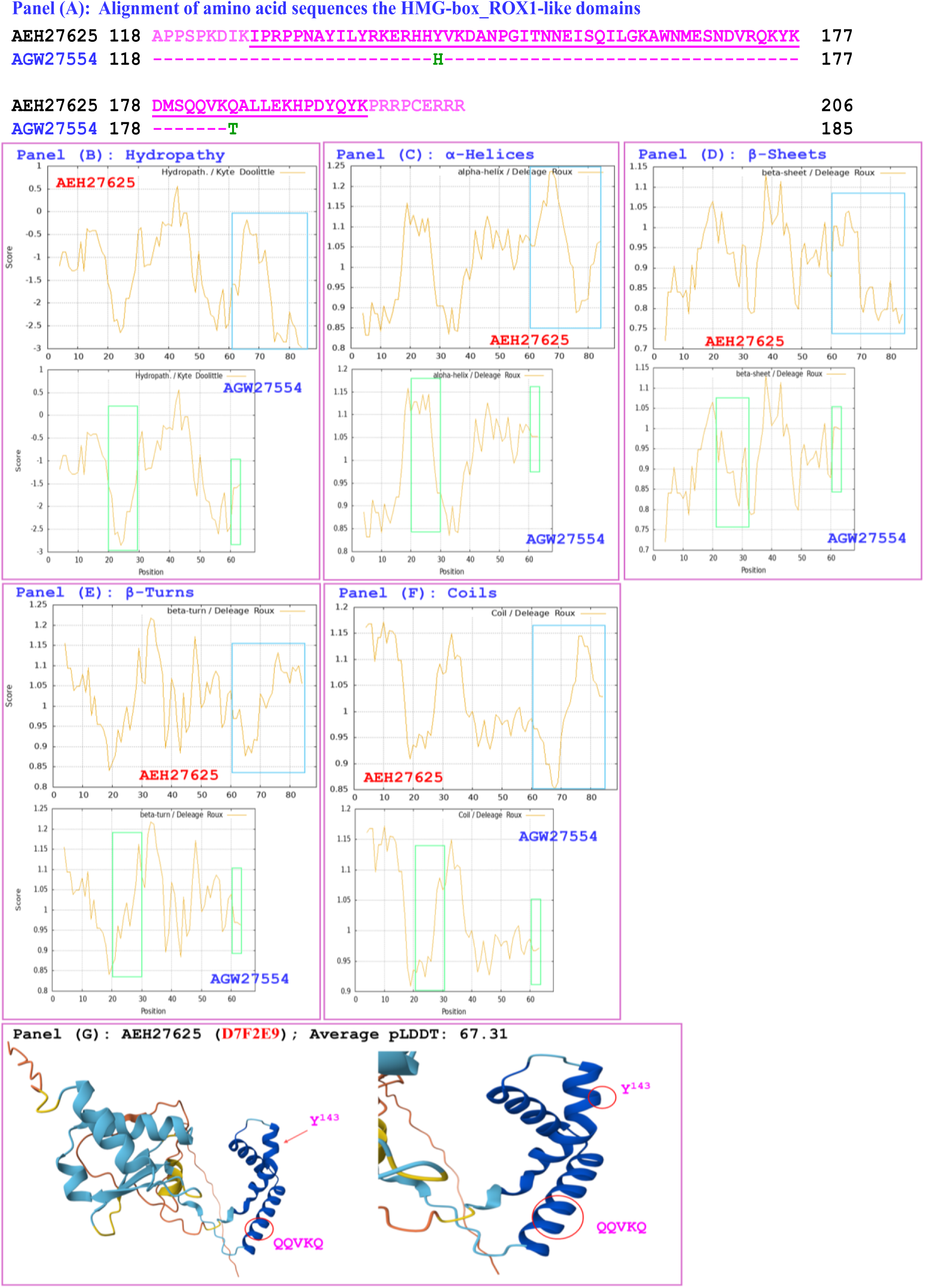
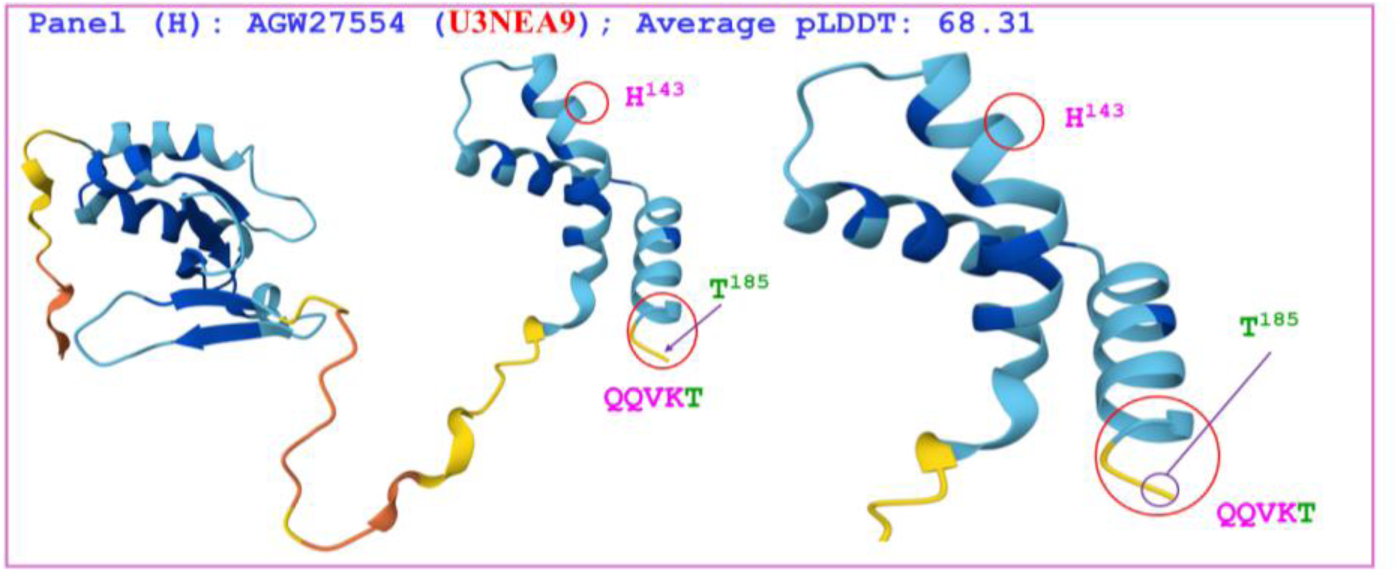
Correlations between changes in hydrophobicity and the primary, secondary, and tertiary structures of the HMG-box_ROX1-like domains of MAT1-2-1 proteins. The reference protein AEH27625 (under the AlphaFold code D7F2E9) is derived from the H. sinensis strain CS2, and the variant MAT1-2-1 protein AGW27554 (under the AlphaFold code U3NEA9) is derived from the O. sinensis strain CS71-1219. Panel (A) shows an alignment of the amino acid sequences of the HMG-box_ROX1-like domainsof the MAT1-2-1 proteins; amino acid substitutions are shown in green, the hyphens indicate identical amino acid residues, and the spaces denote unmatched sequence gaps. The ExPASy ProtScale plots show the changes in hydrophobicity and the 2D structure of the protein in Panels (B)–(F) for hydropathy, α-helices, β-sheets, β-turns, and coils, respectively; the open rectangles in blue highlight the C-terminal truncation regions in the ExPASy plots, and those in green highlight the changes in the topological structure and waveform. Panels (G)–(H) show representations of the 3D structures of the full-length proteins on the left; the locally magnified structures at the variation sites are shown on the right. The model confidence for the AlphaFold-predicted 3D structures is as follows: 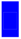 very high (pLDDT>90); 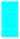 high (90>pLDDT>70); 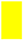 low (70>pLDDT>50); and 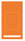 very low (pLDDT<50).

The hydrophobicity and structural characteristics of the HMG-box_ROX1-like domains of the MAT1-2-1 proteins AGW27548 and AGW27555 are compared with those of the reference protein AEH27625, as shown in Figure 15. These 2 variant proteins (both under the AlphaFold code U3N9W5) are derived from the *O. sinensis* strains CS68-2-1229 and CS71-1220, respectively (*cf*. Table 1) [14], while the reference protein AEH27625 (under the AlphaFold code D7F2E9) is derived from the *H. sinensis* strain CS2 [70]. Panel (A) of Figure 15 shows 22-residue truncations at the C-termini of the HMG-box_ROX1-like domains of the MAT1-2-1 proteins AGW27548 and AGW27555, along with a Y-to-H (tyrosine to histidine) amino acid substitution within the HMG-box_ROX1-like domains of both variant proteins. This substitution changes the hydropathy index from -1.3 to -3.2 (Table S3) [73]. The C-terminal truncations and amino acid substitutions caused decreased hydrophobicity and altered the secondary structures surrounding the variation sites in the HMG-box_ROX1-like domains of the MAT1-2-1 proteins AGW27548 and AGW27555. This alteration is reflected by changes in the topological structure and waveform of the ExPASy ProtScale plots, which illustrate hydropathy, α-helices, β-sheets, β-turns, and coils in Panels (B)−(F), respectively. As shown in the 3D structures in Panels (G)−(H), the truncations and replaced amino acid residues are located in 2 of the 3 core α-helices. These modifications result in altered tertiary structures of the HMG-box_ROX1-like domains of the MAT1-2-1 proteins

**Figure 15.**
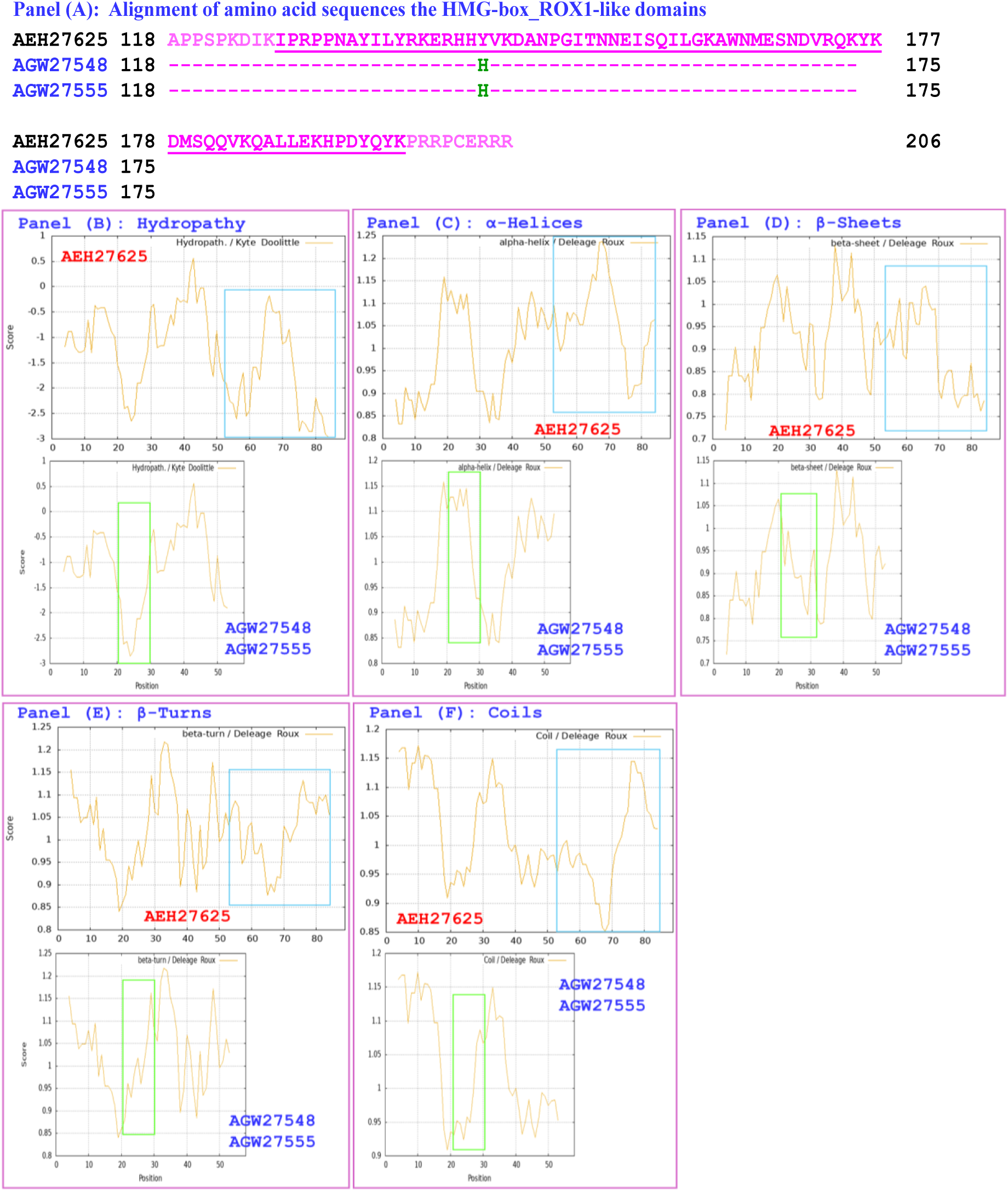
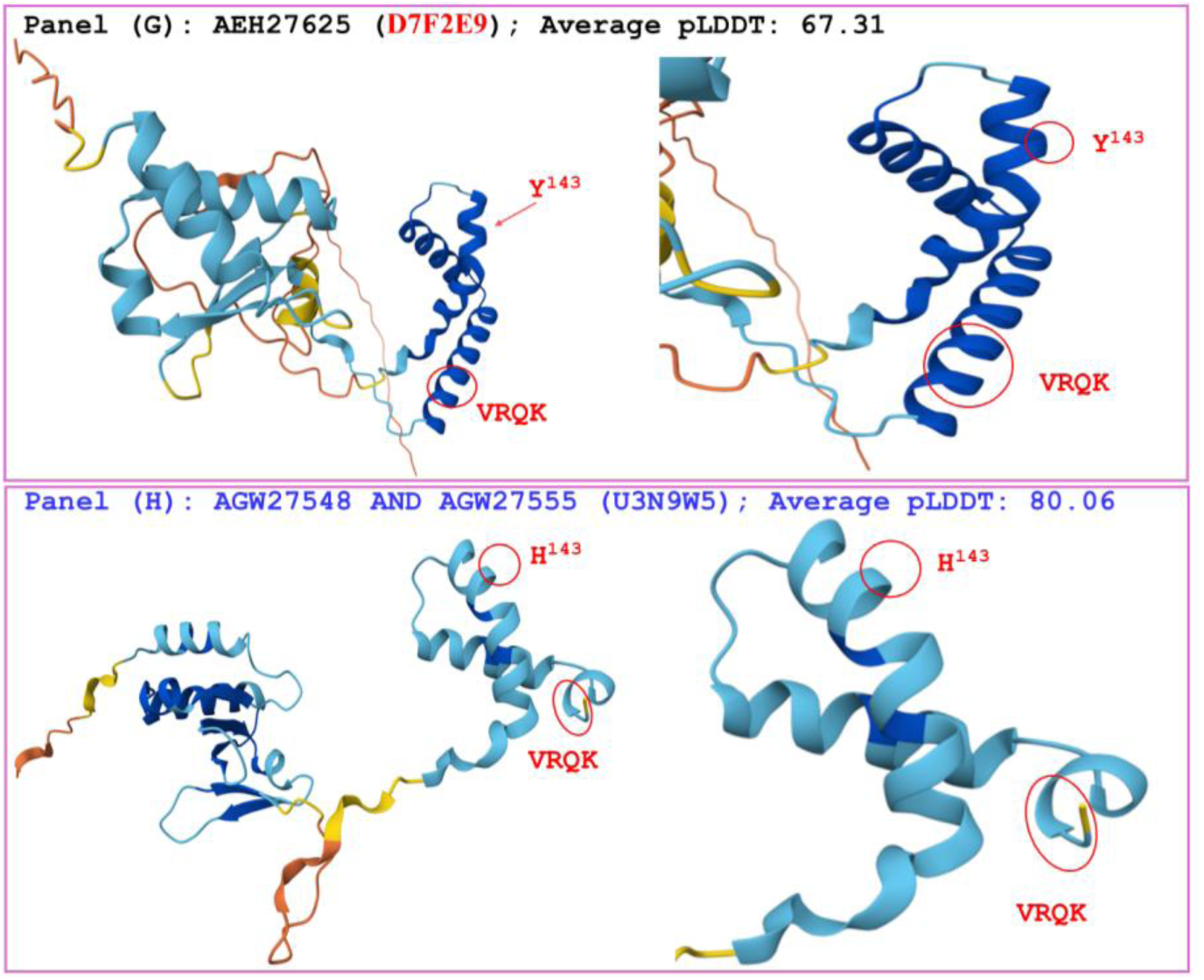
Correlations between changes in hydrophobicity and the primary, secondary, and tertiary structures of the HMG-box_ROX1-like domains of MAT1-2-1 proteins. The reference protein AEH27625 (under the AlphaFold code D7F2E9) is derived from the H. sinensis strain CS2, and the variant MAT1-2-1 proteins AGW27548 and AGW27555 (under the AlphaFold code U3N9W5) are derived from the O. sinensis strains CS68-2-1229 and CS71-1220, respectively. Panel (A) shows an alignment of the amino acid sequences of the HMG-box_ROX1-like domains of the MAT1-2-1 proteins; amino acid substitutions are shown in green, the hyphens indicate identical amino acid residues, and the spaces denote unmatched sequence gaps. The ExPASy ProtScale plots show the changes in hydrophobicity and the 2D structure in Panels (B)–(F) for the hydropathy, α-helices, β-sheets, β-turns, and coils of the proteins, respectively; the open rectangles in blue highlight the C-terminally truncated regions in the ExPASy plots, and the changes in topological structure and waveform observed in the plots are highlighted in red. Panels (G)–(H) show representations of the 3D structures of the full-length proteins on the left; the locally magnified structures at the variation sites are shown on the right. The model confidence for the AlphaFold-predicted 3D structures is as follows: 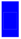 very high (pLDDT>90); 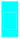 high (90>pLDDT>70); l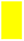 low (70>pLDDT>50); and 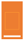 very low (pLDDT<50).

The hydrophobicity and structural characteristics of the HMG-box_ROX1-like domain of the MAT1-2-1 protein AGW27552 are compared with those of the reference protein AEH27625, as shown in Figure 16. The variant protein (under the AlphaFold code U3N6W6) is derived from the *O. sinensis* strain CS68-5-1216 [14], while the reference protein AEH27625 (under the AlphaFold code D7F2E9) is derived from the *H. sinensis* strain CS2 [70]. Panel (A) of Figure 16 shows a 23-residue truncation at the C-terminus of the HMG-box_ROX1-like domain in the MAT1-2-1 protein AGW27552, along with a Y-to-H substitution (tyrosine to histidine; hydropathy indices changed from -1.3 to -3.2) (Table S3) [73]) within the HMG-box_ROX1-like domain. This amino acid substitution caused decreased hydrophobicity and modified the secondary structures surrounding the sites of variation within the HMG-box_ROX1- like domain of the MAT1-2-1 protein AGW27552. Changes in the topological structure and waveform are reflected in the ExPASy ProtScale plots corresponding to hydropathy, α-helices, β-sheets, β-turns, and coils in Panels (B)–(F), respectively. As shown in the comparison of the 3D structures in Panels (G)– (H) for a comparison of 3D structures, the truncation and replaced residue occur within 2 of the 3 core α-helices and completely remodel the overall stereostructure of the HMG-box_ROX1-like domain of the MAT1-2-1 protein AGW27552 (under AlphaFold code U3N6W6).

**Figure 16.**
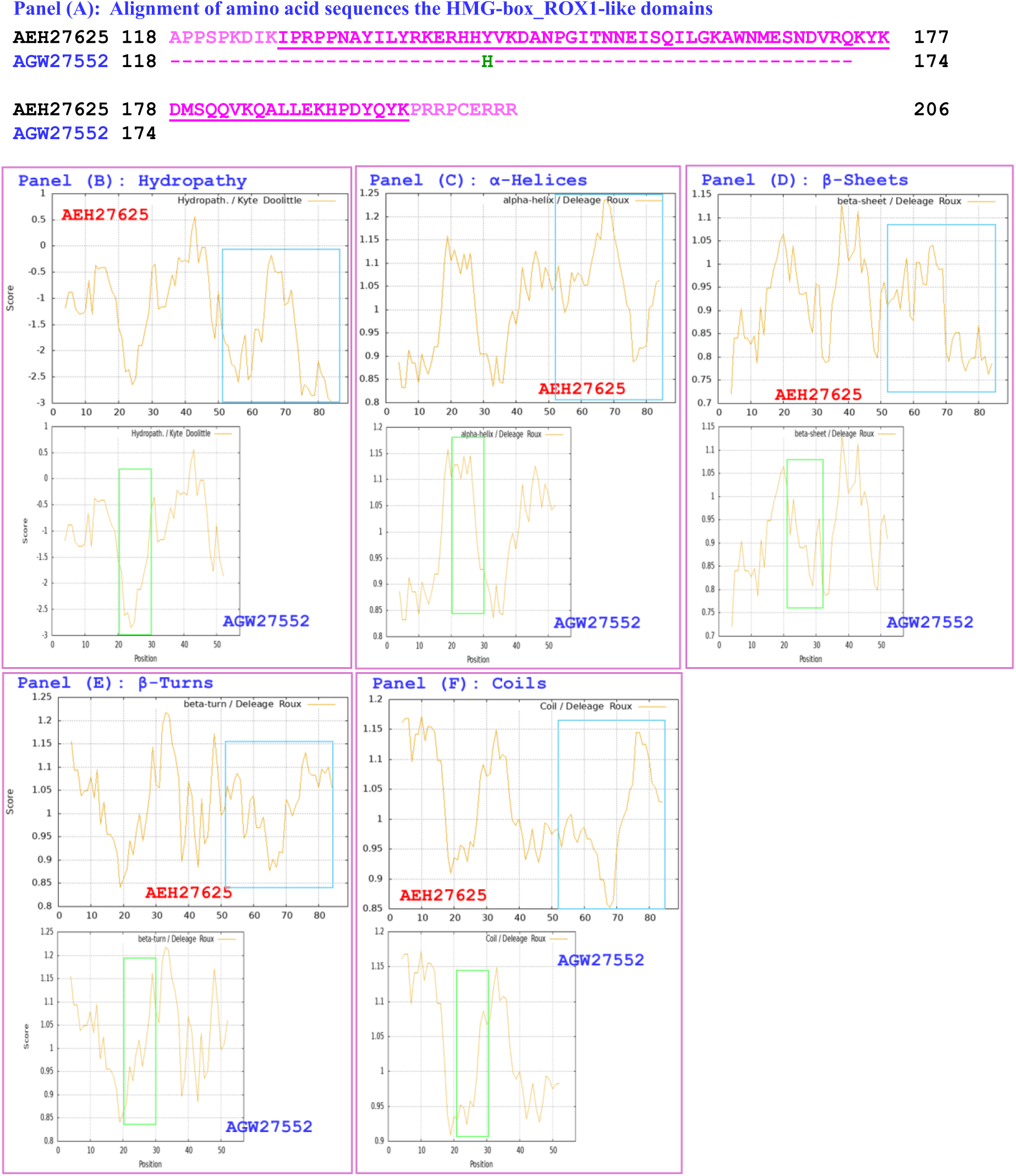
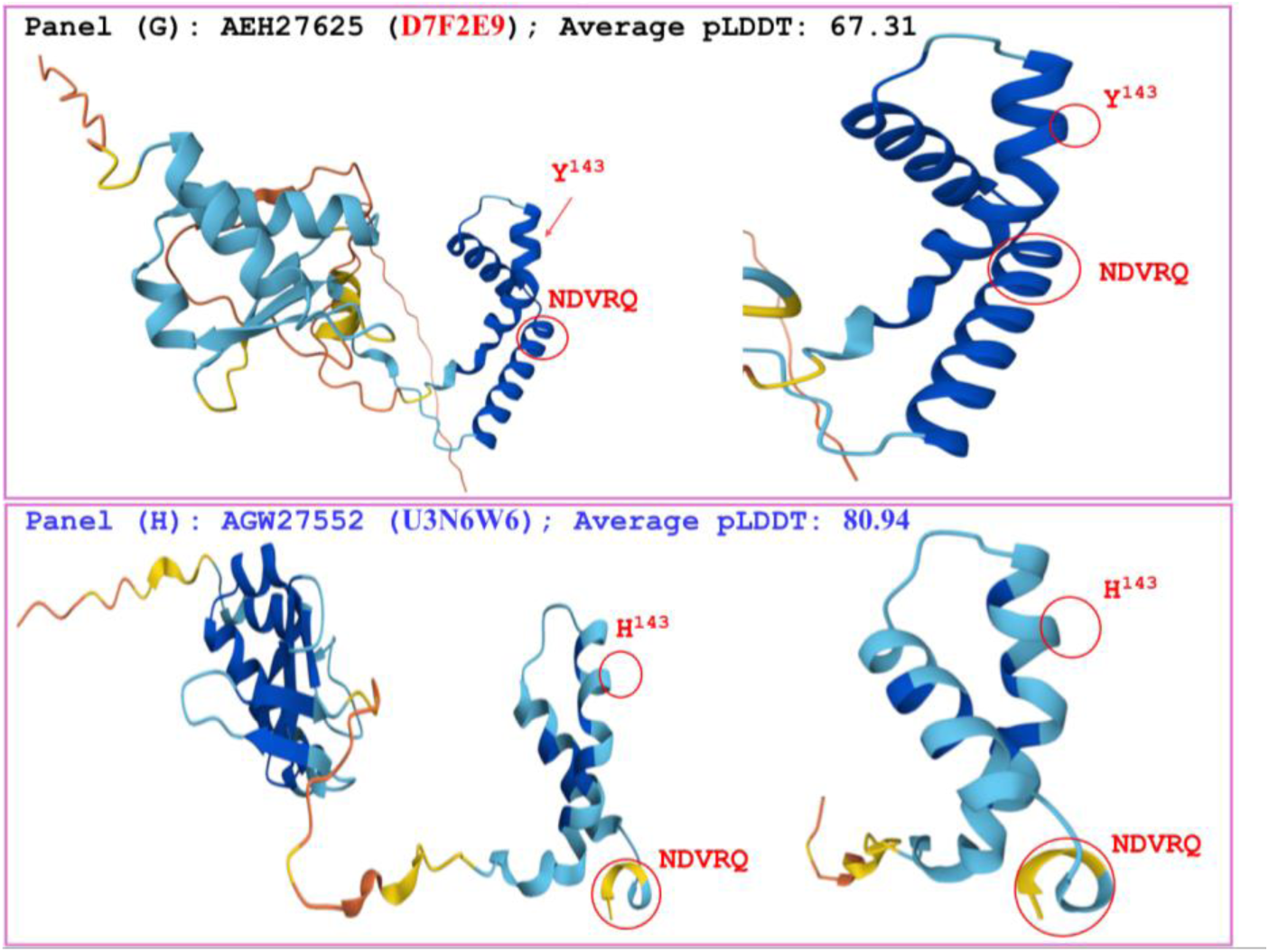
Correlations between changes in hydrophobicity and the primary, secondary, and tertiary structures of the HMG-box_ROX1-like domains of MAT1-2-1 proteins. The reference protein AEH27625 (under the AlphaFold code D7F2E9) is derived from the H. sinensis strain CS2, and the variant MAT1-2-1 protein AGW27552 (under the AlphaFold code U3N6W6) is derived from the O. sinensis strain CS68-5-1216. Panel (A) shows an alignment of the amino acid sequences of the HMG-box_ROX1-like domains of the MAT1-2-1 proteins; amino acid substitutions are shown in green, the hyphens indicate identical amino acid residues, and the spaces denote unmatched sequence gaps. The ExPASy ProtScale plots show the changes in hydrophobicity and the 2D structure of the protein in Panels (B)–(F) for the hydropathy, α-helices, β-sheets, β-turns, and coils, respectively; the open rectangles in blue highlight the C-terminally truncated regions in the ExPASy plots, and those in red highlight the changes in topological structure and waveform observed in the plots. Panels (G) –(H) show representations of the 3D structures of the full-length proteins on the left; the locally magnified structures at the sites of substitutions are shown on the right. The model confidence for the AlphaFold-predicted 3D structures is as follows: 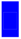 very high (pLDDT>90); 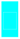 high (90>pLDDT>70); 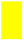 low (70>pLDDT>50); and 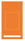 very low (pLDDT<50).

Collectively, the data in Figures 13–16 in Section 3.4 demonstrate that variant MAT1-2-1 proteins derived from various *O. sinensis* strains exhibit C-terminal truncations (ranging from 12 to 23 residues) within their HMG-box_ROX1-like domains, accompanied by tyrosine-to-histidine (Y-to-H) substitutions. These structural modifications alter the hydrophobicity, secondary structures, and tertiary structures of the HMG-box_ROX1-like domains. Both truncations and substitutions are localized within the 3 core α-helices of the domains. The variant MAT1-2-1 proteins correspond to one of four distinct AlphaFold 3D structural conformations (*cf*. Table 1, Figure 4), highlighting the structural diversity of this protein family among different *O. sinensis* strains.

### 3.5 Diverse fungal sources of the MAT1-1-1 and MAT1-2-1 protein variants produced simultaneously in pairs by the 20 *O. sinensis* strains

#### 3.5a Complex genetic heterogeneity of the *O. sinensis* strains

The ITS sequences used for phylogenetic analysis are available in GenBank for 15 of the 20 *O. sinensis* strains obtained by Li *et al*. [36] but are lacking for the remaining 5 strains (*cf*. Table 1). The design of the PCR experiment in that study included the use of a pair of fungal universal primers (ITS5/ITS4 [92]) and 3 pairs of self-designed primers (GAF/GAR, GCF/GCR, and 5.8S-F/R) for ITS sequence amplification, and the annealing temperatures were set to 53°C for primers ITS5/ITS4, 55°C for primers GAF/GAR specific for the GC-biased sequences of Group A (Genotype #1 of *O. sinensis*), and 60°C for primers GCF/GCR specific for the AT-biased sequences of group C (Genotype #5 of *O. sinensis*). Table S6 lists the percentage identities and sequence coverages between the sequences of the primers and the representative GC- and AT-biased genotypes of *O. sinensis*. Table S6 shows that many genotypes exhibit no or significantly low sequence identities and coverages, indicating that they may not be amplified if they truly co-occur in the test samples. Compared with other sequences, some genotypic components of *O. sinensis* might be much more efficiently amplified, causing differential dominances of the PCR amplicons. Furthermore, Li *et al*. [36] adopted a direct sequencing strategy for amplicons. Obviously, this study did not employ rigorous amplicon cloning–sequencing techniques with the selection of a sufficient number (30−50) of white colonies to achieve comprehensive, unbiased genetic profiling, nor did it use the MassARRAY single-nucleotide polymorphism (SNP) MALDI‒TOF mass spectrum genotyping technique with multiple extension primers for comprehensive phylogenetic examinations. Consequently, molecular examination after 25 days of shaker incubation of *C. sinensis* monoascospores and tissues of caterpillar bodies (sclerotium) would have inevitably overlooked nonculturable fungal sequences and some co-occurring amplicon sequences whose abundance in the amplicon pools might have been low because of low amplification efficiency under the design used in this study [36,37].

Under these experimental settings, Li *et al*. [36] reported that 10 of the 15 *O. sinensis* strains are homogenous, harboring only GC-biased Genotype #1 *H. sinensis* (*cf*. Table 1); the remaining 5 strains are heterogeneous, containing both GC-biased Genotype #1 (Group A) and AT-biased Genotype #17 (Group C) (*cf*. Tables S1−S2) [9,10,12,13,36,37]. However, because of the insufficiently refined research design adopted by Li *et al*. [36], a high level of vigilance is warranted regarding the potential coexistence of additional fungal species or *O. sinensis* genotypes in previously purportedly homogenous and heterogeneous strains. Given the complex genetic background of the *O. sinensis* strains, the possibility cannot be excluded that the detected MAT1-1-1 and/or MAT1-2-1 protein variants, which are characterized by differential truncations and various amino acid substitutions at distinct variation sites, originate from other unnoticed coexisting fungi.

Notably, the remaining 5 *O. sinensis* strains (CS26-277, CS36-1294, CS68-5-1216, CS70-1211, and CS70-1212, shown in black in Table 1) lack ITS sequence records in the GenBank database, and their homo/heterogeneous characteristics remain unknown. Researchers who possess these strains [14,36] are expected to upload the missing ITS sequences to GenBank, ensuring the comprehensive and impartial molecular documentation of these strains.

#### 3.5b Differential pairing of the MAT1-1-1 and MAT1-2-1 proteins simultaneously produced by the 20 *O. sinensis* strains

Sections 3.1−3.4 present various AlphaFold tertiary structural models of the mating protein variants derived from 20 *O. sinensis* strains. These strains simultaneously produced paired MAT1-1-1 and MAT1-2-1 proteins that harbor differential truncations at the N- and/or C-termini and diverse amino acid substitutions. As shown in Table 2, 10 of the 20 strains are “homogenous” strains shown in brown, 5 are heterogeneous strains shown in red, and the remaining 5 strains have unknown homo/heterogeneous characteristics and are indicated in black. On the basis of the pairing patterns of the 2 mating proteins, these strains can be categorized into 2 groups (Table 2). **Group I** consists of 5 strains, which simultaneously produce truncated MAT1-1-1 and MAT1-2-1 proteins in pairs; **Group II** is composed of the remaining 15 *O. sinensis* strains, which simultaneously produce the truncated MAT1-1-1 proteins and the full-length MAT1-2-1 proteins in pairs (*cf*. Table S5). The MAT1-1-1 and MAT1-2-1 proteins that share the same AlphaFold 3D structural morph are labeled with the same suffix symbol (⁋, †, *, ⁑, ‡, or ♦) in Table 2.

**Table 2.**
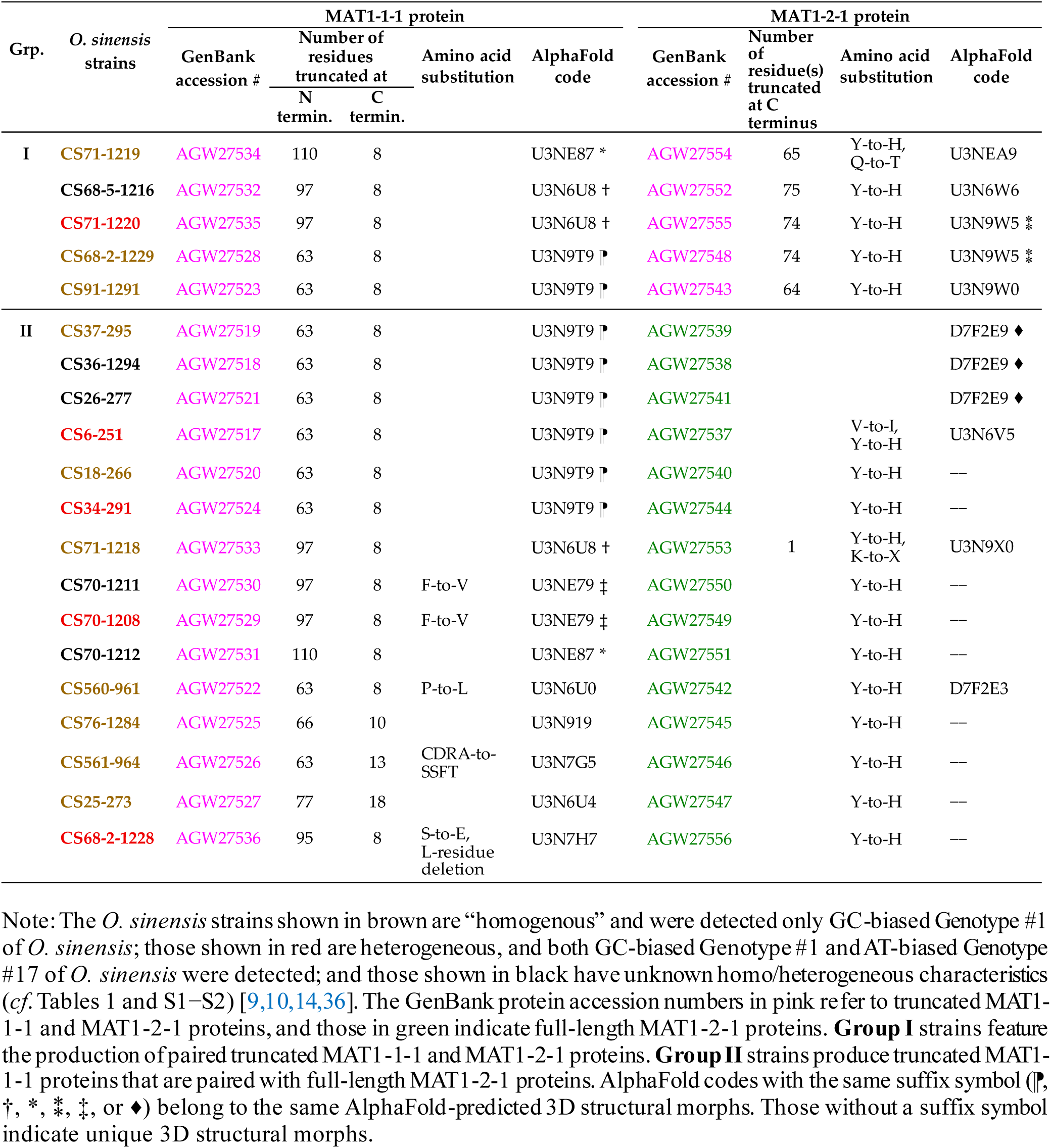
Two groups of the 20 *O. sinensis* strains based on the differential co-occurrence of pairs of MAT1-1-1 and MAT1-2-1 proteins with different truncations and amino acid substitutions

The 15 **Group-II** *O. sinensis* strains produced the full-length MAT1-2-1 proteins shown in green in Table 2. Three (AGW27538, AGW27539, and AGW27541) of them are 100% identical to the authentic MAT1-2-1 proteins represented by protein AEH27625 and share the same tertiary structures under the AlphaFold code D7F2E9. The remaining 12 MAT1-2-1 proteins (AGW27537, AGW27540, AGW27542, AGW27544−AGW27547, AGW27549−AGW27551, AGW27553, and AGW27556) contain 1−2 amino acid substitutions (Figure S3, Tables 2 and S5), all of which contain Y-to-H substitutions in the HMG-box_ROX1-like domains. The MAT1-2-1 protein AGW27537 derived from the *O. sinensis* strain CS6-251 contains an additional V-to-I (valine-to-leucine) substitution located upstream of the HMG-box_ROX1-like domain. The protein AGW27553 derived from the *O. sinensis* strain CS71-1218 contains an additional K-to-X (lysine-to-an unidentified residue) substitution located within the HMG-box_ROX1-like domain. Six of the 12 MAT1-2-1 proteins exhibit diverse tertiary structures under distinct AlphaFold 3D structural codes, whereas the remaining 6 proteins have not been assigned AlphaFold-predicted 3D structural morphs (Tables 2 and S5).

#### 3.5c *O. sinensis* strains simultaneously produced MAT1-1-1 proteins with the same tertiary structures and MAT1-2-1 proteins with various tertiary structures

The sequence distributions of the paired MAT1-1-1 and MAT1-2-1 proteins that were simultaneously produced by each of the 20 *O. sinensis* strains are shown in Figure 17.

**Figure 17.**
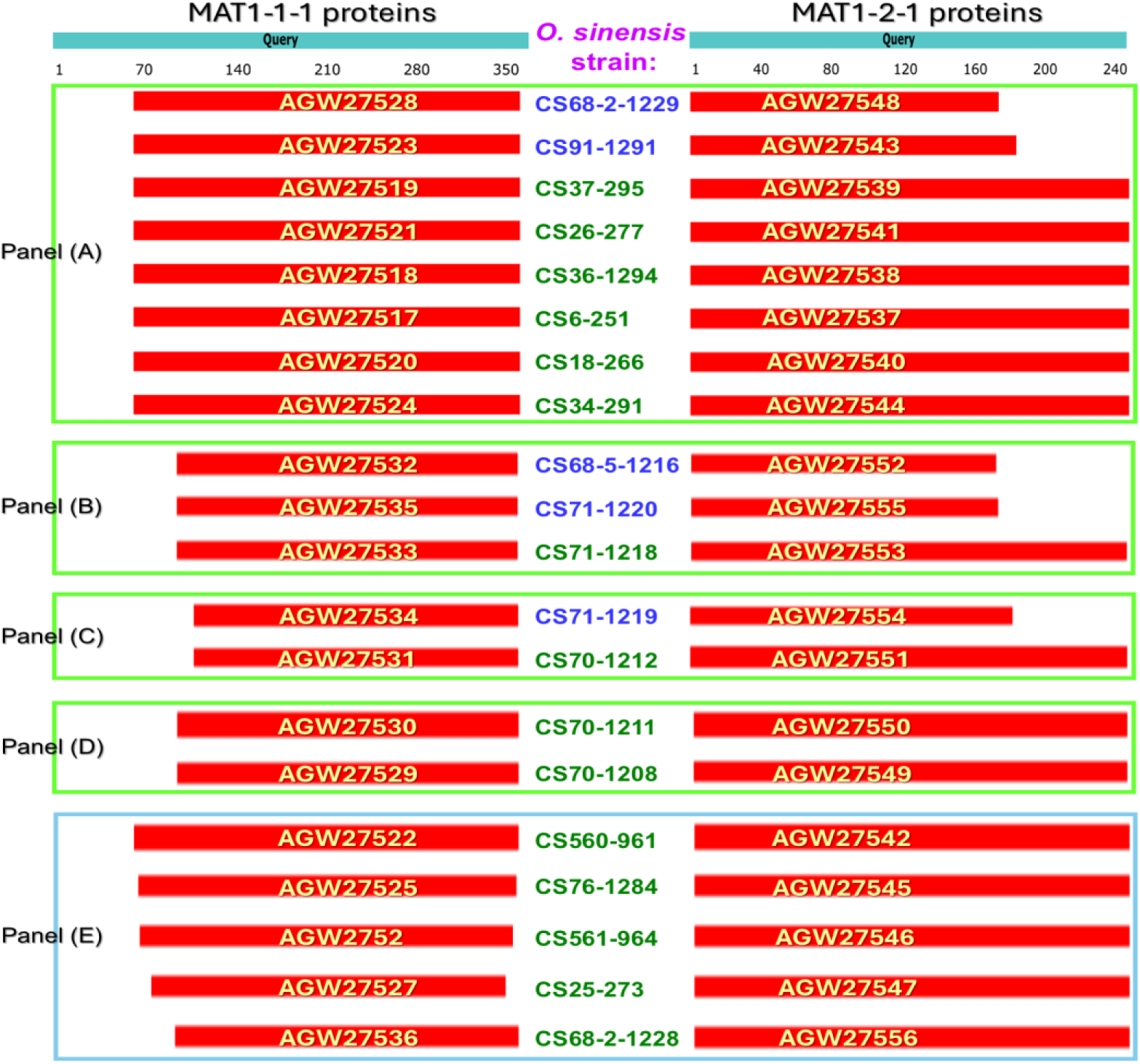
Alignment and sequence distributions of the paired MAT1-1-1 and MAT1-2-1 proteins derived from the O. sinensis strains in Group I shown in blue, and those in Group II shown in green (cf. Table 2). The MAT1-1-1 protein AGW27560 (AlphaFold code U3N942) derived from the H. sinensis strain CS68-2-1229 was used as the query MAT1-1-1 sequence, and the MAT1-2-1 protein AEH27625 (AlphaFold code D7F2E9) derived from the H. sinensis strain CS2 was used as the query MAT1-2-1 sequence in the alignment. The panels outlined with open rectangles in green and blue are arranged according to the AlphaFold 3D structural models for the MAT1-1-1 and MAT1-2-1 proteins, respectively.

Panel (A) of Figure 17 includes 8 *O. sinensis* strains: four purportedly homogenous strains are shown in brown in Table 2, two heterogeneous strains are shown in red, and the remaining two strains are shown in black with unknown homo/heterogeneous characteristics. All these strains produced MAT1-1-1 proteins truncated by 63- and 8-residues at the N- and C-termini, respectively, which exhibit the same tertiary structures under the AlphaFold 3D structural code U3N9T9 labeled with the suffix symbol “⁋” in Table 2 and analyzed above in Figure 5. Two purportedly homogenous strains, CS68-2-1229 and CS91-1291, belong to **Group I**, as shown in blue in Figure 17; these strains simultaneously produce the MAT1-2-1 proteins AGW27548 and AGW27543, which are differentially truncated by 74- and 64-residues, respectively, at the C-termini and contain Y-to-H (tyrosine-to-histidine) substitutions within the HMG-box_ROX1-like domains (Table 2; Figures 13, 15, and 17), resulting in altered 3D structures under AlphaFold codes U3N9W5 and U3N9W0, respectively.

The remaining 6 *O. sinensis* strains in Panel (A) belong to **Group II**, as shown in green in Figure 17, including 2 purportedly homogenous strains, 2 heterogeneous strains, and 2 additional strains with unknown homo/heterogeneous characteristics. In addition to the production of the same truncated MAT1-1-1 proteins (under the AlphaFold code U3N9T9, labeled with the suffix symbol “⁋” in Table 2), these 6 strains simultaneously produced full-length MAT1-2-1 proteins with heteromorphic 3D structures, which correspond to different AlphaFold codes or lack AlphaFold 3D structure codes and can be subgrouped as described below (*cf*. Table 2 and Panel (A) of Figure 17):

1. The coproduced full-length MAT1-2-1 proteins AGW27539, AGW27541, and AGW27538 exhibit the same 3D structure under the AlphaFold 3D structural code **D7F2E9**, which is marked with the suffix symbol “♦” in Table 2. These proteins are 100% identical to the authentic MAT1-2-1 protein AEH27625 (*cf*. Figure S3, Table S5) and were generated by the “homogenous” *O. sinensis* strain CS37-295, as well as the strains CS26-277 and CS36-1294 that lack ITS sequence records in GenBank (*cf*. Tables 1−2, Panel (A) of Figure 17).
2. The coproduced full-length MAT1-2-1 protein AGW27537 has an altered tertiary structure under the AlphaFold 3D structural code **U3N6V5** (*cf*. Figure 12 of [65] and Table S5) and was produced by the heterogeneous *O. sinensis* strain CS6-251 (*cf*. Tables 1−2, Panel (A) of Figure 17). This protein is 99.2% similar to the authentic MAT1-2-1 protein AEH27625 and contains V-to-I and Y-to-H substitutions, which are located upstream of and within the HMG-box_ROX1-like domain, respectively (*cf*. Figures S3 and 17, Tables 2 and S3).
3. The coproduced full-length MAT1-2-1 proteins AGW27540 and AGW27544 were generated by the “homogenous” *O. sinensis* strain CS18-266 and the heterogeneous strain CS34-291. The predicted 3D structures of these proteins are not available in the AlphaFold database (*cf*. Tables 1−2, Panel (A) of Figure 17). These proteins contain Y-to-H substitutions within the HMG-box_ROX1-like domains and show 99.6% identity to the authentic MAT1-2-1 protein AEH27625 (*cf*. Figure S3).

Panel (B) of Figure 17 includes 3 *O. sinensis* strains: CS68-5-1216 and CS71-1220 from **Group I** and CS71-1218 from **Group II** (Table 2). Strain CS71-1218 is “homogenous”, strain CS71-1220 is heterogeneous, and strain CS68-5-1216 has unknown homo/heterogeneous characteristics [14,36] (*cf*. Table 1). All 3 strains produced the MAT1-1-1 proteins (AGW27532, AGW27535, and AGW27533; *cf*. Figures 9 and 17) with 97- and 8-residue truncations at the N- and C-termini, respectively, which conform to the same AlphaFold 3D structural model (U3N6U8 marked with the suffix symbol “†” in Table 2). Simultaneously, strains CS68-5-1216 and CS71-1220 produced the MAT1-2-1 proteins AGW27552 and AGW27555 with 75- and 74-residue truncations, respectively, at the C-termini, whereas strain CS71-1218 produced the MAT1-2-1 protein AGW27553 with a single residue deletion at the C-terminus (*cf*. Figures 3,15−17, and S3). These 3 MAT1-2-1 proteins contain Y-to-H substitutions within the HMG-box_ROX1-like domains, whereas the full-length protein AGW27553 contains an additional K-to-X (lysine-to-an unidentified residue) substitution in the HMG-box_ROX1-like domain. They show 99.6%, 99.6% and 99.2% similarity to the authentic MAT1-2-1 protein AEH27625 (*cf*. Figure S3) and exhibit distinct 3D structures under the AlphaFold codes U3N6W6, U3N9W5, and U3N9X0, respectively.

Panel (C) of Figure 17 lists the **Group-I** “homogenous” *O. sinensis* strain CS71-1219 and the **Group-II** strain CS70-1212, which have unknown homo/heterogeneous characteristics (Tables 1−2). These 2 strains produced the MAT1-1-1 proteins AGW27534 and AGW27531 with 110- and 8-residue truncations at the N- and C-termini, respectively (*cf*. Figures 12 and 17), which conform to the same AlphaFold 3D structural code U3NE87 (labeled with a suffix symbol “*” in Tables 1−2). Simultaneously, strain CS71-1219 produced the MAT1-2-1 protein AGW27554, which is 65-residue truncated at the C-terminus and contains Y-to-H and Q-to-T (glutamine-to-threonine) substitutions within the HMG-box_ROX1-like domain. These variations resulted in an altered tertiary structure under the AlphaFold 3D structural code U3NEA9 (*cf*. Figures 14 and 17). In contrast, strain CS70-1212 simultaneously produced the full-length MAT1-2-1 protein AGW27551, which contains only a Y-to-H substitution within the HMG-box_ROX1-like domain (*cf*. Figure S3, Table 2). No AlphaFold-predicted 3D structural model has been assigned to the MAT1-2-1 protein AGW27551.

Panel (D) of Figure 17 lists 2 **Group-II** strains, CS70-1211 and CS70-1208; the former has unknown homo/heterogeneous characteristics, and the latter is heterogeneous (Tables 1−2). These strains produced the MAT1-1-1 proteins AGW27530 and AGW27529, whose N- and C-termini were truncated by 97 and 8 residues, respectively. Both proteins contain F-to-V (phenylalanine-to-valine) substitutions within the MATα_HMGbox domains, which conform to the same AlphaFold 3D model U3NE79 labeled with a suffix symbol “‡” in Tables 1−2 (*cf*. Figures 10 and 17). These strains simultaneously produced the full-length MAT1-2-1 proteins AGW27550 and AGW27549, which contain Y-to-H (tyrosine-to-histidine) substitutions within the HMG-box_ROX1-like domains (*cf*. Tables 1−2; Figures 17 and S3) and show 99.6% sequence identity to the authentic MAT1-2-1 protein AEH27625. No AlphaFold-predicted 3D structural models have been assigned to the 2 MAT1-2-1 proteins.

#### 3.5d *O. sinensis* strains simultaneously produced MAT1-2-1 proteins with the same tertiary structures and MAT1-1-1 proteins with various tertiary structures

As shown in Table 2, the **Group-I** “homogenous” strain CS68-2-1229 and the heterogeneous strain CS71-1220 produced the 74-residue truncated MAT1-2-1 proteins AGW27548 and AGW27555, respectively, with Y-to-H substitutions within the HMG-box_ROX1-like domains (*cf*. Figures 15 and 17). These 2 proteins share the same tertiary structures under the AlphaFold code U3N9W5 labeled with the suffix symbol “⁑” in Tables 1−2. These strains simultaneously produced the MAT1-1-1 proteins AGW27528 and AGW27535, which have 63- and 8-residue truncations and 97- and 8-residue truncations at the N- and C-termini of the proteins, respectively, and form distinct 3D structures under AlphaFold codes U3N9T9 and U3N6U8 labeled with the suffix symbols “⁋” and “†” in Tables 1−2, which were fall differently under Panels (A) and (B) of Figure 17, respectively (*cf*. Figures 5 and 9).

Panel (E) in Figure 17 (highlighted by the open rectangle in blue) includes 5 **Group-II** *O. sinensis* strains (CS560-961, CS76-1284, CS561-964, CS25-273, and CS68-2-1228), which produced the full-length MAT1-2-1 proteins (AGW27542, AGW27545, AGW27546, AGW27547, and AGW27556; *cf*. Figure S3), respectively. These MAT1-2-1 proteins contain the same Y-to-H substitutions within the HMG-box_ROX1-like domains. Unfortunately, only the protein AGW27542 was given an AlphaFold 3D structural code D7F2E3; the remaining 4 proteins exhibited the same structures as the protein AGW27542 but were not assigned an AlphaFold 3D structural code. These strains simultaneously generated truncated MAT1-1-1 protein variants (AGW27522, AGW27525, AGW27526, AGW27527, and AGW27536; *cf*. Figures 9 and 17; Table 2) with different truncations (63-/8-residue, 66-/10-residue, 63-/13-residue, 77-/18-residue, and 95-/8-residue truncations) at the N-/C-termini, respectively, and contained various amino acid substitutions, as shown in Table 2 and Figures 5−8, 11, and 17. The diverse variations result in altered tertiary structures of the MAT1-1-1 proteins under distinct AlphaFold 3D structural codes (U3N6U0, U3N919, U3N7G5, U3N6U4, and U3N7H7) (*cf*. Table 2, Figures 5−8 and 11).

#### 3.5e Distinct patterns of variation between the mating-type genes in the genome of pure *H. sinensis* and the mating proteins derived from the *O. sinensis* strains

Table S7 summarizes the co-occurrence and differential occurrence of the MAT1-1-1 and MAT1- 2-1 proteins in *C. sinensis* insect‒fungal complexes, wild-type *C. sinensis* isolates, and *O. sinensis* strains, including various *O. sinensis* genotypes. Among the 183 total samples, the MAT1-1-1 and MAT1-2-1 proteins co-occur in 25.1% of the samples but differentially occur in 52.5% and 22.4% of the samples, respectively.

Genomically, both the *MAT1-1-1* and *MAT1-2-1* genes are present in the genome assemblies ANOV00000000 and LKHE00000000 of *H. sinensis* strains Co18 and 1229 (GC-biased Genotype #1 of *O. sinensis*), respectively [64,93]. However, the *MAT1-1-1* gene is absent in the genome assemblies LWBQ00000000 and NGJJ00000000 of the *H. sinensis* strains ZJB12195 and CC1406-20395, respectively, and the *MAT1-2-1* gene is absent in the genome assembly JAAVMX000000000 of the *H. sinensis* strain IOZ07 [94−96]. However, this pattern of the differential absence of the *MAT1-1-1* and *MAT1-2-1* genes in the genome of *H. sinensis* was not observed at the protein level, *i.e.*, all the 20 *O. sinensis* strains simultaneously produced the MAT1-1-1 and MAT1-2-1 proteins in pairs without omissions. The inconsistency of the absence patterns at the genetic and protein levels indicates divergent fungal origins of the MAT1-1-1 and MAT1-2-1 proteins coproduced by the 20 *O. sinensis* strains.

Figure S1 of [65] revealed Y-to-M (tyrosine-to-methionine) substitutions in the MATα_HMGbox domains of MAT1-1-1 proteins encoded by the genome assemblies LKHE00000000 and JAAVMX000000000 of the *H. sinensis* strains 1229 and IOZ07, respectively, but not by the genome assembly ANOV00000000 of the *H. sinensis* strain Co18 [64,65,93,96]. However, Y-to-M substitution was not detected in the MATα_HMGbox domains of variant MAT1-1-1 proteins derived from the 20 *O. sinensis* strains.

Figure S2 of [65] revealed S-to-A (serine-to-alanine) substitutions in the HMG-box_ROX1-like domains of the MAT1-2-1 proteins encoded by the genome assemblies of ANOV00000000, LKHE00000000, LWBQ00000000, and NGJJ00000000 of *H. sinensis* strains Co18, 1229, ZJB12195, and CC1406-20395, respectively. However, S-to-A substitution does not occur in the HMG-box_ROX1-like domains of MAT1-2-1 proteins, regardless of whether they are truncated or full-length proteins derived from the 20 *O. sinensis* strains.

Aggregating the observations presented in Section 3.5, the 20 *O. sinensis* strains simultaneously produced the MAT1-1-1 and MAT1-2-1 proteins in pairs, with differential truncations and diverse amino acid substitutions. These findings highlight the genetic variations within the *MAT* loci in the fungal genomes of impure *O. sinensis* strains, regardless of whether the strains were previously reported as “homogenous” or heterogeneous by Li *et al.* [36]. The apparent differences in the presence/absence of the *MAT1-1-1* and *MAT1-2-1* genes in the genome assemblies of the 5 *H. sinensis* strains were not replicated at the protein level in the coproduction of MAT1-1-1 and MAT1-2-1 proteins in pairs from the 20 *O. sinensis* strains. Because repetitive copies of the *MAT1-1-1* or *MAT1-2-1* genes are not present in the *H. sinensis* genome [69], these observations suggest that the paired MAT1-1-1 and MAT1-2-1 proteins are not necessarily the products of GC-biased Genotype #1 *H. sinensis* but are instead most likely the products of different co-occurring heterospecific or genotypic fungi within the impure *O. sinensis* strains.

## 4. Discussion

### 4.1 Altered tertiary structures of the DNA-binding domains of mating proteins produced by various *O. sinensis* strains

Li *et al*. [67,68] previously reported the differential occurrence, differential transcription, and alternative splicing of *MAT1-1-1* and *MAT1-2-1* and pheromone receptor genes in *H. sinensis*, wild-type *C. sinensis* isolates, and the *C. sinensis* insect−fungal complex. The full-length MAT1-1-1 and MAT1-2-1 proteins contribute to 15 and 17 heteromorphic stereostructures, belonging to 5 and 5 Bayesian clusters, respectively [69], whereas the DNA-binding domains of the full-length mating proteins derived from wild-type *C. sinensis* isolates clustered into 5 and 2 branched Bayesian clusters, respectively, as summarized in Tables S6−S9 of [65]. Thus, the evidence at the genetic, transcriptional, and protein structural levels suggests that *O. sinensis* is self-sterile and employs a heterothallic, hybrid, or even parasexual strategy to accomplish sexual reproduction during the lifecycle of the *C. sinensis* insect–fungal complex [65,67–69]. The data analyzed in this paper extend the previous observations by focusing on differential truncations and diverse amino acid substitutions within the MATα_HMGbox and HMG-box_ROX1-like domains of the MAT1-1-1 and MAT1-2-1 proteins, respectively, which are coproduced simultaneously in pairs by the 20 *O. sinensis* strains (Figure 17; Tables 1−2). N- and/or C-terminal truncations and 1−4 amino acid substitutions within the DNA-binding domains significantly alter the hydrophobic properties and modify the structures of the functional domains of mating proteins, particularly the tertiary structures of the hydrophobic cores formed by 3 α-helices within the DNA-binding domains. These structural variations ultimately modulate the synergistic interactions between the key functional domains of mating proteins during the sexual reproduction of *O. sinensis*.

The HMGbox domains in mating proteins are multifunctional motifs that are central to the transcriptional regulation of mating-related genes, *O. sinensis* sexual reproduction, and host adaptation within the *C. sinensis* insect‒fungal complex [59,65,67–69]. It has been inferred that the MATα-HMGBox in the MAT1-1-1 protein has a conserved ancestral mating-type HMG lineage *via* conservative continuation model, whereas the HMG-box_ROX1-like domain within the MAT1-2-1 protein represents evolutionary cooperation of a different regulatory lineage (*e.g*., ROX1-like) through functional convergence, where it was repurposed to control mating-type specification [62,63,97−100].

The MATα_HMGbox and HMG-box_ROX1-like domains of the MAT1-1-1 and MAT1-2-1 proteins, respectively, are structurally defined DNA-binding domains characterized by a hydrophobic core containing 3 asymmetrically arranged α-helices. The hydrophobic core, which is composed of tightly packed amino acid residues, is the foundation of the structures and provides a precise conformational stability and constraints that enable the 3 α-helices to form the characteristic L-shaped fold [73,90,91]. These α-helices converge and create stable resistance to environmental perturbations that might unfold or distort the domains. The unique tertiary structures of these sequence-specific DNA binders preferentially ensure high-affinity, sequence-dependent binding to short AT-rich DNAsequences within the promoters of their target mating-type-specific genes [61,101−106]. The sequence-specific binding of the HMG-box domains to AT-rich promoter motifs serves as the initial trigger, thereby playing pivotal roles in controlling fungal mating identity and gene expression [62,97,107,108].

Disruption of this hydrophobic core, whether by 1−4 amino acid substitutions at distinct sites that introduce incompatible residues or by differential peptide chain truncations that remove parts of it, as identified in this study, directly impairs structural integrity and abolishes the crucial synergistic functions of the domains in DNA binding specificity in recognizing the target DNA segments and the transcriptional regulation of mating-type-specific genes. These variations have significant consequences for the regulation of *O. sinensis* sexual reproduction during the sexual life of the *C. sinensis* insect−fungal complex. Compared with the authentic MAT1-1-1 and MAT1-2-1 proteins of *O. sinensis* analyzed previously by Li *et al*. [65,69], the differentially altered DNA-binding domains clustered into diverse clades/branches in the Bayesian clustering trees [65,69].

Although the MATα_HMGbox domain within the MAT1-1-1 protein and the HMG-box_ROX1-like domain within the MAT1-2-1 protein both belong structurally to the HMG-box superfamily and share a conserved L-shaped fold that enables DNA binding and bending, they are functionally distinct in the context of mating-type regulation in *O. sinensis* because they regulate different gene sets, resulting in opposite regulatory outcomes. The MATα_HMGbox domain of the MAT1-1-1 protein activates the transcription of genes encoding MAT1-1-specific receptors or other regulators. Although the HMG-box_ROX1-like domain of the MAT1-2-1 protein binds to specific DNA sequences in the *MAT1-2*-specific gene promoter encoding the *MAT1-2*-specific pheromone receptor, it may overlap differently with the MATα_HMGbox targets and act through an opposite pathway to counterbalance the activity of the MAT1-1-1 protein [109–111, the MATα_HMGbox domain primarily functions as an activator of its cognate mating-type genes (*i.e., MAT1-1* genes). In contrast, the HMG-box_ROX1-like domain acts as a repressor of genes specific to the opposite mating type (*MAT1-1* or “a-specific” genes) and may also serve as an activator for *MAT1-*2-specific genes. This repressive capacity represents a key functional hallmark derived from its homology to ROX1 [112]. Thus, despite their structural similarity as DNA-binding and DNA-bending HMG-box domains, the MATα_HMGbox and HMG-box_ROX1-like domains are functionally specialized to control distinct, complementary sets of target genes, thereby defining the two different mating types. Their divergent regulatory outputs (activation *vs*. repression of key target gene sets) constitute the molecular and protein structural basis of mating-type specification during the sexual reproduction of *O. sinensis* during the lifecycle of the *C. sinensis* insect–fugal complex.

The putative heterodimerization between the MATα_HMGbox domain of the MAT1-1-1 protein and the HMG-box_ROX1-like domain of the MAT1-2-1 protein has been proposed as a potential regulatory mechanism to enforce outbreeding and initiate the sexual cycle in heterothallic fungi [113–115]. However, because of the lack of direct evidence in *O. sinensis*, it remains a compelling hypothesis and future research must confirm the occurrence of this heterodimerization, characterize the interaction interface, and define its precise regulatory targets and functional effects on mating and fruiting body development in this economically and ecologically important insect–fungal complex. Nevertheless, the MATα_HMGbox and HMG-box_ROX1-like domains may interact electrostatically and complementarily, providing a plausible physicochemical basis for cooperative synergy in DNA binding. This interaction could coordinate the transition from asexual growth to sexual development, thereby distorting target DNA to repress haploid programming and activating the mating/meiosis genes of *O. sinensis* during the sexual life of the *C. sinensis* insect–fungal complex [116]. Nevertheless, direct experimental validation using protein biochemistry and reproductive physiological approaches is warranted to confirm this hypothetical model, clarify its species-specific characteristics, and determine the effects of protein-level variations on the complementary interactions and alterations in the mating function of *O. sinensis* [65,67−69], even as the genome-independent, multifungal, and multigenotypic nature of *O. sinensis* remains to be fully elucidated [9,10,12,13,19,24,27,30,35,37,38,54,117]. Such protein-level variations may provide critical evidence to support or refute the hypotheses of the anamorph–teleomorph linkage and sexual reproduction strategy of *O. sinensis* [14,20,36,45,64,65, 67–69,72].

### 4.2 Heterogenous fungal sources of the MAT1-1-1 and MAT1-2-1 proteins simultaneously produced in pairs by the impure *O. sinensis* strains

#### 4.2a Phylogenetic heterogeneity of *O. sinensis* strains

By employing a dual-fluorescence microscopy technique, Bushley *et al*. [14] observed the multicellular heterokaryotic microstructures of *C. sinensis* ascospores and hyphae, which contain mono-, bi-, tri-, and tetra-nucleate cellular structures. Zhang & Zhang [72] subsequently commented on these heterokaryotic microstructures and expressed doubts regarding whether binucleate (as well as tri- and tetra-nucleate) cells might inconsistently harbor homogenous or heterogeneous hereditary substances. Molecular biological observations reported by Li *et al*. [12,13,36–38] revealed multiple fungal components in the ascospores and ascocarps of *C. sinensis* insect‒fungal complex and in purportedly “pure” *H. sinensis* strains. These findings provide indisputable evidence supporting the hypothesis proposed by Zhang & Zhang [72] that multicellular heterokaryotic cells indeed contain heterogeneous hereditary substances.

Evidence of culture dependence was reported by Li *et al*. [36], who deposited the ITS sequences of 15 of the 20 *O. sinensis* strains in GenBank. Among the 15 *O. sinensis* strains, 10 were reported as “homogenous” strains, purportedly harboring only the GC-biased Genotype #1. The remaining 5 strains were characterized as heterogeneous, containing the ITS sequences of both GC-biased Genotype #1 (Group A) and AT-biased Genotype #17 (Group C). Li *et al*. [36,118,119] hypothesized that AT-biased genotypic sequences are nonfunctional repetitive copies within the single genome of GC-biased *H. sinensis* and are referred to as “ITS pseudogenes” and “rRNA pseudogenes”, which are generated through repeat-induced point (RIP) mutation. However, subsequent studies have demonstrated that the sequences of Genotypes #2−17 of *O. sinensis*, including both GC- and AT-biased genotypes, do not reside in the genomic assemblies of 5 pure *H. sinensis* strains [9,10,12,13,38,120]. Thus, the 17 *O. sinensis* genotypes are actually genomically independent and belong to distinct fungal lineages, although they may share a common ancestral heredity [24]. These findings directly invalidate the hypotheses that “RIP” mutation induces AT-biased “ITS pseudogenes” and “rRNA pseudogenes” as nonfunctional repetitive copies in the genome of GC-biased Genotype #1 *H. sinensis* [36,118,119].

The less sophisticated research methodology employed by Li *et al*. [36] was analyzed in Section 3.5a. The technical limitation led to an imbalance in the amplification efficiency of coexisting fungal sequences. Consequently, Li *et al*. [36] detected the sequence(s) that exhibited the highest PCR amplification efficiency, likely precluding the comprehensive detection of all the coexisting fungal taxa in the samples. Notably, the different abundances (high *vs*. low) of amplicons primarily reflect the level of PCR amplification efficiency and velocity under specific research settings and, obviously, should not be equated with the natural ratio of coexisting fungi in the tested samples. The natural ratio of coexisting fungal taxa can be quantified using other specialized non‒PCR techniques, such as Southern blotting with the use of specific probes [27]. Notably, naturally occurring, less abundant fungal taxa may still play synergistic roles in the lifecycle of the *C. sinensis* insect–fungal complex.

Furthermore, Li *et al*. [12,13,37] adopted an exhaustive culture-independent approach, including the use of multiple primers, “touch-down” PCR protocols, and the selection and sequencing of multiple (>30) white colonies, as well as MALDI-TOF SNP mass spectrum genotyping approaches using multiple extension primers, to exhaustively investigate the comprehensive fungal composition within *C. sinensis* ascospores and the “pure” *H. sinensis* strains Hs-Ma-4, Hs7.7, Hs7.8, and Hs-Qing2 (gifts from the renowned mycologist Prof. Gou YL). These studies demonstrated that *C. sinensis* ascospores and “pure” *H. sinensis* strains co-occur with genomically independent fungal taxa, including the psychrophilic GC-biased Genotypes #1 and #14 and AT-biased Genotypes #5−6 and #16 of *O. sinensis*, as well as the mesophilic *Paecilomyces hepiali* (recently reclassified as *Samsoniella hepiali* [121]). These findings clearly indicate that the study by Li *et al*. [36] overlooked some coexisting fungal sequences with low PCR amplification efficiency and nonculturable fungal components under particular experimental conditions. Thus, the origins of paired MAT1-1-1 and MAT1-2-1 proteins with differential truncations and various amino acid substitutions at distinct sites require further clarification using rigorously designed mycological and molecular protocols by the custodians of the original *O. sinensis* strains.

The paired heteromorphic MAT1-1-1 and MAT1-2-1 proteins are likely to have distinct fungal origins and were possibly derived from the co-occurring GC-biased Genotypes #1 and #3 and AT-biased Genotype #17 of *O. sinensis* in wild-type *C. sinensis* isolates [65] and the *O. sinensis* strains analyzed in this study. Given the limited available information on the diverse coexisting genotypes of *O. sinensis*, the different pairings of the MAT1-1-1 and MAT1-2-1 protein variants that exhibit differential truncations and diverse amino acid substitutions at distinct sites were coproduced simultaneously by the differentially coexisting genomically independent fungi in *O. sinensis* strains [6,9–13,15–44].

#### 4.2b Differential pairings of the MAT1-1-1 and MAT1-2-1 proteins simultaneously produced by *O. sinensis* strains

Table S7 summarizes the differential occurrence or co-occurrence of both authentic and variant MAT1-1-1 and MAT1-2-1 proteins in *C. sinensis* insect−fungal complexes, wild-type *C. sinensis* isolates, and various *O. sinensis* strains (including GC-biased Genotype #1 *H. sinensis* strains). The differential occurrence pattern of the *MAT1-1-1* and *MAT1-2-1* genes in the genome of the Genotype #1 *H. sinensis* protein is inconsistent with the genetic expression pattern at the protein level; namely, the MAT1-1-1 and MAT1-2-1 proteins were produced simultaneously by the 20 *O. sinensis* strains without any omission. The paired MAT1-1-1 and MAT1-2-1 proteins exhibit differential truncations and diverse amino acid substitutions (*cf*. Tables 2 and S5, Figures 1−17 and S3) [14,36,69]. The distinct pairings of the highly divergent MAT1-1-1 and MAT1-2-1 protein variants that were simultaneously and differentially produced by the 20 *O. sinensis* strains present a complex, nonuniform pairing pattern (*cf*. Table 2 and Figure 17), indicating diverse fungal origins of the mating proteins from co-occurring fungal taxa in the samples. In addressing the issue of overlooked co-occurring fungal taxa due to the unsophisticated study designed by Li *et al*. [36] (*cf*. Section 3.5a), namely, the nonculturable fungal taxa and low-abundance coexisting fungal sequences that arise from less efficient PCR amplification and sequencing bias associated with the suboptimal experimental design, the 20 *O. sinensis* strains presumably exhibit a much greater diversity of detected and undetected fungal sequences than the “homogenous” and heterogeneous cultures of *C. sinensis* monoascospores and tissues of caterpillar bodies previously reported by Li *et al*. [36]. Further clarification is warranted, employing unbiased and rigorous culture-independent methods to comprehensively profile coexisting genomically independent genotypes of *O. sinensis* and heterospecific fungal taxa [12,13,27,35,37].

#### 4.2c Paradoxical dilemmas regarding the highly possible heterogeneity of purportedly “pure” *O. sinensis* strains

The first dilemma concerns the purportedly homogenous strain CS68-2-1229 of **Group I** (highlighted in dark green and underlined in Figures 2 and 4). Only the GC-biased Genotype #1 was previously detected in this “pure” culture of *C. sinensis* monoascospores under the experimental settings adopted by Li *et al*. [36], which was incapable of completely covering all co-occurring fungal taxa (*cf*. Section 3.5a). Two MAT1-1-1 protein sequences from this strain have been deposited in GenBank: a full-length MAT1-1-1 protein, AGW27560, containing 372 amino acids, and a truncated MAT1-1-1 protein, AGW27528, containing 301 amino acids (*cf*. Figure 5). These 2 proteins share 100% sequence identity, yet AGW27528 exhibits only 81% query coverage with 63- and 8-residue truncations at the N-and C-termini, respectively (*cf*. Figure 17, Table 2). The truncation pattern was likely attributable to the use of different primers for amplifying the *MAT1-1-1* gene or its transcript from the total RNA pool in the experiments. This strain simultaneously produced the truncated MAT1-2-1 protein AGW27548 (175 amino acids; *cf*. Figure 15), which shares 99.4% sequence identity with the authentic full-length MAT1-2-1 protein and has a Y-to-H substitution within the HMG-box_ROX1-like domain; notably, it shows only 70% sequence coverage because of a 74-residue truncation at the C-terminus relative to that of the authentic full-length MAT1-2-1 proteins, which contain 249 amino acids. All 3 mating protein sequences derived from strain CS68-2-1229 were released in GenBank on the same date (28-SEP-2013). The truncated MAT1-1-1 AGW27528 and MAT1-2-1 protein AGW27548 were derived from cDNAs that were amplified using the primer pair *m1F3*/*m1R3* to amplify *MAT1-1-1* cDNAand the primer pair *Mat1-2F*/*Mat1-2R* to amplify *MAT1-2-1* cDNA from the same batch of total RNA pool extracted from the strain CS68-2-1229 (*cf*. Figures S1−S2) [14,36,70]. Figure 2 shows that the MATα_HMGbox domain (162 amino acids) of the MAT1-1-1 protein AGW27528 is among the longest domains of all 9 truncated MAT1-1-1 proteins. In contrast, Figure 4 shows that the HMG-box_ROX1-like domain (49 amino acids) of the MAT1-2-1 protein AGW27548 (*cf*. Figures 4 and 15) is 10 residues shorter than the longest truncated HMG-box_ROX1-like domain (59 amino acids) of the MAT1-2-1 protein AGW27543 [65,70]. The truncated MAT1-2-1 protein AGW27548, which harbors a Y-to-H substitution, exhibits altered secondary and tertiary structures under the AlphaFold 3D structural code U3N9W5 (*cf*. Figure 15, Table 2). Li *et al*. [69] reported no repetitive copies of the *MAT1-1-1* and *MAT1-2-1* genes in the genome of the *H. sinensis* strain. These findings suggest that the MAT1-1-1 protein AGW27528 is likely encoded by the authentic *MAT1-1-1* gene in the *H. sinensis* genome. However, the MAT1-2-1 protein AGW27548 is presumably either encoded by a variant *MAT1-2-1* gene within the *H. sinensis* genome or by an independent co-occurring fungal taxon in the actual “impure” *O. sinensis* strain CS68-2-1229. Thus, the true genetic characteristics of strain CS68-2-1229 require re-evaluation: whether this strain is genuinely homogenous, containing only Genotype #1 *H. sinensis*, or is indeed heterogeneous, harboring 2 or more fungal taxa, remains to be determined.

Another paradoxical “homogenous” *O. sinensis* strain, CS561-964 (**Group II**), was also reported to harbor only GC-biased Genotype #1 *H. sinensis* [14,36]. This strain produced the MAT1-1-1 protein AGW27526, featuring 63- and 13-residue truncations at the N- and C-termini, respectively, along with 4 amino acid substitutions (CDRA-to-SSEF) within the MATα_HMGbox domains (*cf*. Figures 7 and 17, Tables 1−2). Clearly, the altered protein AGW27526 is encoded by a variant *MAT1-1-1* gene, plausibly from a genome-independent, co-occurring fungal taxon in the *O. sinensis* strain. The ITS sequence of this fungal taxon was most likely overlooked in the study by Li *et al*. [36] because of the use of suboptimal culture-dependent experimental protocols (*cf*. Section 3.5a).

Analyses of the purportedly homogenous *O. sinensis* strains raise fundamental questions regarding fungal purification, *in vitro* culture characteristics, and comprehensive profiling of all co-occurring fungal taxa in such cultures. Two predicaments remain unresolved: First, do the purportedly homogenous strains listed in Tables 1−2 truly harbor only GC-biased Genotype #1, or are they indeed heterogeneous and contain 2 or more fungal taxa? Second, do the characterized heterogeneous strains contain only the GC-biased Genotype #1 and AT-biased Genotype #17 or an even greater number of undetected coexisting fungal taxa? A reasonable hypothesis is that the differentially paired truncated and full-length MAT1-1-1 and MAT1-2-1 protein variants, which exhibit diverse amino acid substitutions at distinct sites of variation, are not necessarily produced by GC-biased Genotype #1 *H. sinensis*. Instead, these protein variants likely originate from AT-biased Genotype #17 (*cf*. Tables S1−S2) or from other uncharacterized co-occurring fungal taxa that were not detected in the culture-dependent, suboptimal study by Li *et al*. [36].

### 4.3 Self-sterility of *O. sinensis* and the requirement for sexual partners to accomplish heterothallic or hybrid reproduction

Wei *et al*. [45] proposed that *H. sinensis* (GC-biased Genotype #1 of *O. sinensis*) represents the sole anamorph of *O. sinensis* teleomorph. Zhang *et al*. [51] proposed the implementation of the “One Fungus=One Name” nomenclature principle established by the International Mycological Association [52,53] and replaced the anamorphic name *H. sinensis* with the teleomorphic name *O. sinensis* [45]. This renaming implementation, however, was not only applied to GC-biased Genotype #1 *H. sinensis*, which was used as the nomenclature reference, but was also uniformly applied to all 17 genomically independent genotypes of *O. sinensis*, although they may share a common hereditary ancestor [9,10,24,54]. Eleven of the 17 genotypes are GC biased, whereas the remaining 6 genotypes are AT biased. The genotypes in various combinations coexist in different compartments of the *C. sinensis* insect‒fungal complex, and their natural abundance changes dynamically and reciprocally during *C. sinensis* maturation, indicating a genomically independent nature [9,10,12,13,27,35,37]. In support of the sole anamorph hypothesis for *H. sinensis* [45], Bushley *et al*. [14] and Hu *et al*. [64] studied the mating-type genes in the genome of GC-biased Genotype #1 *H. sinensis* and proposed the self-fertilization hypothesis for *O. sinensis* to accomplish homothallic or pseudohomothallic reproduction. However, several research groups [64,122,123] have reported unsuccessful attempts to artificially cultivate the fruiting bodies and ascospores of *C. sinensis* insect–fungal complexes using pure fungal cultures. Zhang *et al*. [51] then summarized the 40-year history of unsuccessful attempts to cultivate insect–fungal complexes in academic research-oriented settings. Qin *et al*. [124] elaborated on the key obstacles hindering the artificial cultivation of *C. sinensis* fruiting bodies and ascospores. Li *et al*. [125] further attributed the decades-long history of failures in the cultivation of *C. sinensis* insect–fungal complexes to “the low rate of the fungal invasion to host insects and primordium induction after infection”. Compared with the nearly noninfectious characteristics of pure *H. sinensis*, Li *et al*. [37] advanced an inoculation technique using cocultures of natural fungal clusters with a natural abundance ratio of multiple genotypic and heterospecific fungal taxa isolated from the *C. sinensis* insect–fungal complex. This advanced technique resulted in up to 39-fold greater infectivity in the larvae of *Hepialus armoricanus* and significantly shortened the latency (from 35−50 days to 5−8 days) from inoculation to mummification/death. Evidently, the hypotheses of contentious sole anamorph and concomitant self-fertilization have not yet been scientifically validated in academia to meet all 4 criteria of Koch’s postulations.

In contrast to the hypothesis that *H. sinensis* represents the sole anamorph of *O. sinensis* [45], molecular studies have revealed complex genetic diversity of *O. sinensis,* and this Latin name encompasses at least 17 distinct genotypes [9,10,12,13,27,30,35,54]. The sequences of Genotypes #2−17 of *O. sinensis* are not present in the genome assemblies of 5 Genotype #1 *H. sinensis* strains, indicating that they represent diverse genome-independent fungal taxa [9,10,24,30]. A medicinal mycology group led by Prof. S.P. Wasser reported that *H. sinensis* and *Tolypocladium sinensis* serve as conjoint anamorphs of *O. sinensis* [34], supporting the prior detection of *T. sinensis* and *T. sinense* from natural *C. sinensis* insect–fungal complexes [15,20,21,25]. Quandt *et al*. [126] subsequently proposed taxonomic revisions within the genus *Tolypocladium*, reassigning several species to the family Ophiocordycipitaceae. To date, the taxonomic positions of 17 evolutionarily related genotypes as independent fungi, as well as the precise anamorph–teleomorph relationship of *O. sinensis*, remain unresolved. With respect to reproductive modes, although early hypotheses propose homothallic or pseudohomothallic reproduction for *O. sinensis* [10,58], Zhang and Zhang [72] reported the differential presence of *MAT1-1-1* and *MAT1-2-1* genes across numerous wild-type *C. sinensis* isolates and proposed a facultative hybridization hypothesis for *O. sinensis*. Li *et al*. [67,68] further documented the differential occurrence, alternative splicing, and differential translation of the *MAT1-1-1* and *MAT1-2-1* genes and the pheromone receptor genes. More recently, the studies by Li *et al*. [65,69] and the present study revealed heteromorphic tertiary structures of the MAT1-1-1 and MAT1-2-1 protein variants, particularly variations within the DNA-binding domains of the mating proteins. These structural observations provide protein-level evidence for heterogeneous fungal origins of mating-type loci in numerous wild-type *C. sinensis* isolates and *O. sinensis* strains. Notably, Li *et al*. [69] reported that both the *MAT1-1-1* and *MAT1-2-1* genes exist as single copies in the *H. sinensis* genome with no repetitive genomic copies. Collectively, the evidence from genetics, transcription, and protein structure is inconsistent with the self-fertilization hypothesis for *O. sinensis* and instead supports self-sterility, implying that *O. sinensis* relies on heterothallic mating, hybridization, or even parasexual outcrossing to accomplish sexual reproduction [13,65,67−69,127–136]. Similar self-sterile reproductive strategies have been documented in many other *Ophiocordyceps* species, which require compatible mating partners to undergo sexual reproduction [104,137–141].

Wei *et al*. [46] reported the successful industrial cultivation of *C. sinensis* insect–fungal complexes. However, this study revealed a puzzling species paradox between the input and output of the artificial cultivation project: 3 GC-biased *H. sinensis* strains were used as inoculants, yet only the teleomorph of AT-biased Genotype #4 was detected in the fruiting bodies of the cultivated *C. sinensis* insect–fungal complexes. Hence, the sole teleomorphic AT-biased Genotype #4 identified in the cultivated fruiting bodies replaced the inoculant GC-biased Genotype #1 strains, resulting in a species contradiction. The sequences of Genotypes #2−17 of *O. sinensis* are not present in the genome assemblies ANOV00000000, JAAVMX000000000, LKHE00000000, LWBQ00000000, or NGJJ00000000 of the pure *H. sinensis* strains Co18, IOZ07, 1229, ZJB12195, and CC1406-20395, respectively [9,10,12,13,24,30,35,36,38,64,93−96,117,120]. Instead, these genotypic sequences are associated with distinct genome-independent fungal taxa. These results invalidate the unfounded assumptions proposed by Li *et al*. [36,118,119] that AT-biased genotypes are nonfunctional “ITS pseudogenes” or “rRNA pseudogenes” that were generated through “RIP” mutation in the *H. sinensis genome*. Supporting evidence at the genomic and transcriptomic levels is not available to validate these unfounded speculations [38,120]. In contrast to the significantly variable maturation-related dynamics of AT-biased genotypes of *O. sinensis*, GC-biased *O. sinensis* genotypes maintain a relatively constant low abundance in the stromata throughout the entire maturation course of the *C. sinensis* insect–fungal complex [27,35]. These observations suggest that the 17 genome-independent genotypes of *O. sinensis*, regardless of whether they are GC or AT biased, represent distinct fungal species [10,12,13,30], further reinforcing the invalidation of the RIP mutation/pseudogene hypotheses proposed by Li *et al*. [36,38,118−120]. The successful industrial cultivation of *C. sinensis* insect‒fungal complexes, coupled with the apparent species paradox, indicates that the sole anamorph hypothesis for *H. sinensis* proposed by Wei *et al*. [46] failed to satisfy all 4 criteria of Koch’s postulates.

Li *et al*. [125] subsequently clarified that the inoculant used in the industrial cultivation project was an *O. sinensis* conidial suspension obtained after 3 months of culture of *C. sinensis* ascospores, rather than the 3 pure *H. sinensis* strains initially described by Wei *et al*. [46]. However, Li *et al*. [125] did not disclose the molecular components of the inoculant to the public or redress the previously reported teleomorphic AT-biased Genotype #4 of *O. sinensis*in the fruiting bodies of cultivated *C. sinensis*insect–fungal complexes. On the other hand, several other studies reported that AT-biased Genotype #4 was detected in the stromata throughout all maturation stages of the *C. sinensis* insect–fungal complexes [10,27,35,36,38]. The abundance of Genotype #4 markedly altered ontogenetic dynamics; *i.e.*, it was highly abundant in immature stromata during the asexual growth phase and was substantially reduced in the mature stromata and the stromal fertile portion of the *C. sinensis* insect‒fungal complex during the sexual reproduction stage or the transition from asexual growth to sexual reproduction [9,10,12,13,35,38]. Regrettably, several studies [12,13,36,37] failed to detect AT-biased Genotype #4 from *C. sinensis* ascospores using both culture-dependent and culture-independent approaches. Thus, the dynamic maturation pattern for AT-biased Genotype #4 of *O. sinensis* and its absence in *C. sinensis* ascospores suggest that Genotype #4 may not participate in the sexual reproduction of *O. sinensis*. Consequently, the puzzling species paradox between the inoculant and the sole teleomorphic AT-biased Genotype #4 of *O. sinensis* detected in the fruiting body of the cultivated *C. sinensis* insect–fungal complex [46] remains unresolved. Unfortunately, Li *et al*. [125] did not address the genetic complexity of the ascospores of cultivated insect–fungal complexes.

Li *et al*. [65] discussed the evidence related to the industrial success of cultivating the insect–fungal complex and postulated that the success of the industrial cultivation project may be attributed to the adaptation of a “mycological impurity” strategy, facilitating the synergistic interplay of coexisting fungi. The supporting evidence includes the following: (1) Li *et al*. [125] reported that the inoculant was a culture of *C. sinensis* ascospores to address the weak infectivity of GC-biased Genotype #1 *H. sinensis*. Li *et al*. [12,13,36] reported that *C. sinensis* ascospores harbor multiple genotypic and heterospecific fungi. (2) *C. sinensis* ascospores do not contain AT-biased Genotype #4 of *O. sinensis* [10,12,13,36,38]. Therefore, the origin of teleomorphic Genotype #4 detected in the fruiting body of the cultivated *C. sinensis* insect–fungal complex by Wei *et al*. [46] might be derived from cocultures of *C. sinensis* ascospores with the stroma and/or stromal fertile portion of the *C. sinensis* insect–fungal complex that indeed contain AT-biased Genotype #4 of *O. sinensis*, along with other genotypic and heterospecific fungi. Li *et al*. [37] demonstrated that cocultures of multiple genotypic and heterospecific fungal taxa in wild-type *C. sinensis* isolates exhibit up to 39-fold greater infectivity in the larvae of *Hepialus armoricanus* than those of pure *H. sinensis.* (3) The industrial cultivation system was supplemented with soil collected from natural *C. sinensis* production areas on the Qinghai‒Tibet Plateau [46].

To date, the purification and genomic analysis of GC- and AT-biased Genotypes #2−17 of *O. sinensis* have not been documented. Thus, genomic and transcriptomic information for the divergent *MAT1-1-1* and *MAT1-2-1* genes in Genotypes #2−17 of *O. sinensis* is lacking, especially concerning the critical DNA-binding domains of the MAT1-1-1 and MAT1-2-1 proteins. Regardless of the precise biophysical details of their interaction, whether the paired MAT1-1-1 and MAT1-2-1 proteins form heterodimers or interact *via* complementary electrostatic surfaces, the synergy of paired mating proteins serves as the central and indispensable molecular trigger for mating compatibility and the initiation of sexual reproduction in *O. sinensis* [56−60]. Molecular evidence regarding the differential occurrence, alternative splicing, and differential transcription of mating-type and pheromone receptor genes in *H. sinensis* unequivocally indicates that *O. sinensis* is self-sterile and relies on heterothallism or hybrid outcrossing to complete sexual reproduction, thereby ruling out homothallism (self-fertilization) as a viable mechanism in this species [14,45,63,66–69,142]. Furthermore, the discovery of heteromorphic tertiary structures of variant MATα_HMGbox domains (in MAT1-1-1 proteins) and HMG-box_ROX1-like domains (in MAT1-2-1 proteins) provides protein-level validation that (1) self-fertilization is biochemically unfeasible and (2) coordinated heterothallic or hybrid reproduction (even parasexuality) is enforced through structural complementarity between the paired mating proteins, which are likely generated by divergent heterogeneous *O. sinensis* strains. This self-sterility hypothesis resolves the longstanding uncertainty in the cultivation of the *C. sinensis* insect–fungal complex, namely, the failure of single-species cultures to produce fruit, and underscores the role of the synergistic effect of multiple fungal symbionts on its ecological success throughout the lifecycle of the naturally occurring or artificially cultivated *C. sinensis* insect–fungal complex.

## Conclusions

This study presents evidence at the protein structure level for self-sterile *O. sinensis* and further verifies the heterothallic or hybrid reproduction of *O. sinensis*. The diverse pairings of MAT1-1-1 and MAT1-2-1 protein variants coproduced simultaneously by the 20 *O. sinensis* strains examined in this study exhibit differential truncations and 1−4 amino acid substitutions in their DNA-binding domains. The primary structural variations caused dramatic changes in the hydrophobic properties and secondary and tertiary structures of the MATα_HMGbox and HMG-box_ROX1-like domains of the differentially paired MAT1-1-1 and MAT1-2-1 proteins, respectively. The structural heterogeneity of differentially paired mating proteins not only indicates the co-occurrence of heterogeneous fungal origins in impure *O. sinensis* strains but also invalidates the hypothesis of homothallic or pseudohomothallic self-fertilization in *O. sinensis*. Complementing the genetic and transcriptional evidence of differential occurrence, differential transcription, and alternative splicing of the *MAT1-1-1* and *MAT1-2-1* genes and pheromone receptor genes, the variations in the protein structures of these DNA-binding domains of the mating proteins impair the homologous interaction required for homothallic or pseudohomothallic self-fertilization. Collectively, our findings of differential pairings of the mating protein variants in *O. sinensis* strains clarify the sexual reproduction mechanism of *O. sinensis* and confirm that the selection of suitable heterologous mating partners is an essential natural strategy for self-sterile *O. sinensis* to complete sexual reproduction during the lifecycle of *Cordyceps sinensis* insect–fungal complexes on the Qinghai–Tibet Plateau.

## Supporting information

C:\JSZ documents\Products\Cordyceps sinensis\Manuscripts\2026\2026 Os strains truncated mating proteins

## Acknowledgments

The authors are grateful to Prof. Mu Zang, Prof. Ru-Qin Dai, and Prof. Zong-Qi Liang for their consultation.

## Supplementary Materials

The following supporting information can be downloaded at www.xxxxxx.

**<u>Figure S1</u>**. Alignment of the DNA sequence of the *MAT1-1-1* gene KC437356 (6530→7748) and the encoded sequence of the MAT1-1-1 protein AGW27560 (1→ 372) derived from *Ophiocordyceps sinensis* strain CS68-2-1229 [14]. Residues in pink indicate the MATα_HMGbox domain of the *MAT1-1-1* gene and MAT1-1-1 protein; those in green indicate introns I and II. The underlined residues in red indicate the forward primer m1F3/ *MAT1-1-1* (CCACTAGGCAGACCAAAGAAG) and the reverse complementary primer m1R3/*MAT1-1-1* (CGCAAAGTAAAAGTCGTCCAGA) reported by [14]. The triplet residues (TAG) in red represent the stop codon.

**<u>Figure S2</u>**. Alignment of the *MAT1-2-1* gene HM212637 (3241→4097) and the encoded sequence of the MAT1-2-1 protein AEH27625 (1→249) derived from *Ophiocordyceps sinensis* strain CS2 [70]. Residues in pink indicate the HMG-box_ROX1-like domain of the *MAT1-2-1* gene and MAT1-2-1 proteins; those in green indicate introns I and II. The underlined residues in red indicate the forward primer *Mat1-2F*/*MAT1-2-1* (TGGAATGCGACTGACTAC GA) and the reverse complementary primer *Mat1-2R*/*MAT1-2-1* (CCAGGAGAGCTTGCTTGACT) reported previously [14,70]. The triplet residues (TAA) in red represent the stop codon.

**<u>Figure S3</u>**. Alignment of the sequences of full-length MAT1-2-1 proteins derived from **Group II** *O. sinensis* strains listed in Table 2 with that of the authentic MAT1-2-1 protein AEH27625 (under the AlphaFold code D7F2E9) derived from the *H. sinensis* strain CS2. The residues in pink indicate the HMG-box_ROX1-like domains of the proteins, those in green indicate the variant amino acids, the hyphens indicate identical amino acid residues, and the space denotes the unmatched sequence gap.

**<u>Table S1</u>**. The *O. sinensis* strains, GenBank accession numbers for the ITS nucleic acid sequences corresponding to the GenBank accession numbers for the MAT1-1-1 and MAT1-2-1 proteins, and percent similarities *vs*. GC-biased Genotypes #1−3, #7−10, and #12 of *O. sinensis*.

**<u>Table S2</u>**. The *O. sinensis* strains, GenBank accession numbers for the ITS nucleic acid sequences corresponding to the GenBank accession numbers for the MAT1-1-1 and MAT1-2-1 proteins, and percent similarities *vs*. AT-biased Genotypes #4−6 and #15−17 of *O. sinensis*.

**<u>Table S3</u>**. Amino acid scales based on the general chemical characteristics of their side chains for ProtScale analysis (https://web.expasy.org/protscale/) to predict the hydrophobicity and secondary structures (α-helices, β-sheets, β-turns, and coils) of proteins.

**<u>Table S4</u>**. GenBank accession numbers (in red in parentheses) for the full-length MAT1-1-1 proteins in the AlphaFold database under the corresponding AlphaFold UniProt codes [87].

**<u>Table S5</u>**. GenBank accession numbers (in red) for the full-length MAT1-2-1 proteins of 69 *H. sinensis* strains or wild-type *C. sinensis* isolates under the corresponding AlphaFold UniProt codes [87].

**<u>Table S6</u>**. Similarity of the 3 pairs of primers that were used by Li *et al*. [36] to amplify the ITS sequences of the *O. sinensis* strains listed in Table 1 that were isolated either from the caterpillar body specimens that were collected from various production areas on the Qinghai‒Tibet Plateau or from the cultures of mono-ascospores that were collected from mature *C. sinensis* insect‒fungal complexes obtained from a single production region (Maqên, Guoluo, Qinghai Provence of China). **<u>Table S7</u>**. Co-occurrence or differential occurrence of the MAT1-1-1 and MAT1-2-1 proteins in the *O. sinensis* strains, wild-type *C. sinensis* isolates, and *C. sinensis* insect‒fungal complex listed in the GenBank database.

## Author Contributions

Conceptualization, XZL, YLL, WL, JZQ, and JSZ; methodology, WL and JSZ; formal analysis, JSZ; investigation, XZLand JSZ; data curation, XZL and JSZ; writing–original draft preparation, JSZ; writing—review and editing, XZL, YLL, WL, JZQ, and JSZ; project administration, YLL; funding acquisition, YLL and JZQ. All authors have read and agreed to the published version of the manuscript.

## Institutional Review Board Statement

Not applicable because this paper is an *in silico* reanalysis of public data.

## Informed Consent Statement

Not applicable because this paper is a public bioinformatic data reanalysis.

## Data Availability Statement

All sequence and 3D structure data are available in the GenBank and AlphaFold databases.

