## Supplementary material for "Heteromorphic tertiary structures of MAT1-1-1 and MAT1-2-1 protein variants simultaneously produced in pairs by *Ophiocordyceps sinensis* strains": C:\JSZ documents\Products\Cordyceps sinensis\Manuscripts\2026\2026 Os strains truncated mating proteins

```
KC437356 6530  ATGACGACAAGAAATGAGGTTATGCAGCGCTTGTCTTCTGTCCGAGCTGACGTTCTTCTG  6589
AGW27560 1      M  T  T  R  N  E  V  M  Q  R  L  S  S  V  R  A  D  V  L  L   20

KC437356 6590  AACTTCCTCACGGACGATGCAATTTTCCAGCTTGCGTCCCGATATCACGAATCGACGACA  6649
AGW27560 21     N  F  L  T  D  D  A  I  F  Q  L  A  S  R  Y  H  E  S  T  T   40

                                                          m1F3/MAT1-1-1
KC437356 6650  GAGGCCGACGTTCTTACACCCGTGAGCACCGCAGCAGCATCTCGCGCCACTAGGCAGACC  6709
AGW27560 41     E  A  D  V  L  T  P  V  S  T  A  A  A  S  R  A  T  R  Q  T   60

     m1F3/MAT1-1-1→
KC437356 6710  AAAGAAGCATCTTGTGATCGAGCGAAGCGACCTCTCAATGCCTTCATGGCATTCCGAAGT  6769
AGW27560 61     K  E  A  S  C  D  R  A  K  R  P  L  N  A  F  M  A  F  R  S   80

KC437356 6770  ATGTTCATCCTCTTTCACGCACGTTGAAGTGGCTGACACGGACATTCTAGGTTACTACTT  6829
AGW27560 81                                                      Y  Y  L   83

KC437356 6830  GAAGCTGTTTCCCGACGTGCAGCAGAAGACCGCTTCTGGGTTTCTCACCACCTTGTGGCA  6889
AGW27560 84      K  L  F  P  D  V  Q  Q  K  T  A  S  G  F  L  T  T  L  W  H  103

KC437356 6890  CAAAGACCCGTTCAGAAACAAGTGGGCGCTGATTGCGAAGGTGTACTCCTTCGTGCGAGA  6949
AGW27560 104     K  D  P  F  R  N  K  W  A  L  I  A  K  V  Y  S  F  V  R  D  123

KC437356 6950  TCAGATTGGCAAGGACAAGGTTTCTCTATCATATTTCATGAGCCTTGCTTGTCCTACCAT  7009
AGW27560 124     Q  I  G  K  D  K  V  S  L  S  Y  F  M  S  L  A  C  P  T  M  143

KC437356 7010  GACCATCATCGAGCCCGCTGCGTACCTGAACGCGCTTGGGTGGTGTGTCCAAGATGACGA  7069
AGW27560 144     T  I  I  E  P  A  A  Y  L  N  A  L  G  W  C  V  Q  D  D  D  163

KC437356 7070  CGCTGGATCGCAGAAGCTATTCCAAGACGAATCTTCTGCAAACCTGGACCAGTCCAGCTT  7129
AGW27560 164     A  G  S  Q  K  L  F  Q  D  E  S  S  A  N  L  D  Q  S  S  L  183

KC437356 7130  GCTCTCGGCGGAATACCCCAGCACCGAAATCGAACTCTTGTCCGCCCTCGTCAACATTGG  7189
AGW27560 184     L  S  A  E  Y  P  S  T  E  I  E  L  L  S  A  L  V  N  I  G  203

KC437356 7190  GTACTTTCCCGATCACGGCGCCGACCTTGTGGAGAGAATGGGATCCAGCCACAGTGGCAT  7249
AGW27560 204     Y  F  P  D  H  G  A  D  L  V  E  R  M  G  S  S  H  S  G  I  223

KC437356 7250  CATGGCTCCGCGTGCCGCCAATTGCACTCCTCCAGTGTCCTACACGAAGGAAAAGATCGA  7309
AGW27560 224     M  A  P  R  A  A  N  C  T  P  P  V  S  Y  T  K  E  K  I  D  243

KC437356 7310  TTTCATCAACACAATCAGAAGCGATCCAGTTCAGGCGACAAAGGAGATCCTCGGTGATTG  7369
AGW27560 244     F  I  N  T  I  R  S  D  P  V  Q  A  T  K  E  I  L  G  D  C  263

KC437356 7370  CTACGATGAAACCACAATCAAGCTTCTGGGTGTCAAGTCACACAATGTGGAGAGTGTTGA  7429
AGW27560 264     Y  D  E  T  T  I  K  L  L  G  V  K  S  H  N  V  E  S  V  D  283

KC437356 7430  CTCCATCACGCACTTGTCCATGCAACGCGAATATCAGGCTCCGCGATTTTTTTATGACTA  7489
AGW27560 284     S  I  T  H  L  S  M  Q  R  E  Y  Q  A  P  R  F  F  Y  D  Y  303

KC437356 7490  CTCCGTCAGTTACGCAGGGATGGACTTCGGCGGTTCGAATGAGCCCGTGATGAACTTGAA  7549
AGW27560 304     S  V  S  Y  A  G  M  D  F  G  G  S  N  E  P  V  M  N  L  N  323

KC437356 7550  CAATCTCCCCGAGAACGAAACTTTCGACATCGACAGTCCTTTTGATCTCGATAAGATCCT  7609
AGW27560 324     N  L  P  E  N  E  T  F  D  I  D  S  P  F  D  L  D  K  I  L  343

KC437356 7610  TGGTCAATCGCAGTCAGAGGGCGAAAGAAGTGAGTGAAGCATCCCATTCACAGATGTTTT  7669
AGW27560 344     G  Q  S  Q  S  E  G  E  R                                   352

                                                             ← m1R3/MAT1-1-1
KC437356 7670  TAGCTAACGGCGCCTAGCTTCTCATCTTCCTCCGAGTCCTCCACACAACCCTCTGGACGA  7729
AGW27560 353                    T  S  H  L  P  P  S  P  P  H  N  P  L  D  D  367

               m1R3/MAT1-1-1  (stop codon)
KC437356 7730  CTTTTACTTTGCGTTCTAG                                           7748
AGW27560 368     F  Y  F  A  F                                               372
```

Figure S1. Alignment of the DNA sequence of the *MAT1-1-1* gene KC437356 (6530→7748) and the encoded sequence of the MAT1-1-1 protein AGW27560 (1→ 372) derived from *Ophiocordyceps sinensis* strain CS68-2-1229 [14]. Residues in pink indicate the MATα_HMGbox domain of the *MAT1-1-1* gene and MAT1-1-1 protein; those in green indicate introns I and II. The underlined residues in red indicate the forward primer m1F3/*MAT1-1-1* (CCACTAGGCAGACCAAAGAAG) and the reverse complementary primer m1R3/*MAT1-1-1* (CGCAAAGTAAAAGTCGTCCAGA) reported by [14]. The triplet residues (TAG) in red represent the stop codon.

```
                                                            Mat1-2F/MAT1-2-1 →
HM212637 3241  ATGGCCAATCCCATCAACATGATTCCCAATCCTCAGTGGAATGCGACTGACTACGAAGCG  3300
AEH27625 1      M  A  N  P  I  N  M  I  P  N  P  Q  W  N  A  T  D  Y  E  A   20

HM212637 3301  ATCTGGAAAGGCCTCGAGGCACAGGTCAATCCTTTCTCGCAGATTCTCTGCCTGGAGGGG  3360
AEH27625 21     I  W  K  G  L  E  A  Q  V  N  P  F  S  Q  I  L  C  L  E  G   40

HM212637 3361  GATTTTTTCCGCCAGCTCGACGATGCTGCGAAGCTGTTCATTGCTCGGAAGCTCATGTAA  3420
AEH27625 41     D  F  F  R  Q  L  D  D  A  A  K  L  F  I  A  R  K  L  M      59

HM212637 3421  GTTTGATGTTCACCGCAACTCGTCACACCTTCCTAATCGAATCTACAGGGAACACGTTCA  3480
AEH27625 60                                                  E  H  V  Q   63

HM212637 3481  GGAGTCAGTCCTGTATGTCAATGACGGCAATGGACCCGATCGCGTCTACCTTGGAGCTCC  3540
AEH27625 64     E  S  V  L  Y  V  N  D  G  N  G  P  D  R  V  Y  L  G  A  P   83

HM212637 3541  CCGACATTTTGTCGTTGGTGGTGGCATGATTCTCCAGATTTCTGGCTACGCGCCGTACTG  3600
AEH27625 84     R  H  F  V  V  G  G  G  M  I  L  Q  I  S  G  Y  A  P  Y  W   103

HM212637 3601  GATCCGACGTTCCGTGTCGAAAGTCGTTACTGCAACAGTGCTCGCGCCTCCCTCGCCCAA  3660
AEH27625 104    I  R  R  S  V  S  K  V  V  T  A  T  V  L  A  P  P  S  P  K   123

HM212637 3661  GGATATCAAGATCCCTCGTCCTCCCAACGCGTACATCTTGTACCGTAAGGAGCGCCACCA  3720
AEH27625 124    D  I  K  I  P  R  P  P  N  A  Y  I  L  Y  R  K  E  R  H  H   143

HM212637 3721  TTATGTCAAGGATGCAAATCCTGGCATTACGAACAACGAGATTTGTAAGTTTCATAGCCG  3780
AEH27625 144    Y  V  K  D  A  N  P  G  I  T  N  N  E  I                     157

HM212637 3781  CTCCTCCTTTCTGCCATCGTGTTCTAATGTCTCTCTCAGCCCAAATCTTGGGCAAAGCTT  3840
AEH27625 158                                           S  Q  I  L  G  K  A    164

                                                                ← Mat1-2R
HM212637 3841  GGAACATGGAGTCGAACGACGTCAGACAGAAGTACAAGGACATGTCTCAGCAAGTCAAGC  3900
AEH27625 165   W  N  M  E  S  N  D  V  R  Q  K  Y  K  D  M  S  Q  Q  V  K    184

               /MAT1-2-1
HM212637 3901  AAGCTCTCCTGGAGAAGCACCCAGACTACCAGTACAAACCGCGTCGTCCTTGCGAGCGCC  3960
AEH27625 185   Q  A  L  L  E  K  H  P  D  Y  Q  Y  K  P  R  R  P  C  E  R    204

HM212637 3961  GGCGCCGTCGTCGTGCCAGTCCGAACCAAAACCCGAAGCAATCTACGTCGAGAAATGCCG  4020
AEH27625 205   R  R  R  R  R  A  S  P  N  Q  N  P  K  Q  S  T  S  R  N  A    224

HM212637 4021  CTACTAGGGACGCCGCGATCTCGAGTGAAGATACTTCTACTGCCACCGGAGATACCAACA  4080
AEH27625 225   A  T  R  D  A  A  I  S  S  E  D  T  S  T  A  T  G  D  T  N    244

                              (stop codon)
HM212637 4081  CTGCGAATGGTTTCTAA                                             4097
AEH27625 245   T  A  N  G  F                                                 249
```

Figure S2. Alignment of the *MAT1-2-1* gene HM212637 (3241→4097) and the encoded sequence of the MAT1-2-1 protein AEH27625 (1→249) derived from *Ophiocordyceps sinensis* strain CS2 [70]. Residues in pink indicate the HMG-box_ROX1-like domain of the *MAT1-2-1* gene and MAT1-2-1 proteins; those in green indicate introns I and II. The underlined residues in red indicate the forward primer *Mat1-2F*/*MAT1-2-1* (TGGAATGCGACTGACTACGA) and the reverse complementary primer *Mat1-2R*/*MAT1-2-1* (CCAGGAGAGCTTGCTTGACT) reported previously [14,70]. The triplet residues (TAA) in red represent the stop codon.

```
AEH27625 1     MANPINMIPNPQWNATDYEAIWKGLEAQVNPFSQILCLEGDFFRQLDDAAKLFIARKLME  60
AGW27538 1     ------------------------------------------------------------  60
AGW27541 1     ------------------------------------------------------------  60
AGW27539 1     ------------------------------------------------------------  60
AGW27556 1     ------------------------------------------------------------  60
AGW27547 1     ------------------------------------------------------------  60
AGW27546 1     ------------------------------------------------------------  60
AGW27545 1     ------------------------------------------------------------  60
AGW27542 1     ------------------------------------------------------------  60
AGW27541 1     ------------------------------------------------------------  60
AGW27549 1     ------------------------------------------------------------  60
AGW27550 1     ------------------------------------------------------------  60
AGW27551 1     ------------------------------------------------------------  60
AGW27544 1     ------------------------------------------------------------  60
AGW27540 1     ------------------------------------------------------------  60
AGW27537 1     ------------------------------------------------------------  60
AGW27553 1     ------------------------------------------------------------  60

AEH27625 61    HVQESVLYVNDGNGPDRVYLGAPRHFVVGGGMILQISGYAPYWIRRSVSKVVTATVLAPP  120
AGW27538 61    ------------------------------------------------------------  120
AGW27541 61    ------------------------------------------------------------  120
AGW27539 61    ------------------------------------------------------------  120
AGW27556 61    ------------------------------------------------------------  120
AGW27547 61    ------------------------------------------------------------  120
AGW27546 61    ------------------------------------------------------------  120
AGW27545 61    ------------------------------------------------------------  120
AGW27542 61    ------------------------------------------------------------  120
AGW27541 61    ------------------------------------------------------------  120
AGW27549 61    ------------------------------------------------------------  120
AGW27550 61    ------------------------------------------------------------  120
AGW27551 61    ------------------------------------------------------------  120
AGW27544 61    ------------------------------------------------------------  120
AGW27540 61    ------------------------------------------------------------  120
AGW27537 61    -----------------I------------------------------------------  120
AGW27553 61    ------------------------------------------------------------  120

AEH27625 121   SPKDIKIPRPPNAYILYRKERHHYVKDANPGITNNEISQILGKAWNMESNDVRQKYKDMS  180
AGW27538 121   ------------------------------------------------------------  180
AGW27541 121   ------------------------------------------------------------  180
AGW27539 121   ------------------------------------------------------------  180
AGW27556 121   -----------------------H------------------------------------  180
AGW27547 121   -----------------------H------------------------------------  180
AGW27546 121   -----------------------H------------------------------------  180
AGW27545 121   -----------------------H------------------------------------  180
AGW27542 121   -----------------------H------------------------------------  180
AGW27541 121   -----------------------H------------------------------------  180
AGW27549 121   -----------------------H------------------------------------  180
AGW27550 121   -----------------------H------------------------------------  180
AGW27551 121   -----------------------H------------------------------------  180
AGW27544 121   -----------------------H------------------------------------  180
AGW27540 121   -----------------------H------------------------------------  180
AGW27537 121   -----------------------H------------------------------------  180
AGW27553 121   -----------------------H------------------------------X-----  180

AEH27625 181   QQVKQALLEKHPDYQYKPRRPCERRRRRRASPNQNPKQSTSRNAATRDAAISSEDTSTAT  240
AGW27538 181   ------------------------------------------------------------  240
AGW27541 181   ------------------------------------------------------------  240
AGW27539 181   ------------------------------------------------------------  240
AGW27556 181   ------------------------------------------------------------  240
AGW27547 181   ------------------------------------------------------------  240
AGW27546 181   ------------------------------------------------------------  240
AGW27545 181   ------------------------------------------------------------  240
AGW27542 181   ------------------------------------------------------------  240
AGW27541 181   ------------------------------------------------------------  240
AGW27549 181   ------------------------------------------------------------  240
AGW27550 181   ------------------------------------------------------------  240
AGW27551 181   ------------------------------------------------------------  240
AGW27544 181   ------------------------------------------------------------  240
AGW27540 181   ------------------------------------------------------------  240
AGW27537 181   ------------------------------------------------------------  240
AGW27553 181   ------------------------------------------------------------  240

AEH27625 241   GDTNTANGF                                                     249
AGW27538 241   ---------                                                     249
AGW27541 241   ---------                                                     249
AGW27539 241   ---------                                                     249
AGW27556 241   ---------                                                     249
AGW27547 241   ---------                                                     249
AGW27546 241   ---------                                                     249
AGW27545 241   ---------                                                     249
AGW27542 241   ---------                                                     249
AGW27541 241   ---------                                                     249
AGW27549 241   ---------                                                     249
AGW27550 241   ---------                                                     249
AGW27551 241   ---------                                                     249
AGW27544 241   ---------                                                     249
AGW27540 241   ---------                                                     249
AGW27537 241   ---------                                                     249
AGW27553 241   --------                                                      248
```

Figure S3. Alignment of the sequences of full-length MAT1-2-1 proteins derived from Group II *O. sinensis* strains listed in Table 2 with that of the authentic MAT1-2-1 protein AEH27625 (under the AlphaFold code D7F2E9) derived from the *H. sinensis* strain CS2. The residues in pink indicate the HMG-box_ROX1-like domains of the proteins, those in green indicate the variant amino acids, the hyphens indicate identical amino acid residues, and the space denotes the unmatched sequence gap.

**Panel (A): Alignment of amino acid sequences of the MATα_HMGbox domains**

```
AGW27560  42   ADVLTPVSTAAASRATRQTKEASCDRAKRPLNAFMAFRSYYLKLFPDVQQKTASGFLTT  100
AGW27517  1                         -------------------------------------  37
AGW27518  1                         -------------------------------------  37
AGW27519  1                         -------------------------------------  37
AGW27520  1                         -------------------------------------  37
AGW27521  1                         -------------------------------------  37
AGW27523  1                         -------------------------------------  37
AGW27524  1                         -------------------------------------  37
AGW27528  1                         -------------------------------------  37

AGW27560  101 LWHKDPFRNKWALIAKVYSFVRDQIGKDKVSLSYFMSLACPTMTIIEPAAYLNALGWCVQ  160
AGW27517  38  ------------------------------------------------------------  97
AGW27518  38  ------------------------------------------------------------  97
AGW27519  38  ------------------------------------------------------------  97
AGW27520  38  ------------------------------------------------------------  97
AGW27521  38  ------------------------------------------------------------  97
AGW27523  38  ------------------------------------------------------------  97
AGW27524  38  ------------------------------------------------------------  97
AGW27528  38  ------------------------------------------------------------  97

AGW27560  161 DDDAGSQKLFQDESSANLDQSSLLSAEYPSTEIELLSALVNIGYFPDHGADLVERMGSSH  220
AGW27517  98  ------------------------------------------------------------  157
AGW27518  98  ------------------------------------------------------------  157
AGW27519  98  ------------------------------------------------------------  157
AGW27520  98  ------------------------------------------------------------  157
AGW27521  98  ------------------------------------------------------------  157
AGW27523  98  ------------------------------------------------------------  157
AGW27524  98  ------------------------------------------------------------  157
AGW27528  98  ------------------------------------------------------------  157

AGW27560  221 SGIMAPRAANCTPP                                                234
AGW27517  158 --------------                                                171
AGW27518  158 --------------                                                171
AGW27519  158 --------------                                                171
AGW27520  158 --------------                                                171
AGW27521  158 --------------                                                171
AGW27523  158 --------------                                                171
AGW27524  158 --------------                                                171
AGW27528  158 --------------                                                171
```

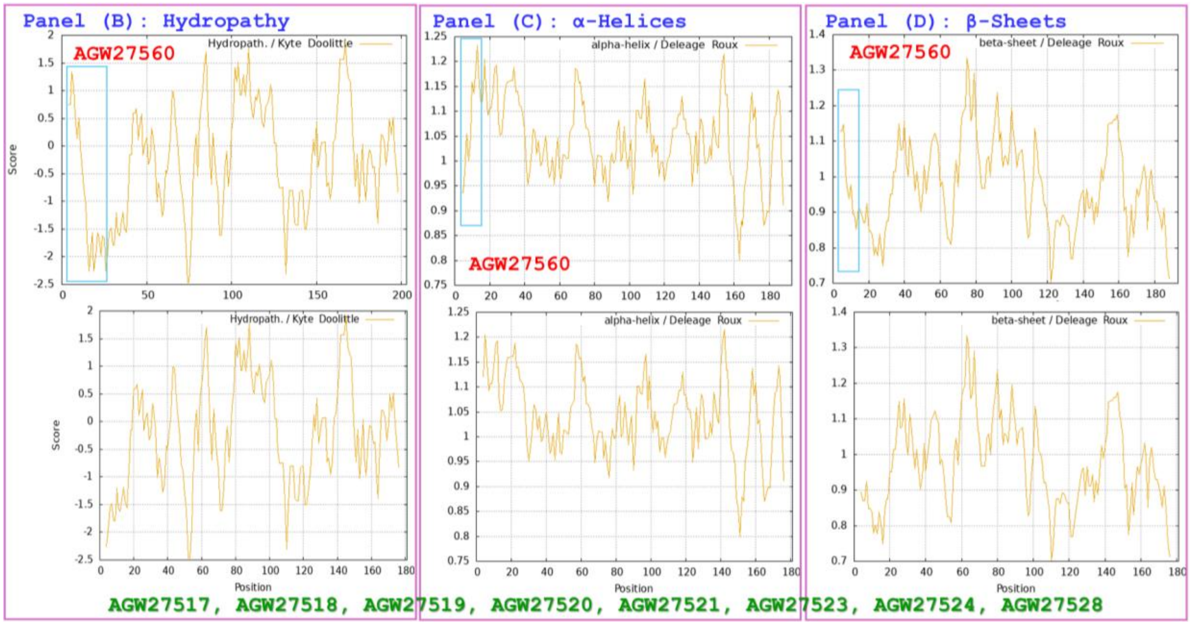

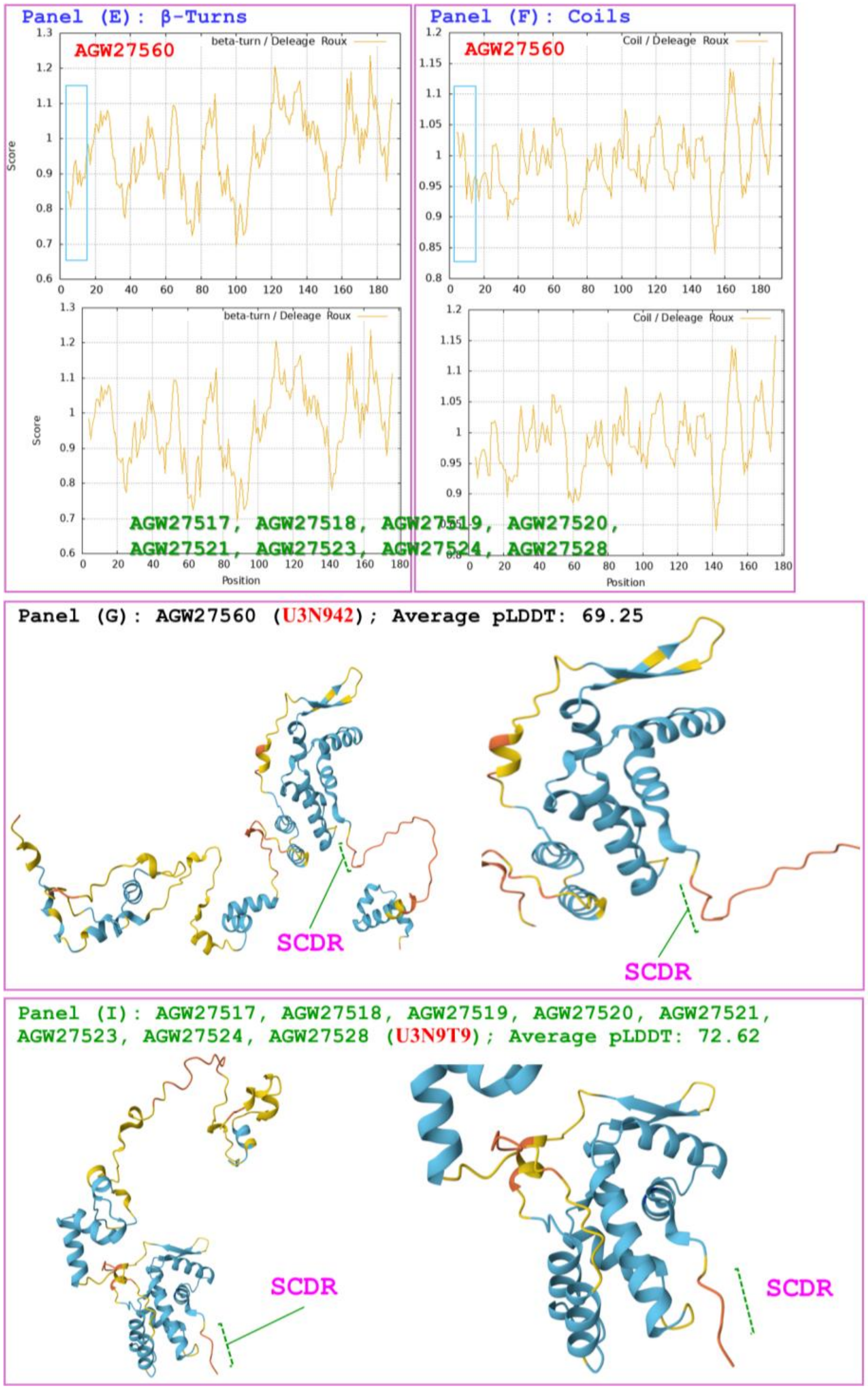


Figure S4x5. Correlations of the changes in the hydrophobicity and the primary, secondary, and tertiary structures of the MATα_HMGbox domains of MAT1-1-1 proteins: the reference protein AGW27560 (under the AlphaFold code U3N942) derived from H. sinensis strain CS68-2-1229 and the MAT1-1-1 truncated proteins (under the AlphaFold code U3N9T9) derived from the O. sinensis strains CS6-251, CS36-1294, CS37-295, CS18-266, CS26-277, CS91-1291, CS34-291, and CS68-2-1229. Panel (A) shows an alignment of the amino acid sequences of the MATα_HMGbox domains of the MAT1-1-1 proteins, where the hyphens indicate identical amino acid residues and the spaces denote unmatched sequence gaps. The ExPASy ProtScale plots show the changes in hydrophobicity and the 2D structures in Panels (B)–(F) for hydropathy, α-helices, β-sheets, β-turns, and coils of the proteins, respectively; the open rectangles in blue highlight the N-terminally truncated region in the ExPASy plots. Panels (G)–(I) show the 3D structures of the full-length proteins on the left, and the locally magnified structures surrounding the truncated region are shown on the right. The model confidence for the AlphaFold-predicted 3D structures is as follows: very high (pLDDT>90); high (90>pLDDT>70); low (70>pLDDT>50); and very low (pLDDT<50).

**Table S1**. The *O. sinensis* strains, GenBank accession numbers for the ITS nucleic acid sequences corresponding to the GenBank accession numbers for the MAT1-1-1 and MAT1-2-1 proteins, and percent similarities *vs*. GC-biased Genotypes #1–3, #7–10, and #12 of *O. sinensis*.

| ***O. sinensis* strain** | **ITS1-5.8S-ITS2** | **Percent similarity between the sequences of ITS and the reference GC-biased Genotypes of *O. sinensis*** | | | | | | | |
|---|---|---|---|---|---|---|---|---|---|
| | | **#1** | **#2** | **#3** | **#7** | **#8** | **#9** | **#10** | **#12** |
| CS6-251 | JQ900163 | **98.2%** | **97.2%** | 93.6% | 93.2% | 88.6% | 93.9% | 81.8% | 93.1% |
| | JQ900180 | 87.3% | 86.9% | 85.9% | 84.3% | 83.2% | 85.1% | 76.1% | 83.5% |
| CS34-291 | JQ900162 | **100%** | **97.2%** | 95.3% | 95.0% | 89.5% | 95.2% | 83.1% | 94.4% |
| | JQ900179 | 87.3% | 86.9% | 85.9% | 84.3% | 83.2% | 85.1% | 76.1% | 83.5% |
| CS70-1208 | JQ900146 | **100%** | **97.2%** | 95.3% | 95.0% | 89.5% | 95.2% | 83.1% | 94.4% |
| | JQ900172 | 87.1% | 86.3% | 85.6% | 84.3% | 83.2% | 85.3% | 76.7% | 83.7% |
| CS71-1220 | JQ900152 | **100%** | **97.2%** | 95.3% | 95.0% | 89.5% | 95.2% | 83.1% | 94.4% |
| | JQ900175 | 87.1% | 86.3% | 85.6% | 84.3% | 83.2% | 85.3% | 76.7% | 83.7% |
| CS68-2-1228 | JQ900157 | **100%** | **97.2%** | 95.3% | 95.0% | 89.5% | 95.2% | 83.1% | 94.4% |
| | JQ900178 | 87.1% | 86.3% | 85.6% | 84.3% | 83.2% | 85.3% | 76.7% | 83.7% |
| CS76-1284 | JQ900159 | **100%** | **97.2%** | 95.3% | 95.0% | 89.5% | 95.2% | 83.1% | 94.4% |
| CS68-2-1229 | JQ900158 | **100%** | **97.2%** | 95.3% | 95.0% | 89.5% | 95.2% | 83.1% | 94.4% |
| CS71-1218 | JQ900150 | **100%** | **97.2%** | 95.3% | 95.0% | 89.5% | 95.2% | 83.1% | 94.4% |
| CS71-1219 | JQ900151 | **100%** | **97.2%** | 95.3% | 95.0% | 89.5% | 95.2% | 83.1% | 94.4% |
| CS560-961 | JQ900143 | **99.8%** | 96.8% | 95.1% | 94.8% | 89.3% | 95.0% | 82.9% | 94.2% |
| CS561-964 | JQ900144 | **99.8%** | 96.8% | 95.1% | 94.7% | 89.3% | 95.0% | 82.9% | 94.2% |
| CS18-266 | JQ900168 | **98.9%** | **97.2%** | 95.5% | 94.0% | 90.6% | 95.7% | 83.9% | 95.1% |
| CS25-273 | JQ900167 | **98.9%** | **97.2%** | 95.5% | 94.0% | 90.6% | 95.7% | 83.9% | 95.1% |
| CS91-1291 | JQ900166 | **98.8%** | 96.8% | 95.3% | 93.7% | 90.4% | 95.5% | 83.7% | 94.9% |
| CS37-295 | JQ900169 | **98.4%** | 96.8% | 95.1% | 94.0% | 90.4% | 95.5% | 83.5% | 95.1% |

Note: The percentage numbers in red refer to the similarities >97% between the sequences of ITS and reference GC-biased Genotypes #1–3, #7–10, and 12 of *O. sinensis*: Genotypes #1 (AB067721), #2 (MG770309), #3 (HM595984), #7 (AJ488254), #8 (GU246286), #9 (GU246288), #10 (GU246287), and #12 (GU246296) [9–10].

**Table S2**. The *O. sinensis* strains, GenBank accession numbers for the ITS nucleic acid sequences corresponding to the GenBank accession numbers for the MAT1-1-1 and MAT1-2-1 proteins, and percent similarities *vs*. AT-biased Genotypes #4–6 and #15–17 of *O. sinensis*.

| ***O. sinensis* strain** | **ITS1-5.8S-ITS2** | **Percent similarity comparing the sequences of ITS and the reference AT-biased genotypes of *O. sinensis*** | | | | | |
|---|---|---|---|---|---|---|---|
| | | **#4** | **#5** | **#6** | **#15** | **#16** | **#17** |
| CS6-251 | JQ900163 | 87.5% | 85.4% | 84.2% | 88.4% | 86.7% | 86.8% |
| | JQ900180 | 89.9% | **97.2%** | 92.3% | 93.3% | 93.1% | **98.3%** |
| CS34-291 | JQ900162 | 88.6% | 86.1% | 85.5% | 89.5% | 87.2% | 87.5% |
| | JQ900179 | 89.9% | **97.2%** | 92.3% | 93.3% | 93.1% | **98.3%** |
| CS70-1208 | JQ900146 | 88.6% | 86.1% | 85.5% | 89.5% | 87.2% | 87.5% |
| | JQ900172 | 90.2% | **97.6%** | 93.4% | 93.5% | 93.3% | **98.1%** |
| CS71-1220 | JQ900152 | 88.6% | 86.1% | 85.5% | 89.5% | 87.2% | 87.5% |
| | JQ900175 | 90.2% | **97.6%** | 93.4% | 93.5% | 93.3% | **98.1%** |
| CS68-2-1228 | JQ900157 | 88.6% | 86.1% | 85.5% | 89.5% | 87.2% | 87.5% |
| | JQ900178 | 90.2% | **97.6%** | 93.4% | 93.5% | 93.3% | **98.1%** |
| CS76-1284 | JQ900159 | 88.6% | 86.1% | 85.5% | 89.5% | 87.2% | 87.5% |
| CS68-2-1229 | JQ900158 | 88.6% | 86.1% | 85.5% | 89.5% | 87.2% | 87.5% |
| CS71-1218 | JQ900150 | 88.6% | 86.1% | 85.5% | 89.5% | 87.2% | 87.5% |
| CS71-1219 | JQ900151 | 88.6% | 86.1% | 85.5% | 89.5% | 87.2% | 87.5% |
| CS560-961 | JQ900143 | 88.4% | 85.9% | 85.2% | 89.3% | 87.0% | 87.4% |
| CS561-964 | JQ900144 | 88.4% | 85.9% | 85.2% | 89.3% | 87.0% | 87.4% |
| CS18-266 | JQ900168 | 88.3% | 87.2% | 85.5% | 90.6% | 86.8% | 88.6% |
| CS25-273 | JQ900167 | 88.3% | 87.2% | 85.5% | 90.6% | 86.8% | 88.6% |
| CS91-1291 | JQ900166 | 88.3% | 87.0% | 85.2% | 90.4% | 86.7% | 88.4% |
| CS37-295 | JQ900169 | 87.7% | 86.7% | 86.0% | 90.0% | 86.3% | 88.1% |

Note: The percentage numbers in red refer to the similarities >97% between the sequences of ITS and reference AT-biased Genotypes #4–6 and #15–17 of *O. sinensis*: Genotypes #4 (AB067744), #5 (AB067740), #6 (KJ720572), #15 (KT232017), #16 (KT232019), #and #17 (KT232010) [9−10].

**Table S3**. Amino acid scales based on the general chemical characteristics of their side chains for ProtScale analysis (https://web.expasy.org/protscale/) to predict the hydrophobicity and secondary structures (α-helices, β-sheets, β-turns, and coils) of proteins.

| Amino acid | | hydropathy index * | α-Helix | β-Sheet | β-Turn | Coil | Chemical-physical property |
|---|---|---|---|---|---|---|---|
| Asp, D | Aspartic acid | -3.500 | 0.924 | 0.541 | 1.197 | 1.197 | Acidic |
| Glu, E | Glutamic acid | -3.500 | 1.504 | 0.567 | 1.149 | 0.761 | Acidic |
| Ile, I | Isoleucine | 4.500 | 1.003 | 1.799 | 0.240 | 0.886 | Aliphatic |
| Val, V | Valine | 4.200 | 0.990 | 1.965 | 0.387 | 0.772 | Aliphatic |
| Leu, L | Leucine | 3.800 | 1.236 | 1.261 | 0.670 | 0.810 | Aliphatic |
| Ala, A | Alanine | 1.800 | 1.489 | 0.709 | 0.788 | 0.824 | Aliphatic |
| Phe, F | Phenylalanine | 2.800 | 1.195 | 1.393 | 0.624 | 0.797 | Aromatic |
| Trp, W | Tryptophan | -0.900 | 1.090 | 1.306 | 0.546 | 0.941 | Aromatic |
| Tyr, Y | Tyrosine | -1.300 | 0.787 | 1.266 | 0.795 | 1.109 | Aromatic |
| His, H | Histidine | -3.200 | 1.003 | 0.863 | 0.970 | 1.068 | Basic |
| Lys, K | Lysine | -3.900 | 1.172 | 0.721 | 1.302 | 0.897 | Basic |
| Arg, R | Arginine | -4.500 | 1.224 | 0.920 | 0.912 | 0.893 | Basic |
| Cys, C | Cysteine | 2.500 | 0.966 | 1.191 | 0.965 | 0.953 | with polar neutral side chains |
| Met, M | Methionine | 1.900 | 1.363 | 1.210 | 0.436 | 0.810 | with polar neutral side chains |
| Ser, S | Serine | -0.800 | 0.739 | 0.928 | 1.316 | 1.130 | with polar neutral side chains |
| Thr, T | Threonine | -0.700 | 0.785 | 1.221 | 0.739 | 1.148 | with polar neutral side chains |
| Asn, N | Asparagine | -3.500 | 0.772 | 0.604 | 1.572 | 1.167 | with polar neutral side chains |
| Gln, Q | Glutamine | -3.500 | 1.164 | 0.840 | 0.997 | 0.947 | with polar neutral side chains |
| Gly, G | Glycine | -0.400 | 0.510 | 0.657 | 1.860 | 1.251 | Unique amino acid |
| Pro, P | Proline | -1.600 | 0.492 | 0.354 | 1.415 | 1.540 | Unique amino acid |

Note: An amino acid scale is defined at https://web.expasy.org/protscale/ by a numerical value assigned to each type of amino acid. The most frequently used scales are the hydrophobicity or hydrophilicity scales and the secondary structure conformational parameter scales, but many other scales exist, which are based on the different chemical and physical properties of the amino acids. The ExPASy ProtScale program provides 57 predefined scales based on the literature [74]. *, Hydropathy index [73]; the larger the value, the stronger the hydrophobicity; negative values indicate hydrophilicity.

**Table S4**. GenBank accession numbers (in red in parentheses) for the full-length MAT1-1-1 proteins in the AlphaFold database under the corresponding AlphaFold UniProt codes [87].

| AlphaFold UniProt code | Strain/isolate number (GenBank accession number for MAT1-1-1 protein) |
|---|---|
| U3N942 | CS68-2-1229 (AGW27560), GS09_111 (ALH24945), GS09_131 (ALH24947), ID10_1 (ALH24954), IOZ07 (KAF4512729), NP10_1 (ALH24955), NP10_2 (ALH24956), QH07_188 (ALH24957), QH09_122 (ALH24959), QH09_131 (ALH24960), QH09_151 (ALH24961), QH09_20L (ALH24965), QH09_33L (ALH24967), QH09_37 (ALH24968), QH09_46 (ALH24969), QH09_56 (ALH24970), QH09_66 (ALH24971), QH09_78 (ALH24972), QH09_93 (ALH24973), QH10_1 (ALH24974), QH10_4 (ALH24975), QH10_7 (ALH24976), SC09_107 (ALH24978), SC09_117 (ALH24979), SC09_128 (ALH24980), SC09_147 (ALH24981), SC09_157 (ALH24982), SC09_167 (ALH24983), SC09_180 (ALH24984), SC09_190 (ALH24985), SC09_200 (ALH24986), SC09_21 (ALH24987), SC09_36 (ALH24988), SC09_37 (ALH24989), SC09_47 (ALH24990), SC09_57 (ALH24991), SC09_77 (ALH24993), SC10_18 (ALH24996), SC10_21 (ALH24997), SC10_4 (ALH24998), XZ05_12 (ALH25000), XZ05_3 (ALH25002), XZ05_7 (ALH25004), XZ06_124 (ALH25006), XZ06_152 (ALH25007), XZ07_108 (ALH25009), XZ07_133 (ALH25010), XZ07_154 (ALH25011), XZ07_166 (ALH25012), XZ07_176 (ALH25013), XZ07_180 (ALH25014), XZ08_10 (ALH25015), XZ08_24 (ALH25016), XZ08_26 (ALH25017), XZ08_4 (ALH25018), XZ08_56 (ALH25019), XZ08_59 (ALH25020), XZ08_A1 (ALH25021), XZ08_B1 (ALH25022), XZ09_106 (ALH25024), XZ09_113 (ALH25025), XZ09_118 (ALH25026), XZ09_15 (ALH25027), XZ09_32 (ALH25028), XZ09_4 (ALH25029), XZ09_46 (ALH25030), XZ09_48 (ALH25031), XZ09_59 (ALH25032), XZ09_71 (ALH25033), XZ09_80 (ALH25055), XZ10_15 (ALH25035), XZ10_17 (ALH25036), XZ10_23 (ALH25037), XZ10_7 (ALH25038), XZ12_1 (ALH25056), XZ12_33 (ALH25058), XZ12_43 (ALH25059), YN07_6 (ALH25039), YN07_8 (ALH25040), YN09_101 (ALH25041), YN09_140 (ALH25042), YN09_3 (ALH25044), YN09_72 (ALH25049), YN09_81 (ALH25050), YN09_85 (ALH25051), YN09_89 (ALH25052), YN09_96 (ALH25053) |
| A0A0N9QMM1 | GS09_121 (ALH24946), GS09_201 (ALH24949), GS09_225 (ALH24950), SC09_1 (ALH24977) |
| T5A511 | Co18 (EQK97643) (KE657544 410←1519) (ANOV01017390 410←1519) |
| A0A0N9R5B3 | SC09_65 (ALH24992) |
| A0A0N7G849 | SC09_97 (ALH24995) |
| A0A0N9QUF3 | GS09_143 (ALH24948) |
| A0A0N9R4V2 | YN09_61 (ALH25047) |
| A0A0N9QMS9 | YN09_22 (ALH25043), YN09_51 (ALH25045), YN09_6 (ALH25046), YN09_64 (ALH25048) |
| A0A0N7G845 | GS09_229 (ALH24951), GS09_281 (ALH24952), GS09_311 (ALH25054), GS10_1 (ALH24953), QH09_164 (ALH24962), QH09_173 (ALH24963), QH09_201 (ALH24964), QH09_210 (ALH24966), SC09_87 (ALH24994) |
| A0A0N9QUK2 | XZ05_8 (ALH25005) |
| A0A0N9QMT4 | XZ07_H2 (ALH24999), XZ12_16 (ALH25057) |
| A0A0N9QMR3 | XZ06_260 (ALH25008), XZ09_100 (ALH25023) |
| A0A0N9QMS4 | XZ09_95 (ALH25034) |
| A0A0N7G850 | XZ05_6 (ALH25003) |
| A0A0N9R4Q4 | XZ05_2 (ALH25001) |

Note: *, Branch 1 in red, Branch 2 in pink, Branch 3 in purple, and Branch 4 in brown under the cluster codes (English letters) in the parentheses were determined *via* the Bayesian analysis shown in Figure 1 of [69]. The "←" arrows indicate sequences in the antisense strands of the genome of the *H. sinensis* strain Co18.

**Table S5**. GenBank accession numbers (in red) for the full-length MAT1-2-1 proteins of 69 *H. sinensis* strains or wild-type *C. sinensis* isolates under the corresponding AlphaFold UniProt codes [87].

| AlphaFold UniProt code | Strain/isolate number (GenBank accession number for MAT1-2-1 protein) |
| --- | --- |
| D7F2E9 | CS2 (AEH27625) (ACV60400), SC-2 (ACV60395), SC-4 (ACV60396), SC-5 (ACV60398), SC-7 (ACV60397), XZ-LZ06-1 (ACV60369), XZ-LZ06-108 (ACV60373), XZ-LZ06-21 (ACV60371), XZ-LZ06-7 (ACV60370), XZ-LZ07-108 (ACV60379), XZ-LZ07-30 (ACV60377), XZ-ML-191 (ACV60376), YN-1 (ACV60390), YN-5 (ACV60392), YN-6 (ACV60393), SC09_77 (AFX66426), YN-8 (ACV60394), SC09_47 (AFX66423), SC09_57 (AFX66424), SC09_97 (AFX66428), XZ05_12 (AFX66444), XZ05_7 (AFX66442), XZ06_152 (AFX66445), XZ07_11 (AFX66447), XZ07_46 (AFX66448), XZ09_106 (AFX66464), XZ09_113 (AFX66465), XZ09_15 (AFX66455), YN09_101 (AFX66482), YN09_72 (AFX66477), YN09_81 (AFX66478), YN09_85 (AFX66479), YN09_89 (AFX66480), SC09-37 (AFH35019), CS26-277 (AGW27541), CS36-1294 (AGW27538), CS37-295 (AGW27539) |
| T5AF56 | Co18 (EQL04085) (ANOV01000063 9329→10182) |
| V9LW10 | SC09_200 (AFX66437) |
| D7F2H1 | YN-4 (ACV60391) |
| D7F2F2 | XZ-LZ06-61 (ACV60372) |
| A0A0A0RCF5 | XZ12_16 (AIV43040) |
| D7F2J7 | XZ-LZ07-H1 (ACV60417), XZ-LZ07-H2 (ACV60418), XZ06-124 (AFH35020), XZ05_8 (AFX66443) |
| D7F2F5 | XZ-LZ05-6 (ACV60415), XZ-SN-44 (ACV60375), XZ05_2 (AFX66441), XZ06_260 (AFX66446), XZ09_100 (AFX66463), XZ09_80 (AFX66461), XZ09_95 (AFX66462) |
| V9LWC9 | YN09_64 (AFX66476) |
| V9LVS8 | YN09_6 (AFX66472), YN09_22 (AFX66473), YN09_51 (AFX66474) |
| D7F2E3 | XZ-NQ-154 (ACV60363), XZ-NQ-155 (ACV60364), GS09_111 (AFX66388), QH09-93 (AFH35018), CS560-961 (AGW27542) |
| U3N9X0 | CS71-1218 (AGW27553) |
| D7F2G5 | QH-YS-199 (ACV60385) |
| D7F2H9 | SC-3 (ACV60399) |
| V9LW71 | QH09_11 (AFX66401) |
| V9LVU8 | YN09_61 (AFX66475) |
| V9LWG5 | ID10_1 (AFX66484) |
| U3N6V5 | CS6-251 (AGW27537) |
| ‡ | CS18-266 (AGW27540), CS76-1284 (AGW27545), CS561-964 (AGW27546), CS25-273 (AGW27547), CS34-291 (AGW27544), CS70-1208 (AGW27549), CS68-2-1228 (AGW27556), CS70-1211 (AGW27550), CS70-1212 (AGW27551), NP10_1 (AFX66485), NP10_2 (AFX66486), YN09_3 (AFX66471), YN09_96 (AFX66481), YN09_140 (AFX66483) |

Note: **, Branch 1 in red and Branch 2 in pink under the cluster codes (Roman numerals) in the paratheses were determined *via* the Bayesian analysis shown in Figure 2 of [69]. ‡, The 14 MAT1-2-1 protein sequences in green are included in the GenBank database, but their 3D structures are absent in the AlphaFold database. The "→" arrow indicates the sequence in the sense strand of the genome of the *H. sinensis* strain Co18.

**Table** S6. Similarity of the 3 pairs of primers that were used by Li *et al.* [36] to amplify the ITS sequences of the *O. sinensis* strains listed in Table 1 that were isolated either from the caterpillar body specimens that were collected from various production areas on the Qinghai–Tibet Plateau or from the cultures of mono-ascospores that were collected from mature *C. sinensis* insect–fungal complexes obtained from a single production region (Maqên, Guoluo, Qinghai Provence of China).

| Accession # for *O. sinensis* genotypes | | Percent identity/percent coverage of primer sequences aligned with the sequences of *O. sinensis* genotypes | | | | | | | |
|---|---|---|---|---|---|---|---|---|---|
| | | | | Specific for Group A ITS sequences | | Specific for Group C ITS sequences | | for amplifying 5.8S gene sequences | |
| | | ITS5(F) | ITS4(R) | GAF | GAR | GCF | GCR | 5.8S-F | 5.8S-R |
| #1 | AB067721 | **100%/100%** | **100%/100%** | **100%/100%** | **100%/100%** | 92%/89% | 93%/56% | 95%/100% | 93%/95% |
| #2 | MG770309 | —/— | —/— | —/— | —/— | —/— | —/— | 95%/100% | 93%/64% |
| #3 | HM595984 | —/— | 45%/100% | 86%/100% | 96%/100% | 92%/95% | —/— | 95%/100% | 93%/64% |
| #7 | AJ488254 | —/— | 35%/100% | 100%/68% | 100%/48% | 100%/68% | —/— | 95%/100% | 93%/64% |
| #8 | GU246286 | —/— | —/— | 100%/77% | 81%/100% | 88%/93% | 85%/80% | 100%/62% | 93%/82% |
| #9 | GU246288 | —/— | 40%/100% | **100%/100%** | 100%/48% | 100%/43% | 100%/40% | 94%/85% | 93%/68% |
| #10 | GU246287 | —/— | —/— | 95%/100% | 100%/74% | 100%/25% | 100%/28% | 94%/81% | 94%/73% |
| #11 | JQ695935 | —/— | **100%/100%** | 100%/55% | 100%/48% | 92%/89% | —/— | 95%/100% | 93%/95% |
| #12 | GU246296 | —/— | 100%/95% | **100%/100%** | 81%/91% | 92%/89% | 100%/40% | 95%/100% | 93%/95% |
| #13 | KT339190 | **100%/100%** | **100%/100%** | **100%/100%** | 81%/100% | 92%/89% | 100%/36% | 95%/100% | 100%/73% |
| #14 | KT339178 | **100%/100%** | **100%/100%** | 100%/78% | **100%/100%** | 100%/50% | 93%/56% | 95%/100% | 93%/95% |
| #4 | AB067744 | **100%/100%** | **100%/100%** | 100%/64% | 87%/63% | 92%/95% | 100%/68% | 95%/100% | 100%/82% |
| #5 | AB067740 | 100%/95% | —/— | 100%/41% | 100%/52% | **100%/100%** | **100%/100%** | **100%/100%** | 95%/100% |
| #6 | KJ720572 | 32%/100% | 35%/100% | 92%/55% | 100%/26% | **100%/100%** | 100%/60% | **100%/100%** | 100%/50% |
| #15 | KT232017 | 32%/100% | **100%/100%** | 100%/64% | 100%/52% | 92%/96% | **100%/100%** | 95%/100% | **100%/100%** |
| #16 | KT232019 | 32%/100% | **100%/100%** | 100%/64% | 86%/78% | **100%/100%** | 94%/68% | **100%/100%** | 100%/82% |
| #17 | KT232010 | 32%/100% | **100%/100%** | 100%/64% | 100%/52% | **100%/100%** | **100%/100%** | **100%/100%** | 95%/100% |

Note: Four pairs of primers ITS5(F)/ITS4(R) in purple, GAF/GCF in green, GCF/GCR in blue, and 5.8S-F/R in pink were designed/selected and used by Li *et al.* [36] for amplification of ITS sequences for phylogenetic and genotypical determination of multiple *O. sinensis* genotypes. According to Li et al. [36]. the primers ITS5(F) (GGAAGTAAAAGTCGTAACAAGG) and ITS4(R) (TCCTCCGCTTATTGATATGC) are fungal universal primers [92]; the primers GAF (TCCCAAACCCCCTGCGAACACC), GAR (AGGTCAACTGGAGGGTGTGGTGGTTTC) were designed specific for GC-biased Group A sequences; the primers GCF (TAGCAGTTGCCTTAGCGGGACCGCCCTA) and GCR (AATCCGAGGTTAACTAAAAGGCGTA) were designed specific for AT-biased Group B sequences; the primers 5.8S-F (ACTTTTAACAACGGATCTCTT) and 5.8S-R (AAGATAACGCTCGGATAAGCAT) were designed using the conserved 5.8S region, which should be effective for all of the paralogs. "—/—"refers to no sequence identity or sequence coverage data available when aligning the primer sequence with the sequence of *O. sinensis* genotype. Genotypes #1−3 and #7−14 are GC-biases; while Genotypes #4−6 and #15−17 are AT-biases.

**Table S7.** Summary of co-occurrence or differential occurrence of the MAT1-1-1 and MAT1-2-1 proteins in the *O. sinensis* strains, wild-type *C. sinensis* isolates, and *C. sinensis* insect–fungal complex listed in the GenBank database.

| | Number of samples | Cooccurrence of MAT1-1-1 and MAT1-2-1 proteins | Differential occurrence of mating proteins | |
|---|---|---|---|---|
| | | | MAT1-1-1 | MAT1-2-1 |
| *C. sinensis* insect–fungal complexes | 5 | 2 (40.0%) | 1 (20.0%) | 2 (40.0%) |
| Wild-type *C. sinensis* isolates | 151 | 31 (20.5%) | 85 (56.3%) | 35 (23.2%) |
| *O. sinensis* strains of different genotypes | 27 | 13 (48.1%) | 10 (37.0%) | 4 (14.8%) |
| Total | 183 | 46 (25.1%) | 96 (52.5%) | 41 (22.4%) |

### [REFERENCES]


1. Zhu, J.-S.; Halpern, G.M.; Jones, K. The scientific rediscovery of a precious ancient Chinese herbal regimen: *Cordyceps sinensis*: Part I. *J. Altern. Complem. Med.* **1998a**, *4*, 289–303. https://doi.org/10.1089/acm.1998.4.3-289.
2. Zhu, J.-S.; Halpern, G.M.; Jones, K. The scientific rediscovery of an ancient Chinese herbal medicine: *Cordyceps sinensis*: Part II. *J. Altern. Complem. Med.* **1998b**, *4*, 429–457. https://doi.org/10.1089/acm.1998.4.429.
3. Zhu, J.-S.; Li, C.-L.; Tan, N.-Z.; Berger, J.L.; Prolla, T.A. Combined use of whole-gene expression profiling technology and mouse lifespan test in anti-aging herbal product study. In Proceedings of the 2011 New TCM Products Innovation and Industrial Development Summit, Hangzhou, China, 27 November 2011; pp. 443–448. Available online: https://xueshu.baidu.com/usercenter/paper/show?paperid=08341c17fa58c8f85584b92572b90f75&site=xueshu_se (accessed on 30 January 2025).
4. Song, L.-R.; Hong, X.; Ding, X.-L.; Zang, Z.-Y. A comprehensive dictionary of modern pharmacy of Chinese traditional medicine. People's Medical Press, Beijing. 2001; pp 733-737.
5. China Ministry of Agriculture and Rural Affairs. Announcement (No. 15 of 2021) of National Forestry and Grassland Administration: List of National Key Protected Wild Plants. 7 September 2021. Available online: https://m.163.com/dy/article/HHCVOJPU055360T7.html (accessed on 3 May 2025).
6. Ren, Y.; Wan, D.-G.; Lu, X.-M.; Guo, J.-L. The study of scientific name discussion for TCM Cordyceps. *LisShenzhen Med. Mater. Medica Res.* **2013**, *24*, 2211–2212.
7. Zhang, Y.-J.; Zhang, S.; Li, Y.-L.; Ma, S.-L.; Wang, C.-S.; Xiang, M.-C.; Liu, X.; An, Z.-Q.; Xu, J.-P.; Liu, X.-Z. Phylogeography and evolution of a fungal–insect association on the Tibetan Plateau. *Mol. Ecol.* **2014**, *23*, 5337–5355. https://doi.org/10.1111/mec.12940.
8. Lu, H.-L.; St. Leger, R.J. Chapter Seven—Insect Immunity to Entomopathogenic Fungi. In *Advances in Genetics*; Lovett, B., St. Leger, R.J., Eds.; Academic Press: Cambridge, MA, USA, 2016; Volume 94, pp. 251–285.
9. Zhu, J.-S.; Li, Y.-L. A Precious Transitional Chinese Medicine, *Cordyceps sinensis*: Multiple heterogeneous *Ophiocordyceps sinensis* in the insect–fungi complex. Lambert Academic Publishing, Saarbrüchen. Germany, 2017.
10. Li, Y.-L.; Li, X.-Z.; Yao, Y.-S.; Xie, W.-D.; Zhu, J.-S. Molecular identification of *Ophiocordyceps sinensis* genotypes and the indiscriminate use of the Latin name for the multiple genotypes and the natural insect–fungi complex. *Am. J. BioMed. Sci.* **2022**, *14*, 115–135. https://doi.org/10.5099/aj220300115.
11. Li, M.-M.; Zhang, J.-H.; Qin, Q.-L.; Zhang, H.; Li, X.; Wang, H.-T.; Meng, Q. Transcriptome and Metabolome Analyses of Thitarodes xiaojinensis in Response to *Ophiocordyceps sinensis* Infection. *Microorganisms* **2023a**, *11*, 2361. https://doi.org/10.3390/microorganisms11092361.

71. Zhang, S.; Zhang, Y.-J.; Liu, X.-Z.; Wen, H.-A.; Wang, M.; Liu, D.-S. Cloning and analysis of the MAT1-2-1 gene from the traditional Chinese medicinal fungus *Ophiocordyceps sinensis*. *Fungal Biol.* **2011**, *115*, 708–714.
72. Zhang, S.; Zhang, Y.-J. Molecular evolution of three protein-coding genes in the Chinese caterpillar fungus *Ophiocordyceps sinensis*. *Microbiol. China.* **2015**, *42*, 1549–1560.
73. Kyte, J.; Doolittle, R.F. A simple method for displaying the hydropathic character of a protein. *J. Mol. Biol.* **1982**, *157*, 105–132. https://doi.org/10.1016/0022-2836(82)90515-0.
74. Deleage, G.; Roux, B. An algorithm for protein secondary structure prediction based on class prediction. *Protein Eng. Des. Sel.* **1987**, *1*, 289–294. https://doi.org/10.1093/protein/1.4.289.
75. Gasteiger, E.; Hoogland, C.; Gattiker, A.; Duvaud, S.; Wilkins, M.R.; Appel, R.D.; Bairoch, A. Protein Identification and Analysis Tools on the ExPASy Server, Chapter 52. In *The Proteomics Protocols Handbook*; Walker, J.M., Ed.; Humana Press: Totowa, NJ, USA, **2005**; pp. 571–607.
76. Peters, C.; Elofsson, A. Why is the biological hydrophobicity scale more accurate than earlier experimental hydrophobicity scales? *Proteins* **2014**, *82*, 2190–2198. https://doi.org/10.1002/prot.24582.
77. Simm, S.; Einloft, J.; Mirus, O.; Schleiff, E. 50 years of amino acid hydrophobicity scales, revisiting the capacity for peptide classification. *Biol. Res.* **2016**, *49*, 31. https://doi.org/10.1186/s40659-016-0092-5.
78. Jumper, J.; Evans, R.; Pritzel, A.; Green, T.; Figurnov, M.; Ronneberger, O.; Tunyasuvunakool, K.; Bates, R.; Žídek, A.; Potapenko, A.; Bridgland, A.; Meyer, C.: Kohl, S,A,A,; Ballard, A.J.; Cowie, A.; Romera-Paredes, B.; Nikolov, S.; Jain, R.; Adler, J.; Back, T.; Petersen, S.; Reiman, D.; Clancy, E.; Zielinski, M.; Steinegger, M.; Pacholska, M.; Berghammer, T.; Bodenstein; S.; Silver, D.; Vinyals, O.; Senior, A.W.; Kavukcuoglu, K.; Kohli, P.; Hassabis, D. Highly accurate protein structure prediction with AlphaFold. *Nature* **2021**, *596*, 583–589. https://doi.org/10.1038/s41586-021-03819-2.
79. Tunyasuvunakool, K.; Adler, J.; Wu, Z.; Green, T.; Zielinski, M.; Žídek, A.; Bridgland, A.; Cowie, A.; Meyer, C.; Laydon, A.; Velankar S, Kleywegt GJ, Bateman A, Evans R, Pritzel A, Figurnov M, Ronneberger O, Bates R, Kohl SAA, Potapenko A, Ballard AJ, Romera-Paredes B, Nikolov S, Jain R, Clancy E, Reiman D, Petersen S, Senior AW, Kavukcuoglu K, Birney E, Kohli P, Jumper J, Hassabis D. Highly accurate protein structure prediction for the human proteome. *Nature* **2021**, *596*, 590–596. https://doi.org/10.1038/s41586-021-03828-1.
80. David, A.; Islam, S.; Tankhilevich, E.; Sternberg, M.J.E. The AlphaFold Database of Protein Structures, A Biologist's Guide. *J. Mol. Biol.* **2022**, *434*, 167336. https://doi.org/10.1016/j.jmb.2021.167336.
81. Monzon, V.; Haft, D.H.; Bateman, A. Folding the unfoldable, using AlphaFold to explore spurious proteins. *Bioinform. Adv.* **2022**, *1*, vbab043. https://doi.org/10.1093/bioadv/vbab043.
82. Rettie, S.A.; Campbell, K.V.; Bera, A.K.; Kang, A.; Kozlov, S.; De La Cruz, J.; Adebomi, V.; Zhou, G.; DiMaio, F.; Ovchinnikov, S.; et al. Cyclic peptide structure prediction and design using AlphaFold. *bioRxiv* **2023**, 26:2023.02.25.529956. https://doi.org/10.1101/2023.02.25.529956.
83. Xu, T.; Xu, Q.; Li, J.-Y. Toward the appropriate interpretation of Alphafold2. *Front. Artif. Intell.* **2023b**, *6*, 1149748. https://doi.org/10.3389/frai.2023.1149748.
84. Abramson, J.; Adler, J.; Dunger, J.; Evans, R.; Green, T.; Pritzel, A.; Ronneberger, O.; Willmore, L.; Ballard, A.J.; Bambrick, J.; Bodenstein SW, Evans DA, Hung CC, O'Neill M, Reiman D, Tunyasuvunakool K, Wu Z, Žemgulytė A, Arvaniti E, Beattie C, Bertolli O, Bridgland A, Cherepanov A, Congreve M, Cowen-Rivers AI, Cowie A, Figurnov M, Fuchs FB, Gladman H, Jain R, Khan YA, Low CMR, Perlin K, Potapenko A, Savy P, Singh S, Stecula A, Thillaisundaram A, Tong C, Yakneen S, Zhong ED, Zielinski M, Žídek A, Bapst V, Kohli P, Jaderberg M, Hassabis D, Jumper JM. Accurate structure prediction of biomolecular interactions with AlphaFold 3. *Nature* **2024**, *630*, 493–500. https://doi.org/10.1038/s41586-024-07487-w.
85. Varadi, M.; Bertoni, D.; Magana, P.; Paramval, U.; Pidruchna, I.; Radhakrishnan, M.; Tsenkov, M.; Nair, S.; Mirdita, M.; Yeo, J.; et al. AlphaFold Protein Structure Database in 2024, providing structure coverage for over 214 million protein sequences. *Nucleic Acids Res.* **2024**, *52*, D368–D375. https://doi.org/10.1093/nar/gkad1011.
86. Wroblewski, K.; Kmiecik, S. Integrating AlphaFold pLDDT Scores into CABS-flex for enhanced protein flexibility simulations. *Comput. Struct. Biotechnol. J.* **2024**, *30*, 4350–4356. https://doi.org/10.1016/j.csbj.2024.11.047.
87. Li, X.-Z.; Li, Y.-L.; Zhu, J.-S. Three-dimensional structural heteromorphs of mating-type proteins in *Hirsutella sinensis* and the natural *Cordyceps sinensis* insect–fungal complex. *J. Fungi.* **2025**, *11*, 244. https://doi.org/10.3390/jof11040244.

106. Ramšak, B.; Markau, J.; Pazen, T.; Dahlmann, T.A.; Krappmann, S.; Kűck, U. The master regulator MAT1-1-1 of fungal mating binds to its targets via a conserved motif in the human pathogen *Aspergillus fumigatus*. *G3 Genes Genom. Genet.* **2020**, *11*, jkaa012. https://doi.org/10.1093/g3journal/jkaa012.
107. Yamamoto, A.; Ando, Y.; Yoshioka, K.; Saito, K.; Tanabe, T.; Shirakawa, H.; Yoshida, M. Difference in affinity for DNA between HMG proteins 1 and 2 determined by surface plasmon resonance measurements. *J. Biochem.* **1997**, *122*, 586–594. https://doi.org/10.1093/oxfordjournals.jbchem.a021793. PMID 9348088.
108. Ait Benkhali, J.; Coppin, E.; Brun, S.; Peraza-Reyes, L.; Martin, T.; Dixelius, C.; Lazar, N.; van Tilbeurgh, H.; Debuchy, R. A Network of HMG-box Transcription Factors Regulates Sexual Cycle in the Fungus Podospora anserina. *PLoS Genet.* **2013**, *9*, e1003642. https://doi.org/10.1371/journal.pgen.1003642.
109. Balasubramanian, B.; Lowry, C.V.; Zitomer, R.S. The Rox1 repressor of the *Saccharomyces cerevisiae* hypoxic genes is a specific DNA-binding protein with a high-mobility-group motif. *Mol. Cell Biol.* **1993**, *13*, 6071–6078. https://doi.org/10.1128/mcb.13.10.6071-6078.1993.
110. Zitomer, R.S.; Limbach, M.P.; Rodriguez-Torres, A.M.; Balasubramanian, B.; Deckert, J.; Snow, P.M. Approaches to the study of Rox1 repression of the hypoxic genes in the yeast Saccharomyces cerevisiae. *Methods* **1997**, *11*, 279–288. https://doi.org/10.1006/meth.1996.0422.
111. Zheng, Q.; Hou, R.; Zhang, J.-Y.; Ma, J.; Ma, J.-W.; Wu, Z.-S.; Wang, G.-H.; Wang, C.-F.; Xu, J.-R. The MAT locus genes play different roles in sexual reproduction and pathogenesis in *Fusarium graminearum*. *PLoS ONE.* **2013b**, *8*, e66980. https://doi.org/10.1371/journal.pone.0066980.
112. Kastaniotis, A.J.; Zitomer, R.S. Oxygen Dependent Repression in Yeast. In *Rox1 Mediated Repression*; Advances in Experimental Medicine and Biology; Springer Nature: Cham, Switzerland, 2000; Volume 475, pp. 185–195. https://doi.org/10.1007/0-306-46825-5_18.
113. Kües, U.; Casselton, L.A. The origin of multiple mating types in mushrooms. *J. Cell Sci.* **1993**, *104*, 227–230. https://doi.org/10.1242/jcs.104.2.227.
114. Asante-Owusu, R.N.; Banham, A.H.; Böhnert, H.U.; Mellor, E.J.C.; Casselton, L.A. Heterodimerization between two classes of homeodomain proteins in the mushroom *Coprinus cinereus* brings together potential DNA-binding and activation domains. *Gene* **1996**, *172*, 25–31. https://doi.org/10.1016/0378-1119(96)00177-1.
115. Jacobsen, S.; Wittig, M.; Pöggeler, S. Interaction Between Mating-Type Proteins from the Homothallic Fungus Sordaria macrospora. *Curr. Genet.* **2002**, *41*, 150–158. https://doi.org/10.1007/s00294-002-0276-0.
116. Hancock, S.P.; Cascio, D.; Johnson, R.C. Cooperative DNA binding by proteins through DNA shape complementarity. *Nucleic Acids Res.* **2019**, *47*, 8874–8887. https://doi.org/10.1093/nar/gkz642.
117. Mao, X.-M.; Zhao, S.-M.; Cao, L.; Yan, X.; Han, R.-C. The morphology observation of *Ophiocordyceps sinensis* from different origins. *J. Environ. Entomol.* **2013**, *35*, 343–353.
118. Li, Y.; Yang, R.-H.; Jiang, L.; Hu, X.-D.; Wu, Z.-J.; Yao, Y.-J. rRNA Pseudogenes in Filamentous Ascomycetes as Revealed by Genome Data. *G3-Genes Genom. Genet.* **2017**, *7*, 2695–2703. https://doi.org/10.1534/g3.117.044016.
119. Li, Y.; Jiang, L.; Wang, K.; Wu, H.-J.; Yang, R.-H.; Yan, Y.-J.; Bushley, K.E.; Hawksworth, D.L.; Wu, Z.-J.; Yao, Y.-J. RIP mutated ITS genes in populations of *Ophiocordyceps sinensis* and their implications for molecular systematics. *IMA Fungus* **2020c**, *11*, 18.
120. Li, X.-Z.; Li, Y.-L.; Wang, Y.-N.; Zhu, J.-S. Translations of mutant repetitive genomic sequences in *Hirsutella sinensis* and changes in secondary structures and functional specifications of the encoded proteins. *Int. J. Mol. Sci.* **2024a**, *25*, 11178. https://doi.org/10.3390/ijms252011178.
121. Wang, Y.-B., Wang, Y., Fan, Q., Duan, D-E., Zhang, G.-D., Dai6, R.-Q., Dai, Y.-D., Zeng, W.-B. Chen, Z.-H., Li, D.-D., Tang, D.-X., Xu, Z.-H., Sun, T., Nguyen, T.-T., Tran, N.-L., Dao, V.-M., Zhang, C.-M., Huang, L.-D., Liu, Y.-J., Zhang, X.-M., Yang, D.-R., Sanjuan, T., Liu, X.-Z., Yang, Z.-L., Yu, H. Multigene phylogeny of the family Cordycipitaceae (Hypocreales): new taxa and the new systematic position of the Chinese cordycipitoid fungus *Paecilomyces hepiali*. *Fungal Diversity* **2020**, *103*, 1–46. https://doi.org/10.1007/s13225-020-00457-3.
122. Holliday, J.; Cleaver, M. Medicinal value of the caterpillar fungi species of the genus *Cordyceps* (Fr.) Link (Ascomycetes). A review. *Int. J. Med. Mushrooms* **2008**, *10*, 219–234. https://doi.org/10.1615/IntJMedMushr.v10.i3.30.
123. Stone, R. Improbable partners aim to bring biotechnology to a Himalayan kingdom. *Science* **2010**, *327*, 940–941. https://doi.org/10.1126/science.327.5968.940.
